# Robust Random Forests for Genomic Prediction: Challenges and Remedies

**DOI:** 10.64898/2026.03.30.715203

**Authors:** Vanda M. Lourenço, Joseph O. Ogutu, Hans-Peter Piepho

## Abstract

Data contamination—from recording errors to extreme outliers—can compromise statistical models by biasing predictions, inflating prediction errors, and, in severe cases, destabilizing performance in high-dimensional settings. Although contamination can affect responses and covariates, we focus on response contamination and evaluate Random Forests through simulation. Using a synthetic animal-breeding dataset, we assess robust Random Forests across several contamination scenarios and validate them on plant and animal datasets. We thereby clarify the consequences of contamination for prediction, develop a robust Random Forest framework, and evaluate its performance. We examine preprocessing or data-transformation strategies, algorithmic modifications, and hybrid approaches for robustifying Random Forests. Across these approaches, data transformation emerges as the most effective strategy, delivering the strongest performance under contamination. This strategy is simple, general, and transferable to other Machine Learning methods, offering a remedy for robust genomic prediction. In real breeding data, robust Random Forests are useful when substantial contamination, phenotypic corruption, misrecording, or train–deployment mismatch is plausible and the goal is to recover a latent signal for genomic prediction and selection; in that setting, ranking-based and weighting-based robust Random Forests offer complementary remedies, the latter through a median-centred formulation that gives shrinkage a coherent interpretation across response distributions. Robustification is not universally necessary, but it becomes important when contamination distorts the link between observed responses and the predictive target; standard Random Forests remain the default for clean data, whereas robust Random Forests should be fitted alongside them whenever contamination is plausible, with the final choice guided by data, trait, and breeding objective.

**Author summary:** Machine learning (ML) methods are widely used for prediction with high-dimensional, complex data, and supervised approaches such as Random Forests (RF) have proved effective for genomic prediction (GP) and selection. Yet their performance can be severely compromised by data contamination if the algorithms rely on classical data-driven procedures that are sensitive to atypical observations. Robustifying ML methods is therefore important both for improving predictive performance under contamination and for guiding their practical use in high-dimensional prediction problems. To address this need, we develop robust preprocessing, algorithm-level, and hybrid strategies for improving RF performance with contaminated data. Using simulated animal data, we show that ranking- and weighting-based robust RF provide the strongest overall compromise for genomic prediction and selection under contamination. Validation on several plant and animal breeding datasets further shows that the benefits of robustification are not universal, but depend on the dataset, trait, and breeding objective. Although motivated by RF, the framework we propose is general, practical, and readily transferable to other ML methods. It also offers a basis for deciding when robustness should complement standard RF rather than replace it outright.

## Introduction

The accelerating availability of high-dimensional data is transforming prediction across disciplines, making machine-learning methods central to modern inference and decision-making. In plant, animal, and human genomics, a flagship application is genomic prediction (GP): forecasting complex quantitative phenotypes—such as yield, height, flowering time, blood pressure—or breeding values from genome-wide markers [1, 2, 3, 4, 5]. GP typically uses thousands of molecular markers (usually single nucleotide polymorphisms - SNPs) distributed across the genome, a scale that demands methods that remain accurate, stable, and computationally efficient in high dimensions. Consequently, machine-learning (ML) approaches have gained prominence in GP because they flexibly represent complex, potentially nonlinear marker–trait relationships while retaining practical computational performance [6, 7, 8, 9, 10, 11, 12, 13, 14].

Yet flexibility does not guarantee reliability. Like classical methods, ML models can degrade sharply when phenotypic data violate common distributional assumptions—especially through contamination and outliers—undermining both predictive accuracy (PA) and precision [15, 16, 17, 18, 19, 20, 21, 22, 23]. Motivated by the combined effects of many small-effect loci and environmental influences under the central limit theorem, approximate normality is often invoked for continuous quantitative traits. In practice, however, phenotypes are often contaminated by unobserved biological or environmental influences that generate a small fraction of observations deviating from the bulk distribution, thereby driving marked departures from normality. These model-relative outliers are not always visually extreme (and may therefore appear as inliers) and can reflect localized environmental effects or biological stressors in plant and animal studies (e.g., specific years, field pests, management conditions), cohort effects or unobserved heterogeneity in human studies, or other latent sources of variability. Although excluding such observations is often advocated [24, 25, 26, 27], reliable outlier detection remains inherently challenging, especially under masking and swamping effects [28]. Importantly, outliers can be scientifically meaningful, particularly in breeding contexts, making it undesirable to discard them; what is needed, instead, are modeling approaches that limit their undue influence while retaining the information they carry [29]. Conceptually distinct—but often conflated—are data collection or recording errors, which arise from data-acquisition failures and are typically addressed through data curation.

This vulnerability motivates a central challenge in GP: robustifying high-dimensional prediction, including ML-based prediction. Even when ML methods do not posit an explicit distribution for the response, their training and stability can remain sensitive to the empirical phenotype distribution; departures from normality can destabilize fitting, distort split criteria, and erode PA [30]. Despite this practical reality, robust variants of ML methods used for high-dimensional prediction in GP remain scarce. Nevertheless, robust statistical approaches have been explored in related problems in genetic studies, including robust variance-component estimation and heritability assessment, as well as the evaluation of PA under contamination [31, 32].

Here we address this gap by developing robust random forests (RF) for GP. We propose and evaluate four complementary strategies to robustify RF for GP—strategies designed to reduce sensitivity to non-normality and phenotype contamination and, in turn, improve predictive performance under realistic data conditions. The strategies include: (1) preprocessing-based remedies, applying data transformations before fitting RF; (2) algorithm-based remedies, modifying RF to enhance robustness; (3) hybrid remedies, integrating the best-performing preprocessing and algorithmic modifications; and (4) comparative evaluation, benchmarking these approaches across multiple plausible contamination scenarios. We assess performance using a simulated animal-breeding data set and then validate the most effective remedies on several empirical plant and animal data sets. Across these four strategies, although robustness is evaluated at both the preprocessing and algorithmic levels, the dominant gains in predictive performance are consistently driven by preprocessing—specifically, data transformation—thereby yielding a winning strategy that is not unique to RF and is readily transferable to other machine-learning approaches.

### Animal data (simulated)

An outbred animal-breeding population dataset, simulated for the 16−th QTLMAS Workshop 2012, underpins our evaluation of contamination effects in GP. The simulation models used to generate the data are described in detail in [33] and are therefore not reproduced here. Although they do not provide the precise mathematical details of the data-generating model, the authors report that the three simulated traits (T1, T2, and T3) represent quantitative milk traits (*milk yield, fat yield & fat content*). They further note that the true breeding values (TBV) for the three traits were computed as the sum of additive effects from 50 QTLs. Random residuals were sampled from normal distributions with mean 0 and trait-specific residual variances, yielding heritabilities of 0.35, 0.35, and 0.50 for T1, T2, and T3, respectively. Mean-centering therefore produces some negative simulated phenotypic values. The resulting dataset comprises 4020 individuals genotyped at 9969 SNP markers; 3000 individuals phenotyped for the three quantitative milk traits, whereas 1020 were not phenotyped. Table 1 summarizes trait 1 (T1), the focus of our simulations, and Figure S1 complements these summaries with graphical distributional diagnostics.

**Table 1.** Summary statistics for the selected animal quantitative trait dataset (trait *T*_1_). Training data contain *n* = 3000 observations and testing data contain *n* = 1020 observations.

|  | Trait Dataset | Min. | 1st Qu. | Median | Mean | 3rd Qu. | Max. | Sd |
| --- | --- | --- | --- | --- | --- | --- | --- | --- |
| $T_1$ | Training | -584.99 | -116.24 | -1.71 | 0.00 | 112.25 | 587.19 | 176.52 |
|  | Testing | -325.04 | -59.65 | 16.88 | 14.70 | 87.98 | 356.29 | 107.71 |

For the *T*_1_ training data, the Normal Q–Q plot indicates close agreement with normality, consistent with the Shapiro–Francia test (*p* − *value* ≃ 0.053), a large-sample approximation to the Shapiro–Wilk test, which shows no evidence of departure from normality at the 5% level. The violin plot indicates an approximately symmetric distribution with slightly long tails, and the boxplot highlights a small number of observations at both extremes as potential outliers; collectively, these features do not materially compromise approximate normality. The corresponding distribution diagnostics for the *T*_1_ testing data (Figure S1, 1st row), display a similar pattern and are likewise consistent with approximate normality (Shapiro–Francia *p* − *value* ≃ 0.557). Summary statistics and distribution diagnostics for both the training and testing data across the three traits are reported in Table S1 and Figure 1S, respectively, completing the characterization of the simulated phenotypic data.

The simulated dataset also provides true genomic breeding values (TGBVs) for the 1020 genotypes for all traits; full details are reported in [14]. Using genomic information and focusing on T1, we predict genomic breeding values (PGBVs) for the 1020 unphenotyped individuals under uncontaminated conditions and use these predictions as a benchmark against which to quantify the impact of contamination across scenarios—a reference standard that anchors the simulated animal analysis.

### Real plant and animal data

Real plant and animal datasets provide a rigorous empirical testbed for comparing the performance of standard and robust RF in GP. We evaluate the models using four example real datasets—three plant and one animal—to assess their relative performance under contrasting biological and genomic settings. For the plant data, we use: (i) the maize and soybean data of [34], obtained from [35] (datasets 00021 & 00023); and (ii) the wheat data made publicly available by [36] and acessible through the R package BGLR [37]. For the animal data we use the mice dataset from [38], available through the R package BLR. These datasets jointly furnish a demanding real-data benchmark, one that tests model performance not in abstraction but in the empirical complexity of biological data.

To ensure analytical consistency, we restricted the analyses to quantitative traits with complete observations and characterized their distributional properties in detail. For the genomic data, markers with substantial missingness were assessed and, when necessary, filtered using a 20% missing-value threshold, whereas datasets with negligible missingness required no additional preprocessing.

The maize dataset comprises 278 accessions obtained from the Panzea database and genotyped for 50, 896 SNPs, with phenotypic records for two quantitative traits, *days to tasseling* and *ear height*. For the analysis only the complete trait *ear height* is used; its broad-sense heritability is 0.65. The SNP data were generated using the Illumina MaizeSNP50 Bead-Chip [39]. In addition, SNPs with *>* 20% missing values were removed before analysis, so only the 47, 259 SNPs remaining after filtering are used in the analyses.

The soybean dataset comprises 346 accessions genotyped for 36, 901 SNPs, with SNP data generated using the Illumina Infinium SoySNP50K iSelect SNP BeadChip [40, 41]. For the analysis, two complete quantitative traits are used, namely *carbon isotope ratio* and *canopy wilting*. In addition, SNPs with *>* 20% missing values were removed before analysis, so only the 31, 260 SNPs remaining after filtering are used in the analyses.

The wheat dataset captures substantial agronomic breadth and thereby strengthens the plant-data benchmark. It consists of 599 historical CIMMYT wheat lines derived from CIMMYT’s Global Wheat Program, which has long conducted international trials across a broad range of wheat-producing environments and thus generated material of wide agronomic relevance. These trial environments were classified into four target sets representing four major agroclimatic regions, a grouping previously defined and widely used within CIMMYT’s Global Wheat Breeding Program. The phenotypic trait analyzed here is average *grain yield* (GY) for the 599 lines evaluated within each of these four mega-environments. Because the average yield values for the 599 genotypes were standardized within each environment to zero mean and unit variance, we first derived genotype-level responses across the four environments before analysis. Specifically, we estimated adjusted means for each genotype using a general linear model with standardized yield as the response and genotype and environment as fixed effects. Because each genotype occurred once in each environment, the design was balanced, and these adjusted means were therefore equivalent to the arithmetic means across the four environments. We used these adjusted means as the response in the standard RF models. The 599 wheat lines were genotyped for 1279 SNP markers. By linking broad environmental representation with a well-characterized yield trait, this dataset adds both depth and rigour to the comparative evaluation. All trait-by-environment data are complete and are therefore retained for analysis. The SNP data contained no SNPs with *>* 20% missing values, and all SNPs are therefore retained for analysis.

The animal dataset extends this benchmark by adding a distinct species, a larger sample size, and a broader trait spectrum. For the animal data, we use the mice dataset from [38], available through the R package BLR, comprising 1814 mice genotyped for 10,346 SNPs and phenotyped for 21 quantitative traits, including 3 obesity-related traits and 18 biochemical traits. For the mice data, we analyze only three complete quantitative obesity traits: *body mass index, body length*, and *final normalized body weight*. The SNP data contained no SNPs with *>* 20% missing values, and all SNPs are therefore retained for analysis.

In combination, the plant and animal datasets span distinct species, numbers of genotypes, marker densities, and trait architectures, thereby providing a stringent real-data benchmark for evaluating standard versus robust RF for GP. This breadth of biological context makes the comparison sharper, more rigorous, and more compelling.

Summary statistics for all selected traits, together with the p-values of the Shapiro–Francia normality test, are reported in Table S2, whereas the distribution diagnostic plots—histograms, Q–Q plots, and violin plots—for all traits are presented in Figures S2–S5. These summaries and diagnostics provide the empirical basis for interpreting model performance across traits that differ not only in biological meaning but also in statistical behaviour. The maize, soybean, wheat, and mice datasets anchor the analysis in real biological complexity and provide a robust platform for judging when standard methods suffice, when robust methods prevail, and how model performance shifts across the varied terrain of real plant and animal data.

### Data contamination

In this work, contamination refers broadly to any mechanism that perturbs the data-generating process, including recording errors, discretization artifacts, placeholder values, and unobserved biological or environmental influences. To model the subset of contamination that generates distributional deviations, we adopt Huber’s *ε*-contamination framework, in which a small fraction of observations is drawn from a contaminating distribution G. In our setting, these observations constitute the model-relative outliers. Although the literature recognizes alternative definitions of outliers, including distance-based, density-based, and residual-based approaches [42, 43, 44, 45], we adopt the model-based perspective induced by the contamination framework. Thus, while contamination spans a wide range of real-world processes, only the subset represented by the G-component is treated as outlying in our model.

We empirically evaluate the comparative predictive performance of standard and robustified RF methods on the uncontaminated and contaminated simulated animal dataset. Contaminated responses are generated under Huber’s contamination mixture model (1 − *ε*)*F* + *εG*, where *F* is the original data distribution, *G* is a contaminant distribution, and *ε* is the proportion of contaminating observations [46, 47]. Because the baseline responses are approximately *N*(*µ, σ*^2^) with *µ* and *σ*^2^ estimated from the data, contamination is implemented by replacing a proportion *ε* of observations with draws from *G*. We consider *ε* = 2%, 5% & 10% and four contamination types: (i) `shift` contamination, with outliers drawn from *N*(*µ* + *kσ, σ*^2^); (ii) `variance-inflated` contamination, with outliers drawn from *N*(*µ*, (*sσ*)^2^); (iii) `central variance-deflated` (pointmass type) contamination, with outliers drawn from *N*(*µ*, (*σ/γ*)^2^); and (iv) `tail variance-deflated` contamination, corresponding to a mixture of scenarios (i) and (iii) with *γ* = 10000 only. Here, *µ* denotes the trait sample mean and *σ*^2^ the trait sample variance, with *k* = 5, 7, 9, *s* = 5, 7 and *γ* = 1000, 10000. The resulting contaminated-response distributions are shown in Figure S6, together with Shapiro–Francia p-values for normality. The corresponding mixture-model representations are illustrated in Figure 1, and the practical interpretations of the contamination mechanisms are summarized in Table 2. `The central` and `tail variance-deflated` contamination scenarios are not intended to reproduce the exact mechanisms underlying missing-value imputation, detection limits, rounding, or placeholder values. Instead, they mimic the distributional consequences of such processes by concentrating contaminated observations either near the mean (central) or at a single extreme value (tail). In this way, the contaminating distribution *G* serves as a stylized, model-based representation of the distortions these procedures can induce. To distinguish contamination effects from randomness inherent in RF, we used, across all scenarios, the same 10 contaminated datasets generated from the same simulated animal dataset for each contamination scenario.

**Fig. 1.**
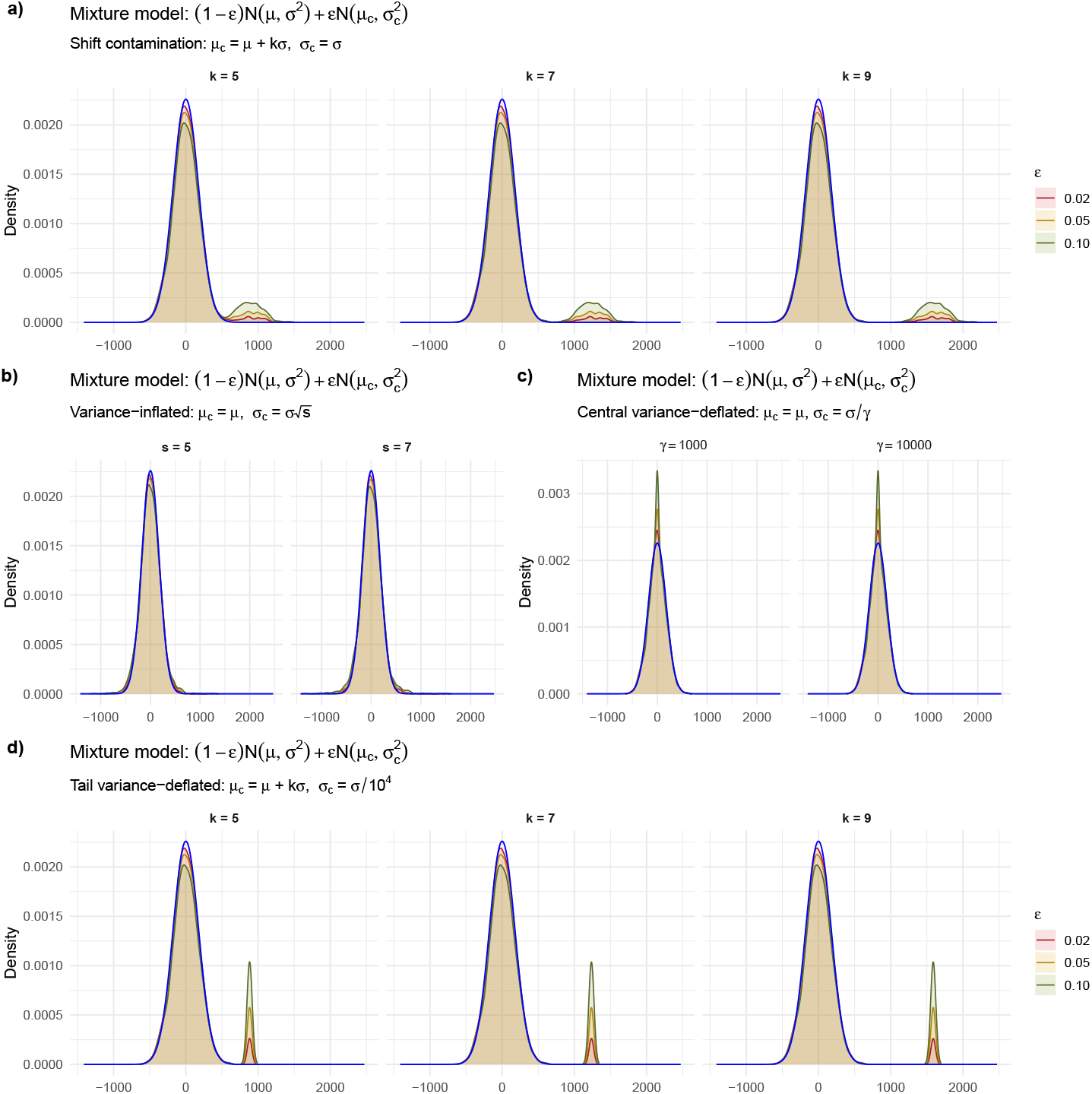
Overlay plots for the Huber mixture model under `shift, variance-inflated`, and `variance-deflated` contaminations. The baseline *N*(*µ, σ*^2^) distribution is shown in blue, and mixtures are shown as shaded densities for contamination levels *ε* ∈ {0.02, 0.05, 0.10}.

**Table 2.**
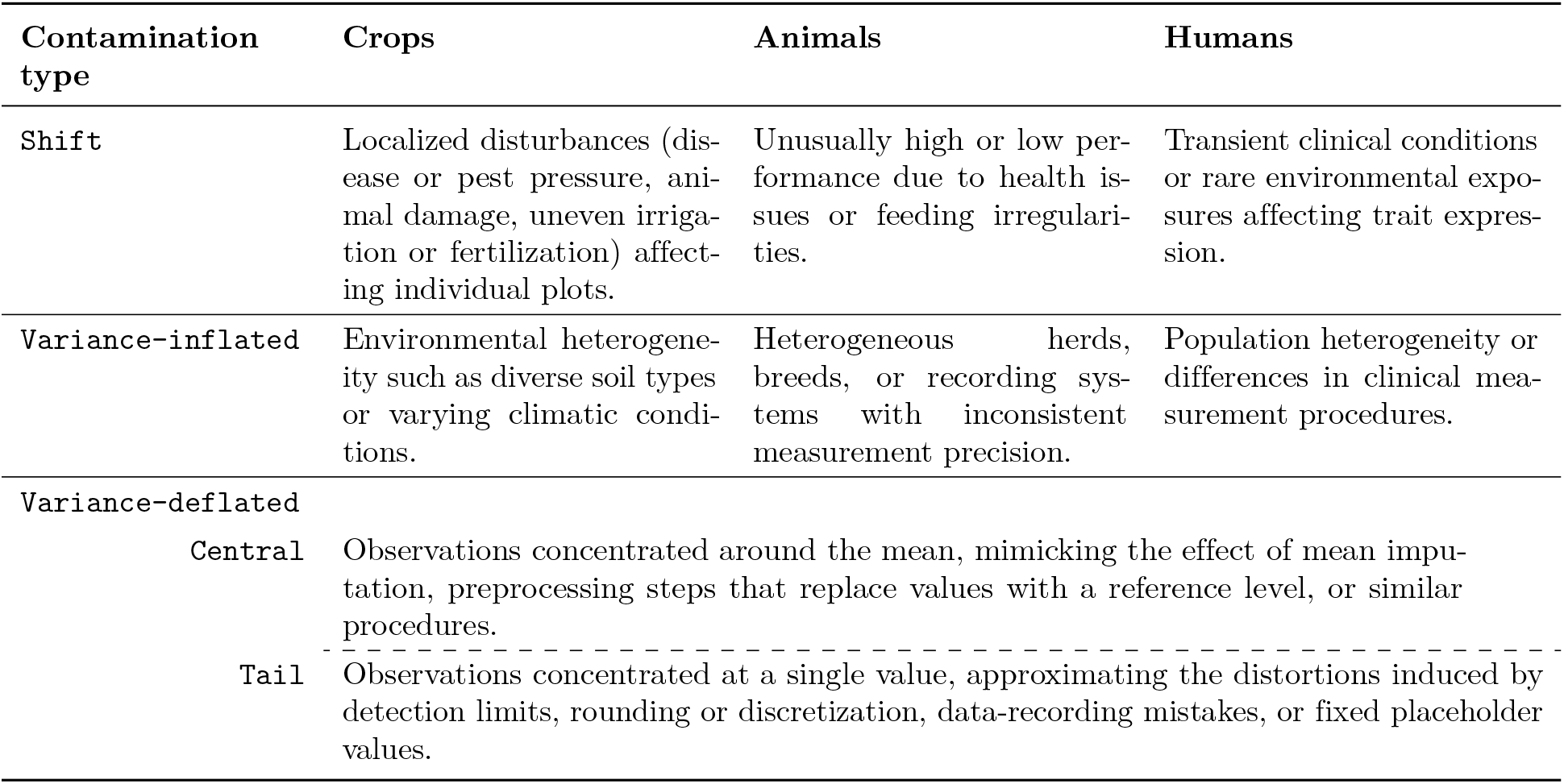
Example contamination generating mechanisms across contrasting application domains.

| Contamination type | Crops | Animals | Humans |
| --- | --- | --- | --- |
| Shift | Localized disturbances (disease or pest pressure, animal damage, uneven irrigation or fertilization) affecting individual plots. | Unusually high or low performance due to health issues or feeding irregularities. | Transient clinical conditions or rare environmental exposures affecting trait expression. |
| Variance-inflated | Environmental heterogeneity such as diverse soil types or varying climatic conditions. | Heterogeneous herds, breeds, or recording systems with inconsistent measurement precision. | Population heterogeneity or differences in clinical measurement procedures. |
| Variance-deflated |  |  |  |
| Central | Observations concentrated around the mean, mimicking the effect of mean imputation, preprocessing steps that replace values with a reference level, or similar procedures. |  |  |
| Tail | Observations concentrated at a single value, approximating the distortions induced by detection limits, rounding or discretization, data-recording mistakes, or fixed placeholder values. |  |  |

## Methods

### Random Forests for Regression

Random forests [48] are tree-based ensemble algorithms for classification and regression and are well suited for high-dimensional problems, including SNP-based predictions. A schematic representation of the RF algorithm is shown in Figure 2. RF draws *N* bootstrap samples from the original dataset; each sample is used to grow one tree in the ensemble. Each tree is constructed by recursively partitioning its bootstrap sample using a splitting rule until a stopping criterion is satisfied. At every split, only a random subset of features (predictors) of size *m* is considered, introducing diversity across trees. Split selection is driven by a node-impurity criterion defined through a loss function, and predictions from the *N* trees are aggregated to yield the final RF predictor.

**Fig. 2.**
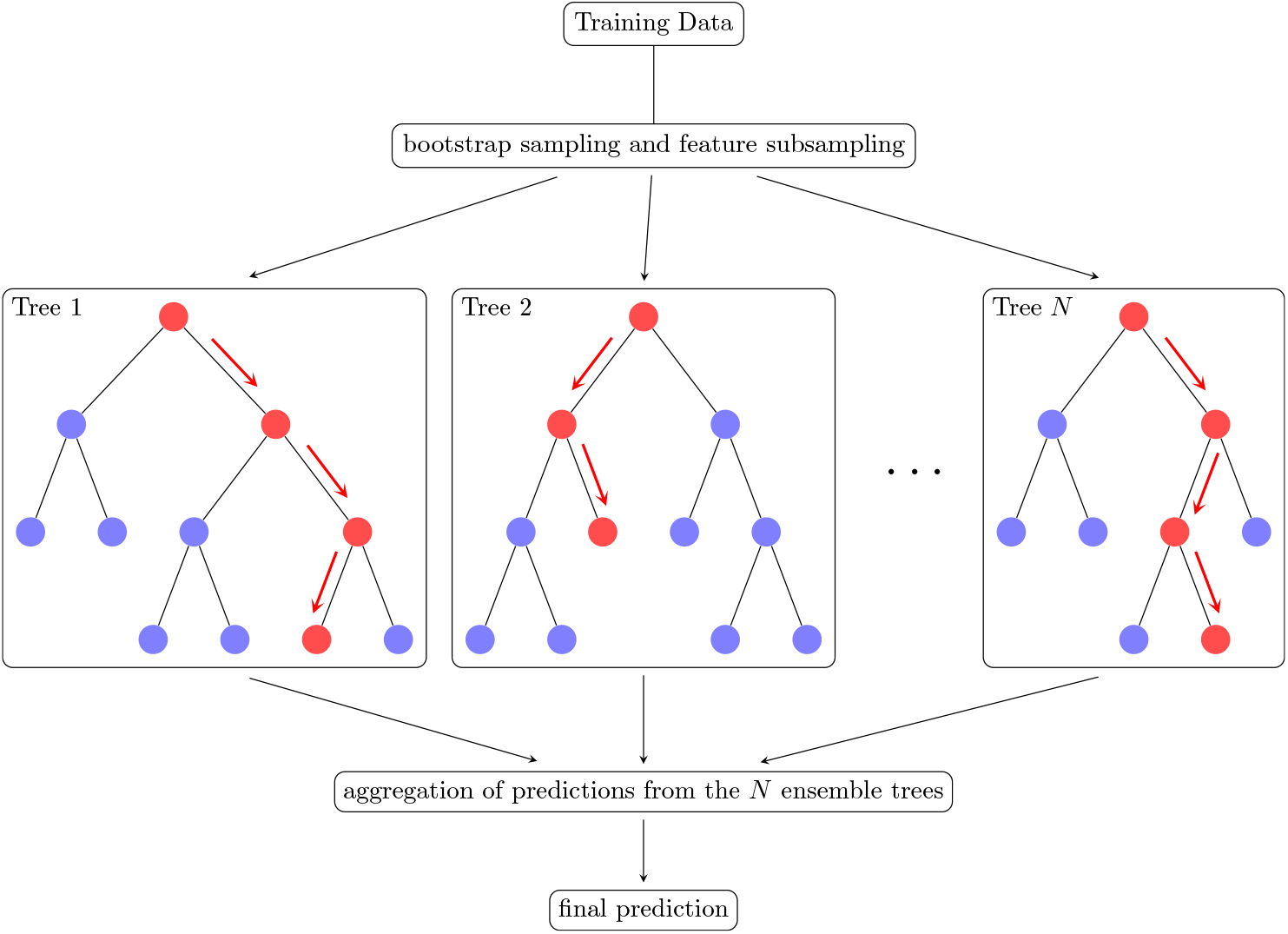
Schematic of random forests (RF) with *N* decision trees, showing split nodes and the aggregation of tree predictions into the final output. Red arrows indicate the path of a sample through each tree during prediction.

For regression, node impurity at node *t* is measured by squared-error loss, namely the mean squared error (MSE),

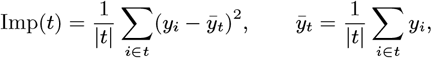

where |*t*| is the number of observations in node *t*.

Among all candidate splits, RF selects the feature and split point minimizing the weighted sum of child-node impurities,

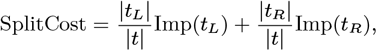

with |*t*_*L*_| and |*t*_*R*_| denoting the numbers of observations in the left and right child nodes. When predictors correspond to SNPs coded as 0*/*1*/*2, only the splits *x* ≤ 0 and *x* ≤ 1 are evaluated, and the split minimizing the weighted child-node impurity is selected. Tree growth proceeds recursively until a stopping criterion is met—commonly a minimum terminal-node size, a maximum depth, or the absence of further impurity reduction—after which the forest aggregates tree-level predictions; for regression, aggregation is the mean across the *N* trees.

RF performance is shaped by hyperparameters that govern ensemble complexity and tree diversity. The number of trees controls a stability–cost trade-off: more trees typically stabilize predictions but increase computational time. The number of candidate features *m* at each split regulates inter-tree correlation and is therefore central to tuning: overly large *m* yields similar trees and limited variance reduction, whereas overly small *m* can undersample informative predictors and degrade accuracy. The minimum number of observations in a terminal (leaf) node controls tree depth and the bias–variance trade-off: node size one produces fully grown trees, whereas larger values constrain depth, increasing bias while reducing variance. Together, these tuning parameters modulate how RF explores high-dimensional predictor spaces and how reliably it generalizes.

Despite strong empirical performance, standard RF for regression is not robust to response contamination. Outliers render the mean—a core building block of both splitting and aggregation—fragile [49, 50]. The mean-based impurity criterion (MSE) is highly sensitive to outlying observations, which can distort split decisions and propagate error through the recursive partitioning, producing poorly constructed trees. Mean aggregation across trees can then transmit these distortions to the ensemble level, yielding biased predictions. The bootstrap itself is also non-robust [51, 52, 53]: bootstrap samples that over-represent contaminated observations can induce biased trees and unreliable splits. Finally, performance assessment based on out-of-bag (OOB) samples, subset of the data consisting of observations not included in the bootstrap sample used to construct a tree, may be misleading under contamination because OOB sets may include outliers, compromising the resulting error estimates.

Collectively—mean-based splitting, mean-based aggregation, non-robust resampling, and contaminated OOB evaluation—these mechanisms explain why robust RF variants are needed to preserve predictive reliability when data depart from idealized assumptions, especially in practical genomic settings where contamination is plausible.

### Robust Strategies

The lack of robustness of standard RF can be traced to several algorithmic components—mean-based splitting criteria, aggregation by averaging, and non-robust resampling and performance assessment—each of which can amplify the influence of outliers. Two complementary strategy classes mitigate these effects: preprocessing-based remedies that aim to robustify the data before fitting RF, and algorithm-based remedies that modify RF itself. Preprocessing strategies leave the RF algorithm unchanged but reduce contamination influence by transforming the inputs—most commonly the response—providing a natural, computationally simple pathway to robustness that can be deployed without redesigning the learning procedure. Algorithmic strategies, in contrast, introduce robustness directly into RF by altering resampling schemes, impurity measures or aggregation rules; several such methods exist, although not in the context of GP studies, and are reviewed in the following sections. In what follows, we examine six preprocessing-based approaches and four algorithm-based approaches, thereby isolating where robustness can be gained most simply—at the data interface of RF.

#### Preprocessing-Based Approaches

When quantitative responses depart from normality or contain outliers, a common remedy is to transform the response prior to model fitting. Below we detail six preprocessing strategies.

1. Box-Cox [54, 55, 56, 57] & robust Box-Cox transformations [58]

For a positive response *y >* 0, the (normalized) Box-Cox transformation is

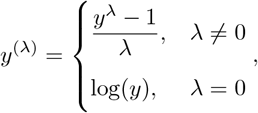

with *λ* typically estimated by maximizing the Gaussian log-likelihood of the transformed response under normality and constant variance.

For independent observations *y*_1_, …, *y*_*n*_, this log-likelihood is

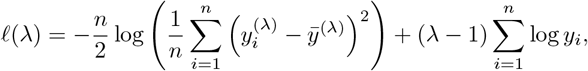

where 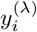 is the Box–Cox transformation of *y*_*i*_ and 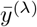 its sample mean. Common values include *λ* = 1 (no transformation; just a unit shift), *λ* = 0 (log-transform), *λ* = 0.5 (square root).

After transformation we denote the transformed variable by *w* = *y*^(*λ*)^; modeling and prediction are performed on *w*, and predictions *ŵ* are back-transformed via

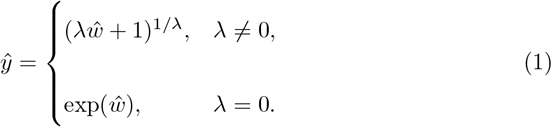

Because this inverse Box–Cox transformation does not generally preserve the conditional mean on the original scale, direct back-transformation can yield biased predictions [59, 60]. To reduce this retransformation bias, we apply a robust location correction,

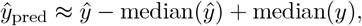

which aligns predicted and observed responses through a robust measure of location. This adjustment preserves the overall location of the response while remaining resistant to contamination and asymmetry. It is best viewed as a robust centering correction rather than as a full mean-recovery procedure, such as the smearing estimator of [59].

When *y* contains zeros or negatives, we first shift by a small constant,

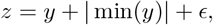

with *ϵ >* 0 (we set *ϵ* = 10^−6^), apply Box–Cox to *z*, model and predict on *w*, back-transform predictions *ŵ* to 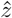 using (1), subtract | min(*y*)| + *ϵ*, and compute

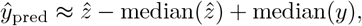

approximately restoring the original scale while preserving central tendency. This median-based correction does not affect rank-correlation metrics such as PA or predictive ability (PAb), but it can improve error-based measures such as MSE or the mean absolute error (MAE) by aligning predictions with the original response scale.

To reduce outlier influence on *λ*, we use [58] robust Box–Cox targeting central normality: the transformed data are approximately Gaussian in the center while allowing a small set of observations to remain outlying. Rather than likelihood maximization, *λ* is estimated by a two-stage procedure: an initial bounded fit comparing transformed values to normal quantiles, followed by a reweighted likelihood refinement based on inliers. With *y*_1_, …, *y*_*n*_ data are robustly standardized by

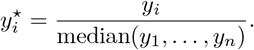

For candidate *λ*, define 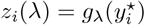, where *g*_*λ*_(·) denotes the Box–Cox transform above. The robust estimate 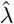 minimizes a bounded objective,

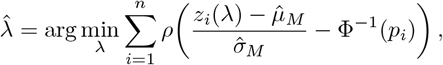

where 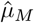 and 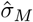 are Huber’s robust location and scale estimators, Φ^−1^ is the standard normal quantile function, *p*_*i*_ = (*i* − 1*/*3)*/*(*n* + 1*/*3), and *ρ*(·) is Tukey’s bisquare loss.

After transformation 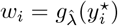 and prediction *ŵ*, back-transformation proceeds by (i) inverse Box–Cox,

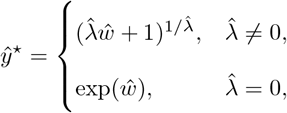

(ii) restoring scale *ŷ* = median(*y*) *ŷ*^⋆^, and (iii) optional robust location correction

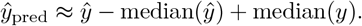

Yeo-Johnson [61] & robust Yeo-Johnson transformations [58]

The Yeo–Johnson transformation extends the Box–Cox transformation to *y* ∈ ℝ:

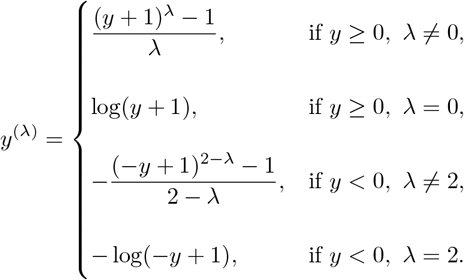

With *w* = *y*^(*λ*)^, models are fit to *w* and predictions are back-transformed using the inverse Yeo–Johnson map:

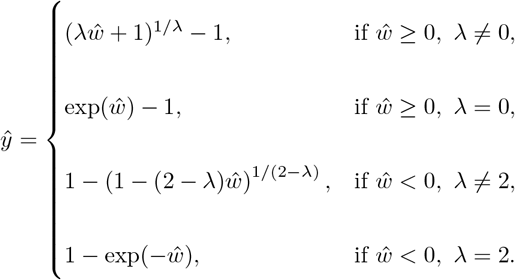

As for Box–Cox, *λ* is often estimated by Gaussian profile likelihood and can be outlier-sensitive; [58] robustify estimation by (i) robust standardization and (ii) minimizing a bounded normal-quantile objective, followed by a reweighted maximum-likelihood refinement. As with Box–Cox, to correct residual location distortions—especially under contamination or robust fitting—we apply

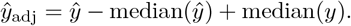

2. Winsorization [62, 63, 64]

Winsorization reduces outlier influence by replacing extreme values with predetermined quantiles. For trimming proportion *α*, the smallest *α*-fraction is set to the *α*-th percentile and the largest *α*-fraction to the (1 − *α*)-th percentile. This attenuates tail leverage while preserving the rank structure and original units. However, replacing values by fixed quantiles can introduce artificial plateaus (spikes) at truncation points, concentrating probability mass at the boundaries and disrupting smoothness, continuity, and symmetry. Because winsorization is not invertible, models fit to winsorized responses yield predictions interpreted directly on the winsorized scale.

4. Rank transformation [65, 66]

Rank transformation replaces each response by its rank, reducing sensitivity to heavy tails and outliers. For *y* = (*y*_1_, …, *y*_*n*_),

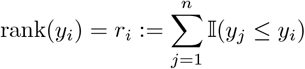

With ties, average ranks are assigned: for 𝒯 = {*k* : *y*_*k*_ = *y*_*i*_},

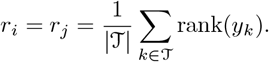

The transformed response lies in [1, *n*] (or [1, *n*^∗^] when ties yield *n*^∗^ *< n*). Predictions from models trained on ranks are typically non-integers on a pseudo-rank scale; to recover original-scale predictions we linearly interpolate between order statistics.

Let *y*_(1)_ ≤ *y*_(2)_ ≤ · · · ≤ *y*_(*n*_∗_)_ be ordered distinct training responses. For predicted rank 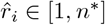,

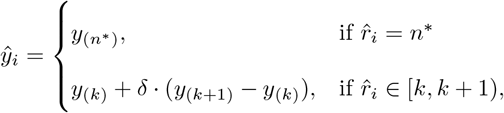

with 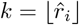 and 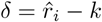.

This differs from [20], who round predicted ranks to the nearest integer and map them to observed training values; our interpolation treats 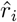 as continuous, yielding smoother predictions not restricted to observed responses while preserving ordinal information. Nonetheless, back-transformation can reintroduce extreme-value influence under contamination, inflating error-based metrics even when rank correlation remains high; we therefore apply the robust location correction

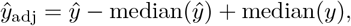

to align centers while preserving ranking.

5. Exploratory: Median Winsorization

Beyond classical winsorization, we consider a variant that replaces extreme values in both tails with the sample median. Unlike standard winsorization—which truncates to tail quantiles (e.g., 5th and 95th percentiles)—median winsorization collapses tail extremes toward the center to further dampen outlier influence. This strategy is not designed to approximate normality and can distort the distribution, especially under asymmetric contamination: concentrating probability mass near the center can inflate the peak and reduce variability, potentially exacerbating departures from normality.

6. Exploratory: Robust Weighting

Robust weighting offers a targeted preprocessing remedy when outlying responses threaten to obscure the predictive signal. We implement this approach by first fitting a robust location-only model to the response by M-estimation, which yields observation-level robustness weights ***ω***_*i*_ ∈ [0, 1], from the Huber influence function applied to standardized residuals. The M-estimation procedure uses the Huber loss with tuning constant *c* = 1.345; thus, large-residual observations are down-weighted, whereas central observations retain full weight. A direct weighted response can be defined as

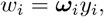

and the random forest can then be fitted with *w*_*i*_ as the response, so predictions are interpreted on the weighted-response scale. Under this direct formulation, observations in the tails of the residual distribution are pulled towards zero, while observations in the central region retain weights close to one [49, 50]. This zero-centred pull, however, is coherent only when zero is a meaningful centre of the response. If the response is strictly positive (or strictly negative), multiplication by weights ***ω***_*i*_ ∈ (0, 1] can move observations in one tail farther from the distributional bulk, stretching the tail rather than shrinking it toward a robust centre. Thus, although the weights arise from a robust procedure, the transformation ***ω***_*i*_*y*_*i*_ does not, in general, represent shrinkage toward a robust measure of location. We therefore define the weighted response as

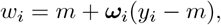

where *m* denotes a robust measure of central tendency, taken here as the median. This median-centred formulation weights deviations from the centre, shrinks observations toward *m*, and preserves the intended interpretation of the robust weighting. When *m* ≈ 0, the two formulations are approximately equivalent, as *m* + ***ω***_*i*_(*y*_*i*_ − *m*) ≈ ***ω***_*i*_*y*_*i*_. In practice, this approximation can be checked with simple summaries of the response distribution. Whether zero falls within the central region of the data, for example within the interquartile range, provides an initial diagnostic; when it does, the closeness of the median to zero can be assessed relative to response scale, for example through the ratio |*m*|*/s*_*y*_.

Weighted responses differ fundamentally from Box–Cox or Yeo–Johnson transformations because they provide no natural back-transformation. The weights ***ω***_*i*_ depend on residual diagnostics from observed responses and therefore cannot be assigned to future observations. Even so, median-centred robust weighting provides a principled exploratory device for testing whether shrinking contaminated responses toward a robust centre during model fitting improves predictive performance under contamination.

These six preprocessing strategies—two parametric transformation families (Box–Cox; Yeo–Johnson) with robust counterparts, two nonparametric remappings (winsorization; ranks), and two exploratory tail-control devices (median winsorization; robust weighting)—form a layered toolkit for robustifying RF at the point of data entry. Their shared logic is simple but consequential: weaken the leverage of extremes before the forest splits and averages, and the downstream ensemble becomes steadier, not by redesign, but by disciplined preprocessing of the response.

#### Four robust algorithm-based approaches

Robustifying RF at the algorithmic level targets the components through which contamination propagates—resampling, splitting, and aggregation—thereby reducing the leverage of outliers without relying on preprocessing. In what follows, we describe four robust algorithm-based strategies: (i) robust bootstrapping, (ii) robust splitting via MAE impurity, and robust aggregation via (iii) median- and (iv) quantile-based rules. Together, these modifications act where RF is most fragile—at the moments it samples, partitions, and pools—turning a variance-driven learner into a more contamination-tolerant predictor.

#### Bootstrap level

1. Standard bootstrap resampling can over-represent contaminated observations, yielding biased trees; robust bootstrapping mitigates this by sampling with observation-specific probabilities. Each observation *y*_*i*_ is assigned a sampling probability

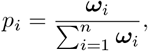

where ***ω***_*i*_ is the robust weight for observation *i*, computed as described earlier (*preprocessing-based approaches* – method 6), and the denominator normalizes probabilities. In practice, many RF implementations use observation weights internally, so the user supplies ***ω***_*i*_ rather than explicitly computing *p*_*i*_. By down-weighting high-residual observations during resampling, robust bootstrapping reduces their replication across trees, improving ensemble stability.

#### Split level

2. In regression RF, classical split selection minimizes within-node MSE around the node mean, i.e., it selects splits that minimize node impurity measured by squared deviations; under contamination, the node mean can be distorted by a few extremes, particularly in small nodes, compromising both node prediction and split choice. At the split level, robustness can be introduced by replacing the squared-error impurity criterion with an absolute-deviation criterion centered on the node median. This substitution makes split selection equivalent to minimizing MAE within each node—i.e., a least absolute deviations (LAD) principle at the node level—thereby down-weighting extreme responses in impurity calculations, limiting outlier leverage on split decisions, and producing more stable trees under contamination.

The impurity at node *t* is defined as

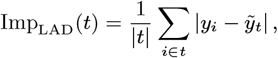

where 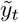 is the median of {*y*_*i*_ : *i* ∈ *t*} and |*t*| is the number of observations in node *t*.

Although alternative robust losses—such as the Huber loss—could, in principle, be embedded in the split criterion to further reduce sensitivity to outliers, most widely used RF implementations do not support user-defined impurity measures, which limits the practical deployment of such split-level modifications.

#### Aggregation level

3. Standard RF aggregates tree predictions by averaging; under contamination, a small subset of extreme tree predictions can bias the mean. A robust alternative aggregates by the median (the 0.5 conditional quantile), which dampens the effect of anomalous predictions and stabilizes the ensemble.

4. More generally, conditional-quantile aggregation, as in Quantile RF, supports robust point prediction (via the median) and uncertainty summaries (via other quantiles; 67).

Operationally, both median aggregation and quantile regression forests alter only the pooling step; tree construction remains identical to that of a standard RF. Under median aggregation, tree-wise point predictions are combined using their median rather than their mean, thereby reducing sensitivity to extreme tree predictions. By contrast, quantile regression forests retain the full set of training responses associated with the terminal nodes reached across trees, preserving distributional information that is then used at the aggregation stage to estimate the conditional response distribution and to derive point predictions such as the median, as well as other conditional quantile approaches explored in the literature [20]. Beyond the median, trimmed-mean and Winsorized-mean aggregation can further reduce sensitivity by discarding or shrinking extreme predictions prior to averaging; however, because our focus is point-prediction accuracy, we emphasize median-based aggregation as the most direct robust alternative.

Computational details of the methods are provided in the Supplementary Materials (Section *Details of model fitting*, Subsection *Computational resources and software*), while the notation used throughout is summarized in Tables 3 & S3.

**Table 3.**
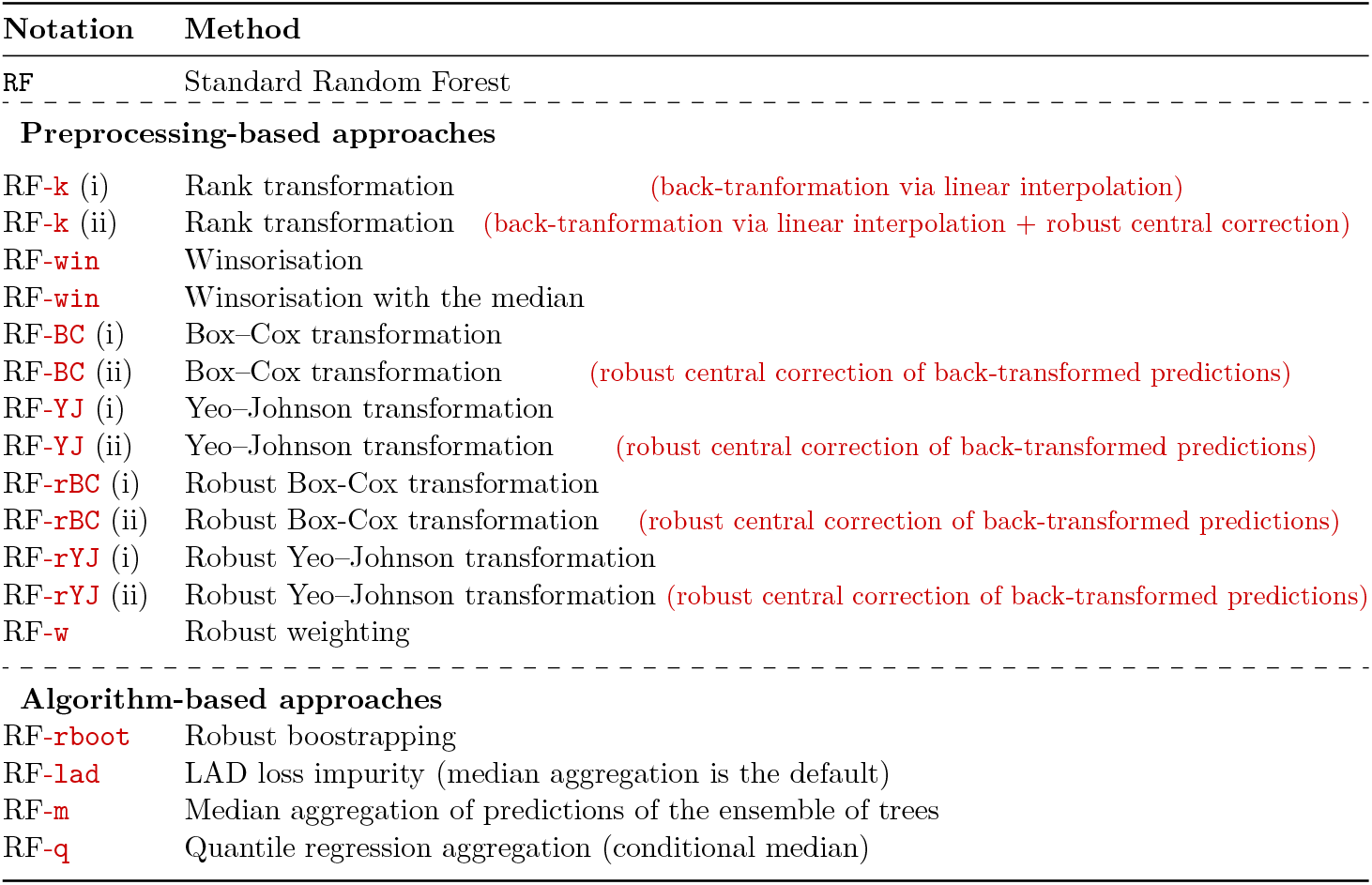
Pre-processing, algorithm-based, and hybrid strategies evaluated in the first simulation stage, including notation for each method.

### Data analysis

#### Simulated data

We adopt a sequential evaluation strategy—screen, retain, and stress-test—to preserve interpretability and avoid an uninformative combinatorial expansion of methods. Preprocessing- and algorithm-level remedies are first assessed separately under `shift` contamination; only competitive methods advance to `variance-inflated` contamination and, if still competitive, to `central variance-deflated`contamination and finally to `tail variance-deflated` contamination. The most competitive preprocessing and algorithm-level approaches are then combined into hybrid strategies, which undergo the same sequential screening; only combinations integrating the most effective components are retained, focusing inference on plausible robustification pathways rather than exhaustive, low-yield search. Each method was screened and retained or advanced as follows. Using the original (uncontaminated) data, the standard RF achieves a PA of approximately 0.755, which defines the baseline against which contamination-induced losses are evaluated. To operationalize the sequential screening across contamination scenarios, we imposed a minimum performance criterion set at ∼ 80% of this baseline, yielding a threshold of PA = 0.6. Methods attaining PA ≥ 0.6 under a given contamination setting were advanced to the next, more stringent stage, whereas methods with PA *<* 0.6 were excluded from further consideration. This rule ensures that only approaches preserving a substantial fraction of baseline predictive performance proceed to subsequent contamination regimes. From the best-performing preprocessing, algorithm-level, and hybrid strategies, we then estimate breakdown points by progressively increasing contamination levels. The sequential evaluation strategy is summarized in Figure 3.

**Fig. 3.**
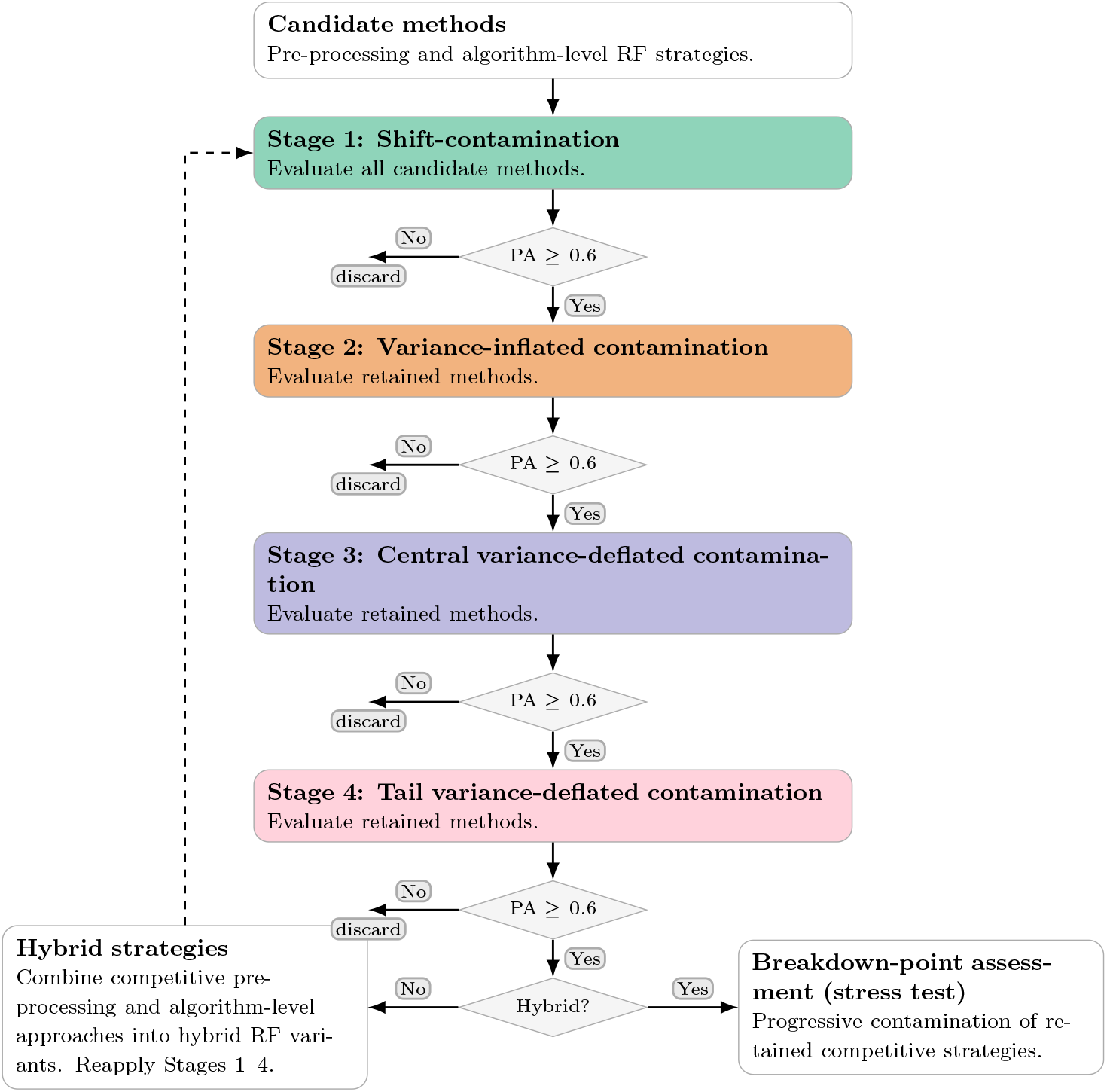
Diagrammatic illustration of the sequential evaluation strategy: preprocessing- and algorithm-level remedies are first evaluated separately, the best-performing approaches are then combined and re-evaluated, and the strongest overall strategy is pruned, assessed for breakdown point, and stress tested.

Further details of hyperparameter tuning, implementation choices, reproducibility safeguards, and other RF specifications are provided in the Supplementary Materials, Section *Details of model fitting*, Subsection *RF hyperparameter tuning and other specifics* (Table S4; Figure S7).

#### Real data

A unified GP pipeline is applied to all traits in the real plant and animal datasets to ensure consistent and rigorous model evaluation (Figure 4). The analytical framework follows the evaluation procedure described by **(author?)** [68] and ss implemented identically for each trait in the maize, soybean, wheat, and mice datasets. For every trait, the observations are randomly partitioned into two subsets: a training set for model estimation and hyperparameter tuning, and a hold-out test set for independent evaluation. Specifically, 20% of the observations are reserved as the test set, while the remaining 80% form the training set used for model fitting and tuning. The test set remains strictly isolated from all stages of model development and is used only for final predictive assessment and preliminary diagnostic checks. This design preserves most of the data for model estimation while maintaining a fully independent sample for evaluating predictive performance. In doing so, the pipeline establishes a disciplined and transparent foundation for comparing predictive models across traits and datasets. Hyperparameter tuning for each trait is conducted on the training set using five-fold cross-validation for both the standard RF and the best-performing robust RF methods. The 80% training set is partitioned into five equally sized folds. For each method, models are trained on four folds and validated on the remaining fold. This process is repeated five times so that each fold serves once as the validation set. The resulting cross-validation cycle produces a stable estimate of predictive performance across folds and allows direct comparison among candidate hyperparameter configurations. The configuration yielding the best average performance across the five folds is selected for each trait and method. To ensure strict comparability across methods, the same fixed random number seed is used so that all methods employ exactly the same training-set folds during hyperparameter tuning. This controlled design removes an important source of stochastic variation and ensures that differences in performance arise from the models themselves rather than from differences in data partitioning.

**Fig. 4.**
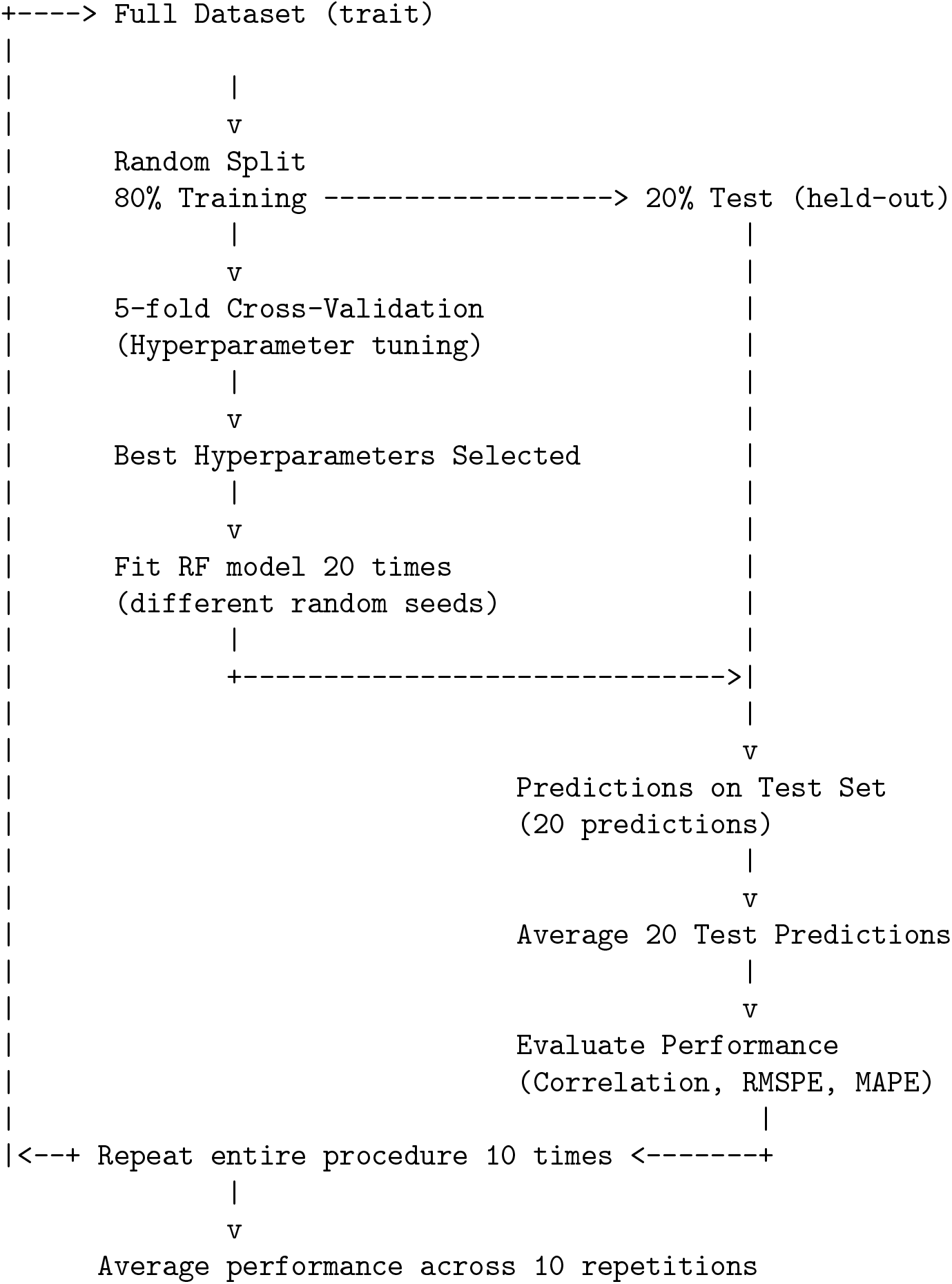
Training–testing framework used for model evaluation for each trait in each real data set. Predictions from the 20 RF runs on the held-out test set are averaged and used to compute performance metrics.

Final model estimation and prediction are then performed using repeated fits to stabilize stochastic variation inherent in RF training. After selecting the optimal hyperparameter settings for each trait–method combination, each model is fitted 20 times to the full training set. The 20 runs differ only in the random number seed used during model initialization. To evaluate predictive performance, predictions from the 20 model fits are first averaged and then compared with the corresponding observed values in the independent 20% hold-out test set that had not been used during training or tuning.

Let *ŷ*^(*i*)^ denote the predicted values obtained from run *i*, where *i* = 1, …, 20. Final predictions for the test set are obtained by averaging across the 20 fits:

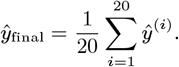

Averaging predictions across repeated fits reduces variability induced by random seeds and yields more stable and reliable estimates of predictive performance. Through this sequence—independent data partitioning, controlled cross-validation, and stabilized prediction—the pipeline provides a coherent and reproducible framework for evaluating GP models on real plant and animal data.

### Predictive performance assessment

Genomic prediction estimates genetic values—such as genomic estimated breeding values (GEBVs)—from molecular markers using statistical or machine-learning models. Because the central objective is accurate numerical prediction, performance is most directly assessed using error-based criteria, including the prediction-error metrics defined above. Predictive accuracy is also widely reported because it is scale-free and easily interpretable, summarizing how well predicted and true values align in rank. Yet PA can mask systematic bias or rescaling and, when absolute error is the primary concern, may not faithfully represent prediction quality.

Genomic selection, in contrast, is a downstream decision framework that uses genomic predictions to rank candidates and choose superior individuals for breeding. Here, correct ordering—not precise calibration—is the operative goal; thus, PA becomes the more decision-relevant metric, capturing the fidelity of the ranking that selection ultimately acts upon.

Accordingly, GP prioritizes accurate values and is best evaluated by error-based measures, whereas genomic selection prioritizes accurate ranks and is therefore best summarized by PA—two aims, two evaluation targets, one coherent pipeline from prediction to selection.

#### Simulated data

Predictive performance of the methods is assessed on the simulated animal data using PA, the root mean squared prediction error (RMSPE), and the mean absolute prediction error (MAPE). Let *y*_*i*_ denote the true genomic breeding values (TGBVs) and *ŷ*_*i*_ the predicted genomic breeding values (PGBVs). The metrics are defined as

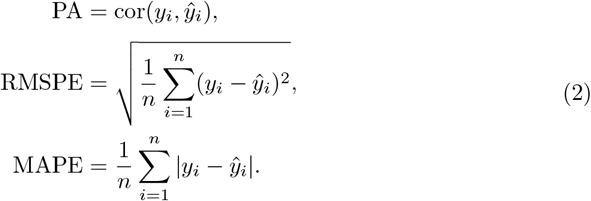

Although RMSPE and MAPE are sometimes reported as percentage errors, we compute both on the original response scale, emphasizing absolute predictive fidelity rather than scale-normalized deviation.

We evaluate model performance using the predefined training–validation split of trait 1 (T1) dataset. RF models are trained on 3000 individuals in the training population and evaluated on the remaining 1020 individuals using their known TGBVs as the external validation target. In contrast to earlier studies that tuned regularization parameters via cross-validation [7, 8] or emphasized reporting PAb [6], we exploit the independent validation population embedded in the QTLMAS simulation design. When such an external validation population is unavailable, RF models are typically tuned and assessed using OOB prediction error (PE), a pragmatic surrogate for independent validation.

#### Real data

Predictive performance for the real plant and animal datasets was evaluated using complementary accuracy and error metrics computed from the final averaged predictions, *ŷ*_*final*_. These metrics comprised PAb, defined as the Pearson correlation between *ŷ*_*final*_ and the observed phenotypic values, together with RMSPE and MAPE. By combining a correlation-based measure with scale-dependent and relative error measures, this framework captures multiple dimensions of predictive performance and thus provides a more balanced assessment of model behaviour across traits. To account for variation introduced by random data partitioning, the entire training–testing procedure was repeated 10 times using different random splits of each dataset. Performance metrics were calculated separately for each replicate, and the final reported values were obtained by averaging them across all 10 replicates. This repeated-splitting design reduces the influence of any single partition and yields more stable estimates of predictive performance. We then compared PAb and prediction-error measures (RMSPE and MAPE) descriptively across the 10 data splits to determine whether differences among methods reflected consistent methodological contrasts rather than random partition effects, and whether between-method variation in RMSPE and MAPE clarified relative error magnitudes and thereby provided an interpretable measure of predictive precision. The use of averaged predictions, multiple performance metrics, and repeated data splits provides a rigorous, reproducible, and methodologically coherent framework for evaluating model performance on real plant and animal data and for supporting robust comparative inference across methods.

### Simulation Results

The relationship between contamination and predictive performance is not invariably monotonic. In some scenarios, increasing the proportion of contamination does not immediately reduce PA or increase PEs. This pattern arises because contaminated observations do not always appear as clearly identifiable outliers. When perturbations remain within the range of the bulk of the data, they can act as masked points or inliers and thus blend partly into the underlying distribution. Moderate increases in contamination may therefore fail to cause an immediate deterioration in predictive performance, a result consistent with the masking effects well documented in the statistics literature [69, 70, 71].

#### Impact of contamination on predictive performance

Across contamination scenarios and contamination percentages, the predictive performance of RF deteriorates as contamination increases: PA of RF declines, whereas predictive error (PE), expressed as MAPE or RMSPE, increases (Table 4, Figures 5 & S8). Within each scenario, larger `shift` magnitudes under shift contamination and larger variances under `variance-inflated` contamination are consistently associated with lower PA and higher PE. By contrast, `variance-deflated` contamination is the least harmful, showing a reduction in PA relative to the uncontaminated benchmark but remaining comparatively stable across contamination percentages and variance levels. This pattern aligns with the expected non-robustness of standard RF: bootstrap resampling, mean-based splitting, and mean aggregation are each sensitive to outliers, so contamination propagates from sampling and partitioning to ensemble prediction, strengthening the case for robustification under realistic data conditions.

**Table 4.**
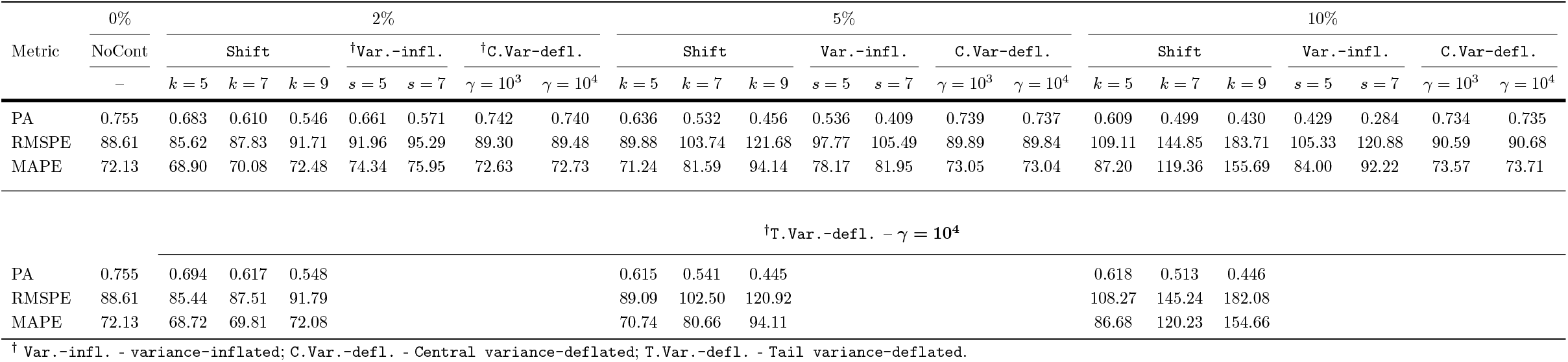
Estimated predictive accuracy (PA) and prediction errors, root mean prediction error (RMSPE) and mean absolute prediction error (MAPE), of random forests under non-contaminated (NoCont) and contaminated scenarios (contaminant *G* ∼ *Normal*).

| Metric | 0% |  | 2% |  |  |  |  |  |  | 5% |  |  |  |  |  |  | 10% |  |  |  |  |  |  |
| --- | --- | --- | --- | --- | --- | --- | --- | --- | --- | --- | --- | --- | --- | --- | --- | --- | --- | --- | --- | --- | --- | --- | --- |
| | NoCont | Shift | | | †Var.-infl. | | †C.Var-defl. | | Shift | | | Var.-infl. | | C.Var-defl. | | Shift | | | Var.-infl. | | C.Var-defl. | | |
| | – | k = 5 | k = 7 | k = 9 | s = 5 | s = 7 | $\gamma = 10^3$ | $\gamma = 10^4$ | k = 5 | k = 7 | k = 9 | s = 5 | s = 7 | $\gamma = 10^3$ | $\gamma = 10^4$ | k = 5 | k = 7 | k = 9 | s = 5 | s = 7 | $\gamma = 10^3$ | $\gamma = 10^4$ | |
| PA | 0.755 | 0.683 | 0.610 | 0.546 | 0.661 | 0.571 | 0.742 | 0.740 | 0.636 | 0.532 | 0.456 | 0.536 | 0.409 | 0.739 | 0.737 | 0.609 | 0.499 | 0.430 | 0.429 | 0.284 | 0.734 | 0.735 |  |
| RMSPE | 88.61 | 85.62 | 87.83 | 91.71 | 91.96 | 95.29 | 89.30 | 89.48 | 89.88 | 103.74 | 121.68 | 97.77 | 105.49 | 89.89 | 89.84 | 109.11 | 144.85 | 183.71 | 105.33 | 120.88 | 90.59 | 90.68 |  |
| MAPE | 72.13 | 68.90 | 70.08 | 72.48 | 74.34 | 75.95 | 72.63 | 72.73 | 71.24 | 81.59 | 94.14 | 78.17 | 81.95 | 73.05 | 73.04 | 87.20 | 119.36 | 155.69 | 84.00 | 92.22 | 73.57 | 73.71 |  |
| †T.Var.-defl. – $\gamma = 10^4$ | | | | | | | | | | | | | | | | | | | | | | | |
| PA | 0.755 | 0.694 | 0.617 | 0.548 |  |  |  |  | 0.615 | 0.541 | 0.445 |  |  |  |  | 0.618 | 0.513 | 0.446 |  |  |  |  |  |
| RMSPE | 88.61 | 85.44 | 87.51 | 91.79 |  |  |  |  | 89.09 | 102.50 | 120.92 |  |  |  |  | 108.27 | 145.24 | 182.08 |  |  |  |  |  |
| MAPE | 72.13 | 68.72 | 69.81 | 72.08 |  |  |  |  | 70.74 | 80.66 | 94.11 |  |  |  |  | 86.68 | 120.23 | 154.66 |  |  |  |  |  |

$\dagger Var.-infl.$ - variance-inflated; $C.Var.-defl.$ - Central variance-deflated; $T.Var.-defl.$ - Tail variance-deflated.

**Fig. 5.**
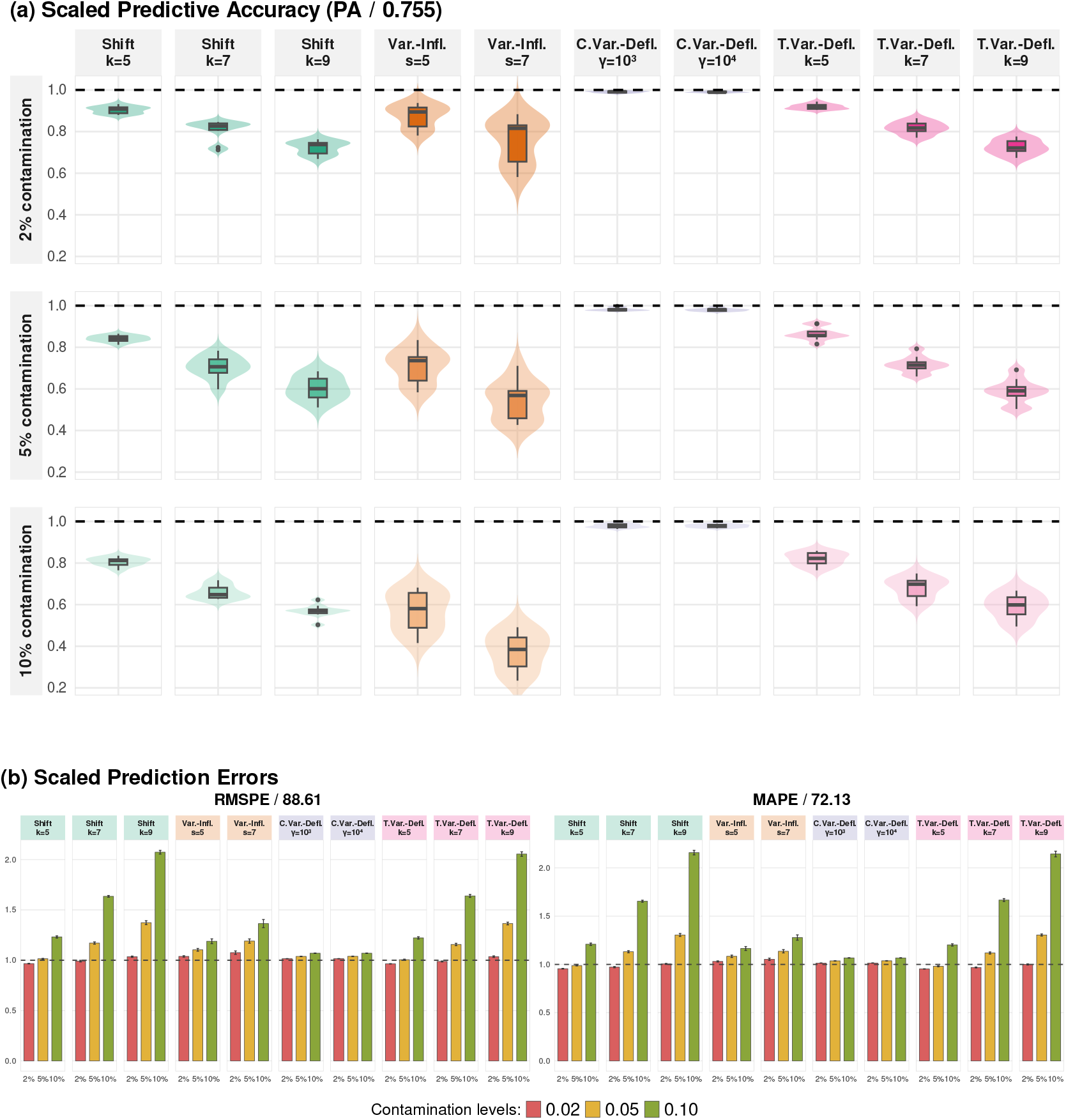
Performance of the Random Forests (RF) model under the four contamination scenarios: `Shift, Variance-Inflated, Central Variance-Deflated`, and `Tail Variance-Deflated`. Panel (a) shows predictive accuracy (PA) across contamination scenarios and levels. Panel (b) summarises prediction errors (RMSPE & MAPE) by contamination level within each scenario. Bars represent means across simulation runs and error bars denote standard errors; dashed lines indicate RF benchmark values.

Among contamination types, `variance-inflated` contamination produces the largest loss of PA, with PA decreasing from 0.755 to 0.284 (a 62.4% decrease). In contrast, shift contamination produces the largest inflation in PE, with MAPE increasing from 72.13 to 155.69 (a 115.8% increase) and RMSPE increasing from 88.61 to 183.71 (a 107.3% increase) (Table 4, Figure 5). Collectively, these results show that contamination degrades RF both by eroding rank fidelity (PA) and by amplifying absolute error (PE), with the dominant failure mode depending on whether contamination primarily inflates variability or displaces the response distribution.

#### Stage 1: Shift contamination

##### Preprocessing-based approaches

Across `shift` contamination scenarios, all preprocessing remedies improve on the baseline RF, indicating that simple data-level adjustments can substantially mitigate sensitivity to distributional displacement (Table S5). Performance, however, differs by objective: the weighting strategy (RF-`w`) delivers the highest PA in most settings, whereas the Box–Cox (RF-`BC`) and Yeo–Johnson (RF-`YJ`) transformations more consistently minimize PEs. Notably, only the rank-based (RF-`k`) and weighting (RF-`w`) approaches sustain PA above 0.7 across all shift magnitudes and contamination levels, showing superior robustness in preserving rank fidelity under contamination. In the rank (RF-`k`), Box–Cox (RF-`BC`), and Yeo–Johnson (RF-`YJ`) settings, the central correction applied to predictions further reduces error, with non-negligible gains in several scenarios, reinforcing that transformation-based preprocessing can both raise PA and curb PE relative to the standard RF.

##### Algorithm-based approaches

Across `shift` contamination scenarios, the LAD split criterion (RF-`lad`) yields only sporadic improvement over the benchmark RF and is the weakest algorithm-based remedy overall, with the largest PEs (Table S6). Although it does not meet the PA screening threshold, robust bootstrapping (RF-`rboot`) typically delivers the smallest PEs, indicating that down-weighting contaminated cases can reduce absolute error even when rank fidelity remains limited. The strongest algorithm-level performers are the aggregation-based remedies—median (RF-`m`) and quantile (RF-`q`)—which most consistently improve performance relative to the standard RF; however, neither sustains PA above 0.7 across all shift magnitudes and contamination levels.

##### Preprocessing-based vs Algorithm-based approaches

In the uncontaminated scenario, standard RF achieves a PA of 0.755 and serves as the reference benchmark. Most preprocessing and aggregation-based methods exhibit only minor reductions relative to this value. Classical transformations (RF-`BC`, RF-`win` (i)) and robust transformations (RF-`rBC`) reduce PA by approximately 0.7 − 1.1%, while rank-based preprocessing (RF-`k`), median aggregation (RF-`m`), and robust bootstrap (RF-`rboot`) incur modest reductions of about 2 − 2.5%. The robust weighting (RF-`w`) and quantile aggregation (RF-`q`) approaches show slightly larger decreases, around 3%. Notably, RF-`rYJ` performs marginally better than standard RF, with a small improvement of 0.3%. In contrast, the LAD-based impurity variant (RF-`lad`) exhibits a substantial reduction of 15.6%, indicating a considerable loss of efficiency in the absence of contamination (Table 5).

**Table 5.** Robustness under shift contamination and relative efficiency in the uncontaminated scenario. The last column reports the percentage difference in predictive accuracy at 0% contamination (PA_0%_) relative to the standard RF.

| Method | $PA_{0\%}$ | Min PA (Shift) | % Loss | %$\Delta$ (0%) |
| --- | --- | --- | --- | --- |
| RF | 0.755 | 0.430 | 43.0 | 0.0 |
| **Preprocessing-based Approaches** |  |  |  |  |
| RF-**k** | 0.739 | 0.718 | 2.8 | -2.1 |
| RF-**win** (i) | 0.747 | 0.433 | 42.0 | -1.1 |
| RF-**win** (ii) | 0.685 | 0.377 | 45.0 | -9.3 |
| RF-**BC** | 0.747 | 0.646 | 13.5 | -1.1 |
| RF-**YJ** | 0.755 | 0.672 | 11.0 | 0.0 |
| RF-**rBC** | 0.750 | 0.503 | 32.9 | -0.7 |
| RF-**rYJ** | 0.757 | 0.506 | 33.2 | +0.3 |
| RF-**w** | 0.731 | 0.722 | 1.2 | -3.2 |
| **Algorithm-based Approaches** |  |  |  |  |
| RF-**rboot** | 0.737 | 0.545 | 26.1 | -2.4 |
| RF-**lad** | 0.637 | 0.491 | 22.9 | -15.6 |
| RF-**m** | 0.738 | 0.625 | 15.3 | -2.3 |
| RF-**q** | 0.730 | 0.645 | 11.6 | -3.3 |

While the efficiency losses under uncontaminated data are generally modest, the behaviour of the competing methods changes markedly in the presence of `shift` contamination. In this setting, the standard RF exhibits a substantial degradation, with a 43% reduction in PA. A similarly pronounced deterioration is observed for the winsorisation approaches, particularly variant (ii), which shows the largest loss (45%). The robust transformation-based methods RF-`rBC` and RF-`rYJ` reduce this impact but still experience losses of approximately 33% (Table 5).

In contrast, classical transformation approaches such as RF-`BC` and RF-`YJ` demonstrate improved stability, limiting the PA reduction to about 11 − 14%. The rank-based preprocessing method RF-`-k` performs considerably better, with only a 2.8% loss. The most stable behaviour is observed for the robust weighting approach RF-`w`, which shows a negligible reduction of approximately 1.2%. Among the algorithm-based approaches, RF-`rboot` shows a moderate reduction (26.1%), indicating partial robustness to systematic shifts. The LAD impurity variant (RF-`lad`) also improves over classical RF but still experiences a 22.9% loss. Median aggregation (RF-m) further reduces the deterioration to 15.3%, while quantile aggregation (RF-`q`) limits the reduction to 11.6%, approaching the performance of the best transformation-based methods (Table 5).

Overall, the results suggest that the efficiency costs incurred by most robust modifications under clean data are small relative to the substantial gains in stability observed under `shift` contamination. The LAD impurity approach constitutes the main exception, as it sacrifices considerable baseline performance without achieving the highest robustness. In contrast, direct weighting and rank-based preprocessing offer the most favourable balance between efficiency and robustness, while aggregation-based strategies provide intermediate protection and classical RF with winsorisation remain highly sensitive to systematic shifts (Table 5).

Under `shift` contamination, the comparative pattern is clear: preprocessing-based remedies are the most effective at preserving PA across scenarios, implying that transforming the response can protect rank fidelity more consistently than modifying the forest alone. Within this class, RF-`k` and especially RF-`w` emerge as the most reliable candidates for advancing to `variance-inflated` evaluation because they stabilize PA with minimal degradation. However, this gain in PA does not always translate into the smallest PEs: median aggregation (RF-`m`) often yields lower absolute errors in heavily shifted settings, underscoring a pragmatic PA–PE trade-off that informs which methods to carry forward (Tables S5-S6; Figures 6 & S9).

**Fig. 6.**
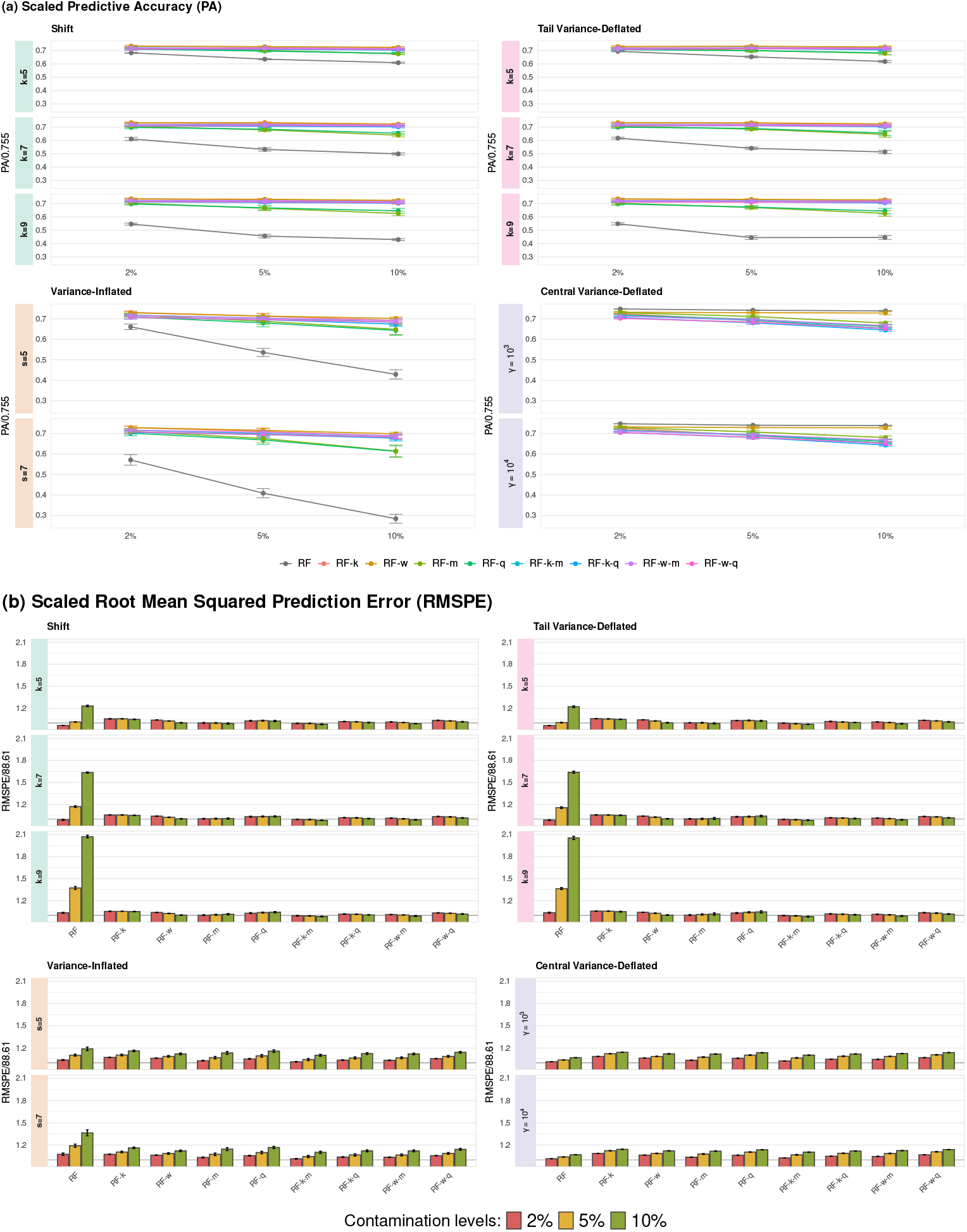
Predictive performance (scaled predictive accuracy - PA; scaled root mean squared prediction error - RMSPE; and scaled mean absolute prediction error - MAPE) of the methods under the four contamination scenarios: `Shift, Variance-Inflated, Central Variance-Deflated`, and `Tail Variance-Deflated`. Points represent the mean values at contamination levels of 2%, 5%, and 10%, whereas vertical bars indicate standard errors across simulation runs. Facets correspond to the contamination parameter values (*k, s*, or *γ*, depending on the scenario).

##### Screening Results

Screening under `shift` contamination eliminated methods that failed to reach the *PA* ≥ 0.6 threshold and were therefore not carried forward in the sequential evaluation. Winsorisation (RF-`win`), robust Box–Cox (RF-`rBC`), robust Yeo–Johnson (RF-`rYJ`), LAD (RF-`lad`), and robust bootstrap (RF-`rboot`) fell below the cutoff and were not advanced to the `variance-inflated` or `variance-deflated` stages. By contrast, RF-`k`, RF-`BC`, RF-`YJ` and RF-`w` met the screening criterion and progressed to the next stage, thereby concentrating downstream evaluation on remedies with demonstrable screening performance (Figure 7).

**Fig. 7.**
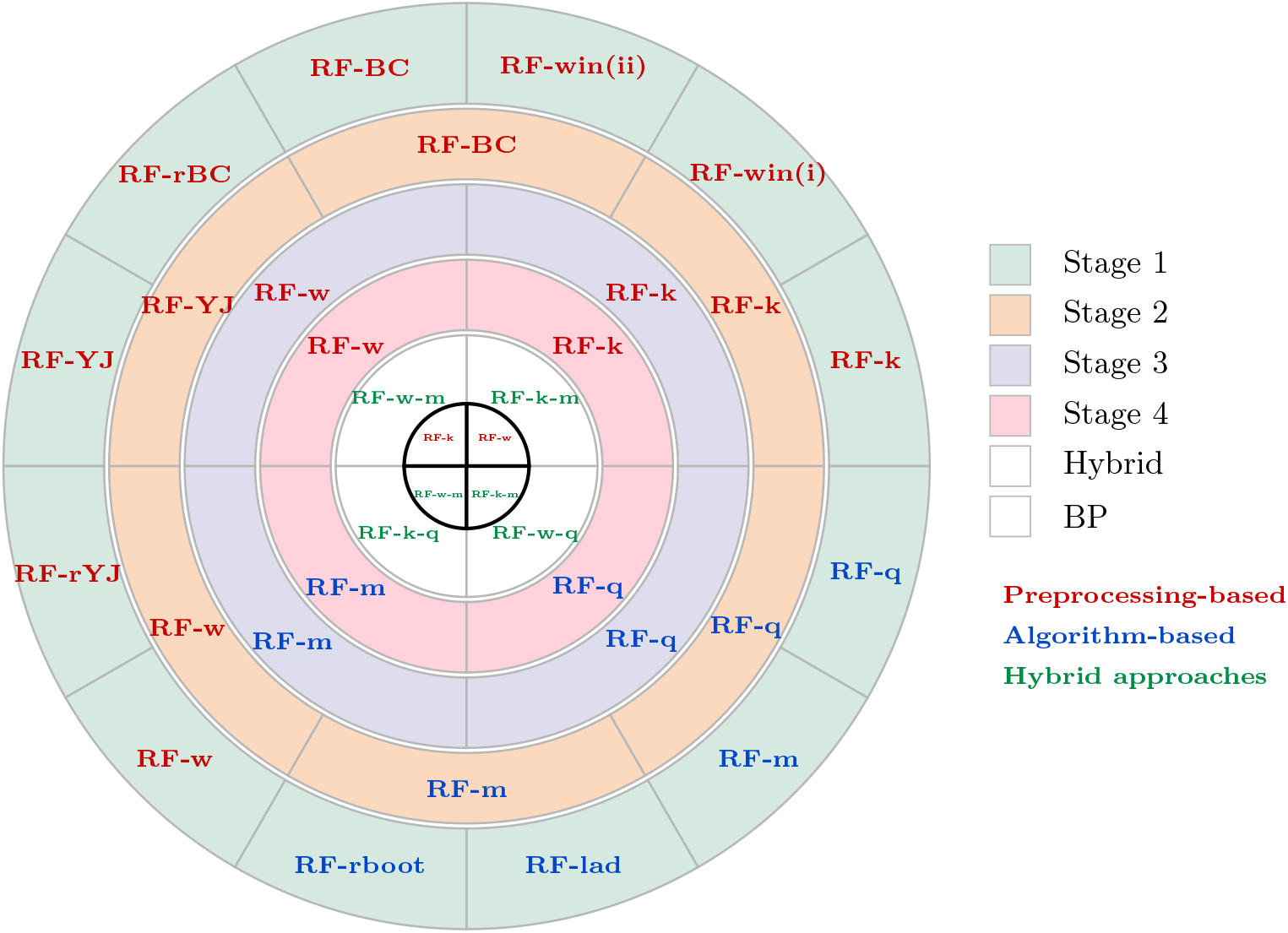
Concentric-ring representation of the five-stage screening framework. The outermost ring (Stage 1) displays the initial set of 12 candidate methods (10 approaches). Subsequent inner rings illustrate the progressive reduction to 6 and then 4 retained procedures. The innermost white ring corresponds to the breakdown-point assessment (Stage 5), in which RF-`k` and RF-`w` are retained.

The comparatively weak screening performance of RF-`rBC` and RF-`rYJ` is consistent with the design of the corresponding robust transformation proposed by [58], which target central normality, that is, approximate normality in the bulk of the data while allowing outliers to remain outlying. Although this property is advantageous for robust estimation and inference, it may be less well suited to prediction settings in which the full response distribution informs model fitting. In the `shift` contamination scenarios both the RF-`rBC` and RF-`rYJ` yielded lower predictive accuracies than their classical counterparts. One possible explanation is that the emphasis on robustness in the central region may preserve shift-induced structure in the data, thereby influencing the predictive performance of RF.

#### Stage 2: Variance-inflated contamination

##### Preprocessing-based approaches

Under `variance-inflated` contamination, the Box–Cox (RF-`BC`) and Yeo–Johnson (RF-`YJ`) transformations track the benchmark RF closely, offering little improvement and thus ranking as the weakest preprocessing remedies among those advanced from the `shift` contamination stage (Table S7a, Figures 6 & S9). In contrast, the weighted strategy (RF-`w`) is consistently strong: it attains a mean estimated PA above 0.7 in all but one scenario, with the only sub-threshold value remaining close (0.698), and it delivers the smallest PEs under the most adverse settings (5% with s = 7, and 10% with *s* = 5 and *s* = 7). The rank-based approach (RF-`k`) also preserves high PA—exceeding 0.7 in all but two scenarios (0.692 and 0.688), both near the threshold—but it produces larger PEs than RF-`w` across all scenarios (Table S7a, Figures 6 & S9). Overall, these results indicate that preprocessing can materially strengthen RF under `variance inflation`, with RF-`w` providing the most reliable balance of high PA and low PE, especially as contamination severity increases.

##### Algorithm-based approaches

Under `variance-inflated` contamination, the median-aggregation (RF-`m`) and quantile-based (RF-`q`) strategies perform similarly, but RF-`m` is consistently marginally stronger (Table S7b, Figures 6 & S9). This advantage is already evident in the uncontaminated setting, where RF-`m` starts from a PA of 0.738 compared with 0.730 for RF-`q`, and it persists across all `variance-inflated` scenarios. The same pattern holds for PE: RF-`m` generally yields slightly lower errors than RF-`q`, although uncertainty bands overlap in some settings when comparing conservative error ranges (e.g., subtracting the largest sd from RF-`m` PEs and adding the largest sd to RF-`q` PEs) (Table S7b, Figures 6 & S9).

##### Preprocessing-based vs Algorithm-based approaches

Under the `variance-inflated` contamination, for both the preprocessing and algorithmic methods, there was marked variation in the percentage loss in PA, computed as the difference between the PA for the uncontaminated (0%) scenario and the smallest observed PA across all `variance-inflated` scenarios (s = 5, 7; 2%, 5%, 10%) (Table S8). The standard RF shows a marked deterioration, with a 62.4% reduction in PA. Among preprocessing strategies, the classical transformation approaches RF-`BC` and RF-`YJ` provide essentially no protection, exhibiting losses comparable to standard RF (61.0% and 62.5%, respectively; Table S8).

In contrast, the rank-based preprocessing RF-`k` substantially limits the deterioration, with a loss of only 6.9%. The robust weighting approach RF-`w` displays the strongest stability, with a negligible reduction of 4.5%. The algorithm-based aggregation strategies yield intermediate robustness: both median aggregation (RF-`m`) and quantile aggregation (RF-`q`) reduce the degradation relative to standard RF, but still incur losses of approximately 16% (Table S8). Overall, these results suggest that `variance inflation` is particularly damaging for methods that do not directly address heteroscedastic contamination, while rank-based preprocessing and robust weighting provide the most effective protection.

Under `variance-inflated` contamination, the post-screening comparison sharpens the same practical trade-off: preprocessing-based strategies—especially RF-`k` and RF-`w`—most consistently retain PA across scenarios, indicating superior preservation of rank fidelity under heteroscedastic contamination. Algorithmic aggregation, by contrast, tends to improve performance through a different pathway: median aggregation (RF-`m`) more often delivers smaller PEs, curbing absolute error even when gains in PA are less pronounced (Table S7a-b, Figures 6 & S9). In aggregate, these results show that robustification remains mechanism-specific—transformations protect ordering, whereas aggregation more directly controls error magnitude—thereby guiding which approaches merit advancement to the variance-deflated stage.

##### Screening Results

Screening under `variance-inflated` contamination identifies a small set of robustification strategies that retain acceptable predictive fidelity and therefore warrant progression to the next stage of the sequential evaluation. Specifically, the Box–Cox (RF-`BC`) and Yeo–Johnson (RF-`YJ`) transformations fail to meet the PA *>* 0.6 threshold under `variance inflation` and are therefore not advanced to the `variance-deflated` stage. By contrast, only RF-`k` and RF-`w` (preprocessing-based) and RF-`m` and RF-`q` (algorithm-based) satisfy the screening criterion and are retained for subsequent evaluation, thereby concentrating inference on the most resilient candidates while avoiding low-yield expansion of the method set (Figure 7).

#### Stage 3: Central variance-deflated contamination

`Central variance-deflated` contamination emerges as the least deleterious scenario, confirming expectations based on the relative performance of the preprocessing and algorithmic strategies that advanced from the `shift` and `variance-inflated` stages. Relative to the standard RF, PA remains high and PEs remain low across settings (Table S9; Figures 6 & S9). Accordingly, none of the robustification strategies meaningfully improves on the standard RF, yet all retain acceptable performance. Under this contamination regime, therefore, the comparison shifts from recovery to stability, emphasizing the capacity of competing methods to preserve predictive reliability rather than to rescue degraded performance.

##### Preprocessing-based approaches

Among the preprocessing remedies carried forward, robust weighting (RF-`w`) is the most stable: it maintains PA above 0.7 across all settings and consistently delivers the smallest PEs, indicating strong preservation of rank fidelity with tight error control under variance reduction. By contrast, rank-based preprocessing (RF-`k`) exceeds PA = 0.7 only under the 2% contamination scenarios and declines to approximately 0.650 in two settings. Although RF-`k` starts with a slightly higher PA in the uncontaminated case (0.739 vs 0.731 for RF-`w`), its steeper deterioration as `variance-deflation` intensifies suggests that performance is more sensitive to the back-transformation step than to the forest fit itself. Overall, RF-`w` provides the clearest preprocessing advantage under `variance-deflation`—robust, consistent, and predictably accurate (Table S9a; Figures 6 & S9).

##### Algorithm-based approaches

Under variance-deflated contamination, the median-aggregation strategy (RF-`m`) is the strongest algorithm-level remedy, achieving estimated PA above 0.7 in all scenarios except the two 10% worst-case settings (0.679 and 0.680). By contrast, the quantile-based approach (RF-`q`) exceeds PA = 0.7 only under the 2% contamination scenarios and declines to 0.661 in both 10% cases. RF-`m` also delivers consistently, albeit modestly, smaller PEs than RF-`q`, indicating more stable point prediction as `variance-deflation` intensifies (Table S9b; Figures 6 & S9).

##### Preprocessing-based vs Algorithm-based approaches

Under this scenario, the benchmark RF is only mildly affected (PA reduction = 2.3%), confirming that `central variance-deflation` has limited impact on PA for the baseline method (Table S10). Against this already-stable baseline, RF-`w` is the most resilient strategy, showing a negligible PA loss of 0.4%. RF-`k` degrades more noticeably (11.8%), consistent with additional sensitivity introduced by rank-based recovery of predictions. The aggregation-based strategies occupy an intermediate position: median aggregation (RF-`m`) and quantile aggregation (RF-`q`) incur PA losses of 8.0% and 9.5%, respectively. Thus, even when `variance-deflation` is comparatively benign, methods differ in how well they preserve rank fidelity, with RF-`w` remaining the most stable option in terms of PA (Table S10).

Viewed jointly, the variance-deflated results reinforce a consistent trade-off observed in earlier stages: robust preprocessing—especially RF-`w`—best preserves PA, whereas robust aggregation—especially RF-`m`—offers the most direct, if modest, reductions in PEs. Under `variance-deflation`, however, the key message is not recovery from failure but the maintenance of already-strong performance, a regime in which stability and parsimony become the primary criteria for prioritizing remedies (Table S9a-b; Figures 6 & S9).

##### Screening Results

All methods from this stage are advanced to the next stage (Figure 7).

#### Stage 4: Tail variance-deflated contamination

##### Preprocessing-based approaches

Under `tail variance-deflated` contamination, standard RF exhibits a substantial degradation in PA, with PA decreasing from 0.755 to a minimum of 0.445 (41.1% loss). This confirms that highly concentrated contamination in the tails can severely distort tree splits and terminal node predictions, particularly at higher contamination levels and larger shift magnitudes (Tables S11a & S12; Figure 6).

In contrast, the preprocessing-based approaches remain remarkably stable. The ranking-based method RF-`k` shows only a modest reduction in PA, with minimum values around 0.716 − 0.723 even at 10% contamination, corresponding to losses below 3.5%. Similarly, the weighting strategy RF-`w` exhibits near invariance across all settings, with PA consistently around 0.723 − 0.733 and losses below 1.2% (Tables S11a & 12S; Fig 6).

The PE measures (RMSPE and MAPE) further support this pattern. While standard RF experiences dramatic increases in error at 5% and 10% contamination, both RF-`k` and RF-`w` maintain stable error levels with very small variability across runs. These results indicate that preprocessing mechanisms effectively mitigate the influence of concentrated tail perturbations before model fitting (Table S11a; Figures 6 & S9).

##### Algorithm-based approaches

The algorithm-based approaches display heterogeneous behavior under `tail variance-deflation`. The median-based impurity measure (RF-`m`) retains strong robustness, with PA remaining above 0.626 even in the most extreme setting and losses considerably smaller than those of standard RF. Moreover, its RMSPE and MAPE remain comparatively stable, confirming that aggregation at the terminal node level reduces sensitivity to highly concentrated tail outliers (Table S11b; Figures 6 & S9).

In contrast, the quantile-based approach (RF-`q`) shows a more noticeable degradation in PA as contamination intensity increases, with minimum PA values around 0.643 at 10%. Although still substantially more stable than standard RF, its performance deteriorates more than that of RF-`m`, suggesting that quantile aggregation may remain partially influenced by extreme but highly concentrated tail observations (Table S11b; Figure 6).

Overall, among algorithm-based strategies, RF-`m` appears more resistant to `tail variance-deflated` contamination, whereas RF-`q` provides intermediate robustness (Table S11b; Figures 6 & S9).

##### Preprocessing-based vs Algorithm-based approaches

Table S12 confirms the pronounced vulnerability of standard RF under `tail variance-deflation`, with a 41.1% reduction in PA. In contrast, preprocessing-based methods exhibit near invariance, with RF-`w` showing no observable loss and RF-`k` incurring only a 3.1% reduction. Among algorithm-based strategies, RF-`m` and RF-`q` provide substantial improvements relative to standard RF, though their losses (15.2% and 11.9%, respectively) remain larger than those of preprocessing-based approaches. These findings indicate that mitigating tail concentration effects prior to model fitting is more effective than relying solely on robust aggregation within the tree-building procedure.

##### Screening Results

All methods passed this screening stage and advanced to the next stage, showing that each satisfied the baseline performance criteria (Figure 7). For the rank method, only the central correction, RF-`k` (ii), was retained to ensure consistency; the notation (ii) is omitted from the text but retained in the tables for reference.

#### Hybrid approaches

Screening of the hybrid approaches in the sequential analysis evaluates four hybrid candidates—RF-`k-m`, RF-`k-q`, RF-`w-m` and RF-`w-q`- each combining one of the two top preprocessing-based methods with one of the two top algorithm-based methods (Tables S2, S13-S17). This focused set therefore carries forward only the strongest components into direct comparison, allowing the relative performance of the hybrid remedies to be assessed across the full range of contamination scenarios. Relative to standard RF (benchmark PA= 0.755), the hybrid approaches incur only a modest efficiency cost under clean data conditions, with baseline losses ranging from 3.6% for RF-`k-m` to 6.1% for RF-`w-q` (Table S17). Thus, integrating preprocessing and aggregation mechanisms does not materially compromise PA when contamination is absent. Across the four simulation scenarios— `shift, variance inflation, central variance-deflation`, and `tail variance-deflation` the hybrid methods nonetheless show clearly differentiated yet consistently robust performance. The central result is therefore both simple and strong: the hybrids sacrifice little under clean conditions but gain substantially under contamination.

`Shift` contamination sharply degrades the standard Random Forest, but leaves the hybrid methods largely intact. By contrast, all four hybrid approaches remained highly stable, with PA losses not exceeding 2.6% and PE varying by no more than 3.2% in absolute value; notably, RF-`w-m` showed complete invariance with 0.0% PA loss (Table S17). These results show that the hybrid methods are highly resistant to mean displacement in the response and that the weighting-plus-median combination provides especially strong protection in this regime. Under `shift` contamination, then, the decisive contrast is not among hybrids but between the standard RF and the hybrid class as a whole, whose superiority is large, consistent, and practically consequential Table S13.

`Variance inflation` imposes the strongest stress on the standard RF and reveals most clearly the value of hybridization. As contamination increased to the most extreme setting (10% and *s* = 7), the PA of the standard RF fell by 62.4%, while PE rose by 27.8% in RMSPE and 36.4% in MAPE (Figures 6 & S9; Tables S14 & S17). In striking contrast, all four hybrid approaches remained stable, with PA decreasing by only 3.5% to 6.1% and PE increasing by only 3.9% to 10.5%; within this group, the weighting-based combinations RF-`w-m` and RF-`w-q` showed the smallest PA losses, 3.5% to 3.8% (Table S17). This pattern indicates that the combination of variance stabilization and robust aggregation effectively blunts the impact of extreme dispersion.

`Tail variance-deflation` again exposes a major weakness in the standard RF and restores a wide performance gap between the classical and hybrid methods. As contamination increased to the most extreme setting (10% and *k* = 9), the PA of the standard RF declined by 40.9% in the simulation summary and 41.1% (Table S17), while PE rose markedly by 105.5% in RMSPE and 114.4% in MAPE (Figures 6 & S9; Tables S15 & S17). By contrast, all four hybrid approaches showed only minor declines in PA, ranging from 0.0% to 2.9%, while their PEs varied from −2.9% to 0.0% and never exceeded their corresponding baseline values; RF-`w-m` again displayed the strongest stability, a 0.0% PA loss followed by RF-`w-q` (Table S17). These results show that the hybrids do not merely limit damage; in some configurations they all but neutralize the contamination effect. Under `tail variance-deflation`, as under `shift` and `variance inflation`, the decisive ranking is between the standard RF and the hybrid class as a whole, while ranking within the hybrid class remains finer and more tentative because the differences are small relative to their shared gain in robustness.

The overall pattern is therefore coherent, instructive, and practically important: hybridization improves robustness where robustness matters most, while imposing only a modest cost where contamination is weak. That balance gives the hybrid methods both their practical force and their scientific value, marking them as clearly superior to the standard RF under contamination and suggesting, with due caution, that RF-`w-m` is the most uniformly stable hybrid, while RF-`k-m` (ii) offers the sharpest compromise between efficiency and robustness.

##### Screening Results

Among the hybrid strategies, only the combinations that coupled the median with ranking (RF-`k-m`) or with weighting (RF-`w-m`) outperformed the corresponding versions that combined quantile RF regression with ranking or weighting. By contrast, the algorithm-based strategies using the median alone (RF-`m`) or quantiles alone (RF-`q`) did not advance, nor did they provide as strong a compromise between PA and PEs as when these same algorithmic components were integrated with the preprocessing strategies RF-`k` and RF-`w`, the two best overall methods across scenarios in terms of PA.

Only hybrid methods RF-`k-m` and RF-`w-m` are advanced to the next stage (Figure 7).

#### Breakdown point stress test

Breakdown point stress tests extend the simulation framework to examine how the most competitive preprocessing and hybrid approaches sustain predictive performance under extreme contamination. The main simulation study evaluates contamination levels up to 10%, which already represent severe perturbations in practical settings. To probe the limits of robustness more rigorously, the breakdown-point (BP) analysis increases the proportion of contaminated observations to 15%, 20%, and 25%, thereby subjecting the leading methods to progressively stronger stress tests. The analysis focuses on the most stable approaches identified earlier—preprocessing methods (RF-`k` and RF-`w`) and hybrid methods (RF-`k-m` and RF-`w-m`)—which advanced to the BP stage because they consistently maintained higher PA and lower PEs than competing alternatives. By concentrating on these best-performing approaches, the BP analysis directly evaluates whether robust preprocessing or hybrid strategies can meaningfully strengthen the predictive reliability of the standard random forest under extreme contamination, thereby providing a stringent test of their practical value for stabilizing GP.

##### Random forests

Breakpoint stress tests distinguish the standard RF from the hybrid procedures by revealing which methods retain stability as contamination intensifies. Under `shift` contamination, increasing contamination to the most extreme setting caused the standard RF to deteriorate sharply, with PA declining by 23.6% and 43.0% in the breakpoint stress comparison, while PE more than doubled, rising by 107.5% in RMSPE and 115.8% in MAPE (Tables S18-S19; Figures S10-S11).

Tables S18-S19 and Figures S10-S11 report stress-test results for the most competitive methods under *shift* contamination at 15%, 20%, and 25%. Unlike the monotone deterioration observed at 2%, 5%, and 10% contamination, PA does not continue to decline at these higher *shift* proportions. Instead, PA shows a systematic recovery as contamination rises from 15% to 25% across all shift magnitudes (*k* = 5, 7, 9). This pattern is unlikely to be a Monte Carlo artefact, because the standard deviations are small relative to the magnitude of the observed changes. In contrast, PEs (RMSPE and MAPE) increase monotonically with both contamination level and shift magnitude, indicating progressively poorer calibration even as rank-based performance partially rebounds. This divergence underscores that PA and error-based metrics quantify different facets of predictive performance—rank fidelity versus absolute error. This non-monotone PA response plausibly reflects a change in the structure of the contaminated training responses. At moderate contamination, shifted observations are sufficiently frequent to destabilize tree partitioning but insufficiently prevalent to form a coherent secondary regime, thereby disrupting ranking. As contamination increases further, the responses more closely approximate a structured mixture distribution; in effect, RF learns a more stable—yet systematically biased—mapping. The monotone rise in PEs is consistent with accumulating bias, whereas the partial recovery in PA is consistent with more stable partitioning and improved rank preservation. The stronger recovery at larger *shift* magnitudes (*k* = 7, 9), where separation between response regimes is clearer, further supports this mixture-based interpretation. Collectively, these results indicate that, when *shift* contamination is confined to the training set, extreme contamination can preserve—or even improve—selection-relevant ranking while substantially degrading predictive calibration.

Under `variance-inflated` contamination, RF performance deteriorates monotonically with contamination level and severity, consistent with progressively stronger heteroscedastic noise. Unlike the `shift` scenario, PA exhibits no recovery at high contamination levels, supporting the inference that the non-monotone PA pattern under *shift* arises from mixture-induced structure rather than contamination magnitude per se.

Stress-test results under `shift, variance-inflated`, and `tail variance-deflated` contamination reveal the relative vulnerability of RF and clarify the conditions under which robust preprocessing or algorithmic approaches are most needed to stabilize predictive performance. The results confirm the strong sensitivity of RF to `shift` and `variance-inflated` contamination. Under `shift` perturbations, PA deteriorates substantially, with losses exceeding 40% already at 10% contamination and remaining around 37% in the breakdown point scenarios. A much stronger deterioration occurs under `variance inflation`, where the minimum PA drops to 0.210, corresponding to a loss exceeding 70% relative to the uncontaminated baseline. This severe degradation indicates that dispersion-based contamination strongly disrupts the tree partitioning mechanism of RF, amplifying noise in the response and degrading the stability of the learned splits. In contrast, `central variance-deflated` contamination has only a minor influence on predictive performance. Even under breakdown point scenarios, PA remains close to the uncontaminated level, with losses below 5%, confirming that perturbations concentrated near the centre of the response distribution exert limited influence on the RF splitting structure. `Tail variance-deflated` contamination, however, produces a deterioration pattern resembling that observed under `shift` contamination: PA declines markedly as contamination increases, reaching losses above 35% in breakdown point scenarios. Collectively, these patterns show that RF is most vulnerable to contamination mechanisms that introduce strong dispersion or systematic displacement in the response variable, whereas centrally concentrated perturbations exert comparatively weaker effects on ranking and predictive stability.

Table S19 further situates these patterns within the extended breakdown point stress tests and clarifies how the severity of contamination alters RF performance relative to the main simulation study. The comparison between the primary simulations (contamination levels up to 10%) and the BP analysis (15–25%) confirms that variance inflation produces the most severe degradation. Under this mechanism, PA deteriorates sharply as contamination increases, exceeding 70% losses in the BP regime. `Shift` and `tail variance-deflated` contamination also generate substantial deterioration, with losses exceeding 35–40%, whereas `central variance-deflated` contamination again shows only minimal effects on PA even under extreme contamination levels. Overall, these results establish a clear hierarchy of vulnerability— `variance inflation` first, `shift` and `tail variance-deflation` second, and `central variance-deflation` last—thereby identifying the contamination regimes in which preprocessing-based or algorithm-based robustification becomes most critical for stabilizing RF predictions and preserving predictive reliability under contamination.

##### Preprocessing-based approaches

Across the breakdown point stress tests, the preprocessing strategies, RF-`k` and RF-`w`, both maintained PA above 0.7 in most cases, with RF-`w` showing slightly stronger overall PA performance than RF-`k`. PA losses were about 18% for both methods, occurring under `variance-inflated` contamination for RF-`k` and under `tail variance-deflated` contamination for RF-`w`. Prediction errors followed the same broad pattern of deterioration but remained moderate relative to those of the standard RF: RMSPE increased by approximately 28.5% for RF-`k` and 26.4% for RF-`w`, whereas MAPE rose by 28.5% for RF-`k` and 27.9% for RF-w (Table S18). These results show that both preprocessing approaches preserve substantial predictive stability under severe stress, with RF-`w` holding a slight edge in PA while both methods markedly limiting the breakdown seen in the standard RF.

##### Hybrid approaches

By contrast, all four hybrid approaches remained highly stable under shift contamination: PA losses did not exceed 2.6%, and PE varied by no more than 3.2% in absolute value; notably, RF-`w-m` showed complete invariance, with 0.0% PA loss (Table S19). Under `variance-inflation` contamination, the hybrid advantage became systematic. Here the hybrids did not merely outperform the standard RF; they formed a robust buffer against breakdown, with the weighting-based hybrids appearing strongest. `Central variance-deflation` produced a weaker stress response and a narrower separation among the hybrid methods. Even so, losses remained moderate and far smaller than the degradations observed under harsher contamination regimes. Under this milder contamination regime, the main result is therefore not dominance but durability: all hybrids remained competitive, although their margin of advantage largely receded. `Tail variance-deflation` contamination further clarified the relative standing of the leading hybrids. The PA of RF-`k-m` decreased by at most 15.9%, while its PE varied between 7.0% and 32.4%. For RF-`w-m`, PA varied between 9.7% and 0.14%, whereas PE varied between 8.8% and 0.0% (Tables S18-S19; Figures S10-S11).

Across all simulation scenarios and contamination percentages, these breakpoint stress tests sharpen the comparative standing of the hybrid methods. These results suggest that RF-`w-m` is somewhat more stable than RF-`k-m` in PA across the full stress spectrum, whereas RF-`k-m` (ii) offers the best compromise between clean-data efficiency and broad robustness (Table S19). The evidence therefore supports a cautious but meaningful ranking: RF-`w-m` emerges as the most uniformly stable procedure, especially under the harsher contamination regimes, whereas RF-`k-m` (ii) appears to balance baseline efficiency and robustness most effectively. Even so, the dominant result is their shared resilience; the hybrids differ at the margin, but they converge in their ability to withstand contamination far better than the standard RF.

These results show that the hybrid approaches achieve a favorable balance between baseline efficiency and worst-case robustness. They incur only a small efficiency loss under clean data, yet deliver dramatic improvements under the contamination regimes that most severely degrade the standard RF, especially `variance inflation, shift` contamination, and `tail variance-deflation`. Under milder contamination, particularly `central variance-deflation`, the gap narrows and the methods become harder to separate, but even there the hybrids remain broadly competitive.

##### Screening Results

A graphical overview of the entire multi-stage screening procedure, from the initial pool of candidate methods to the breakdown-point evaluation, is provided in Figure 7.

#### Animal (simulated) data analysis

The predictive performance of the standard RF and the two robust alternatives, RF-`k` and RF-`w`, differed only modestly across the three milk traits, but the pattern was trait specific. For trait T1, which underpins the simulation experiments, the standard RF achieved the highest PA (0.755) and the lowest PEs, whereas both robust variants yielded slightly lower PA and somewhat larger RMSPE and MAPE values (Table S20). A similar pattern emerged for T2, although the differences among methods were small. For T3, all three methods performed almost identically, with only negligible differences in PA and error measures. These results accord with expectation: methods designed to dampen the influence of contamination may pay a small efficiency cost when the data are clean. They therefore suggest that, in uncontaminated settings, the advantage of robustness is subtle rather than sweeping, and that its cost, when present, is slight rather than severe.

The recovery of elite genotypes provides a sharper measure of practical performance than aggregate PE alone. Let T denote the set of true top 5% genotypes, defined by the true breeding values in the validation set, and *P* the set of genotypes predicted to fall within the top 10%. Recall is defined as Recall = |*P* ∩ *T* |*/*|*T* | and represents the proportion of truly elite genotypes recovered by the predictions, whereas precision is defined as Precision = |*P* ∩ *T* |*/*|*P* | and represents the proportion of predicted elite genotypes that truly belong to the top 5% (Table 6). Because recall and precision do not describe the remaining individuals selected within the predicted top 10%, we also considered the set of false positives, *FP* = *P* \ *T*, namely, genotypes included in the predicted top 10% but absent from the true top 5%. For each genotype *i* ∈ *FP*, its position in the true ranking was summarized by the corresponding true percentile based on the validation breeding values (Table S21). This framework moves the analysis from general predictive performance to breeding relevance, from error in the abstract to merit in selection.

**Table 6.** Recovery of elite genotypes across traits and methods. Higher values of recall and precision indicate better identification of top-performing genotypes.

| Trait | Method | True 5% | Pred. 10% | Captured | Recall | Precision |
| --- | --- | --- | --- | --- | --- | --- |
| $T_1$ | RF | 51 | 102 | 32 | 0.627 | 0.314 |
|  | RF-**k** | 51 | 102 | 30 | 0.588 | 0.294 |
|  | RF-**w** | 51 | 102 | 26 | 0.510 | 0.255 |
| $T_2$ | RF | 51 | 102 | 34 | 0.667 | 0.333 |
|  | RF-**k** | 51 | 102 | 36 | 0.706 | 0.353 |
|  | RF-**w** | 51 | 102 | 37 | 0.725 | 0.363 |
| $T_3$ | RF | 51 | 102 | 30 | 0.588 | 0.294 |
|  | RF-**k** | 51 | 102 | 30 | 0.588 | 0.294 |
|  | RF-**w** | 51 | 102 | 29 | 0.569 | 0.284 |

Across traits and methods, higher recall and precision consistently marked better identification of top-performing genotypes (Table 6). For trait T1, the standard RF showed the strongest recovery, identifying 32 of the true top 5% genotypes, whereas both preprocessing approaches, RF-`k` and RF-`w`, recovered slightly fewer elite individuals. For trait T2, by contrast, the preprocessing methods outperformed the classical RF, with RF-`w` achieving the highest recall and precision. For trait T3, all methods again performed similarly, with only minor differences in the number of elite genotypes recovered. These results suggest that preprocessing can improve the detection of elite individuals in some settings, but that its benefit is not universal and depends on the trait under consideration. The implication is plain but important: no single method dominates across all traits, and method choice therefore remains conditional on trait-specific structure.

The degree of agreement among the methods in identifying elite genotypes further clarifies how similarly the competing approaches rank the most valuable individuals. The Venn diagrams (Figure 8) show substantial overlap in the elite genotypes captured by the three methods across all traits, indicating that the different modelling strategies converge on largely the same set of superior candidates. The strongest consensus occurs for trait T2, for which 33 genotypes are simultaneously identified by all three approaches, demonstrating a remarkably consistent recognition of top-performing individuals. Agreement is similarly strong for trait T3, where 25 elite genotypes are jointly recovered. Although concordance is somewhat lower for trait T1, most elite genotypes identified by any single method are still recovered by at least one of the others. These patterns indicate that the robust preprocessing strategies preserve the core ranking structure produced by the classical RF, altering only a small subset of marginal candidates rather than reshaping the elite set itself. The alignment plot (Figure S12) reinforces this interpretation by showing that, across all traits, most individuals belonging to the true top 5% are simultaneously recovered by the different methods. This close alignment indicates that robust preprocessing does not disrupt the identification of the most valuable genotypes but instead preserves the essential selection signal detected by the classical RF. In practical breeding terms, the methods therefore agree strongly on which individuals truly merit advancement, even when their overall PEs differ slightly.

**Fig. 8.**
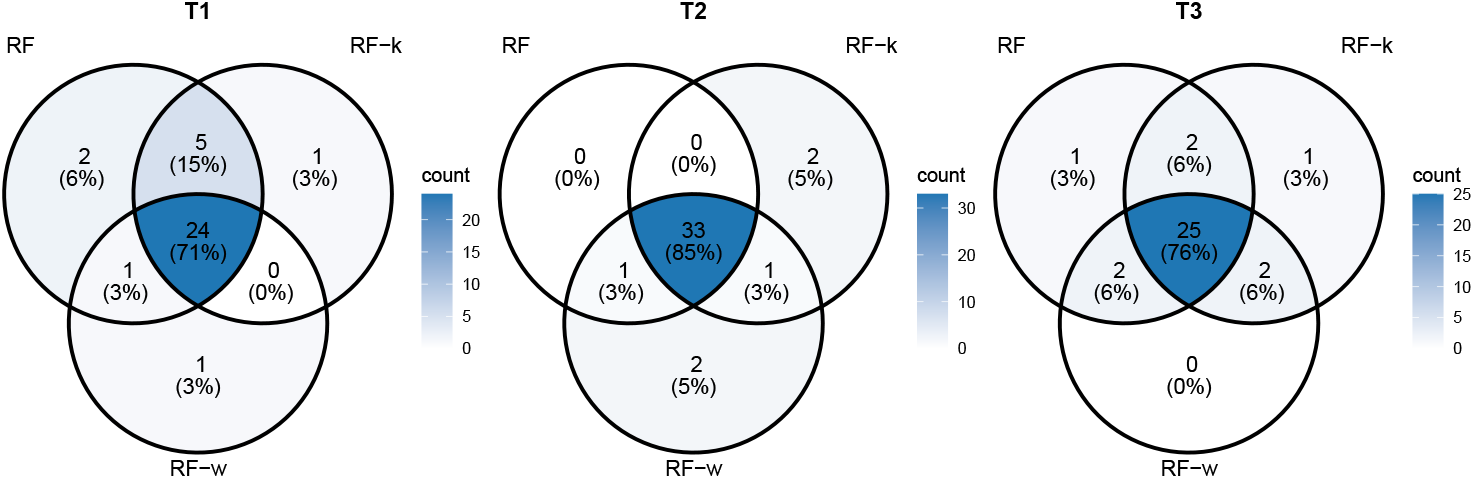
Overlap among the captured true top 5% genotypes across RF, RF-`k`, and RF-`w`. For each trait, the table reports the number of elite genotypes that were jointly identified by pairs of methods and by all three methods.

Agreement among methods was quantified more formally using the Jaccard similarity index, defined as

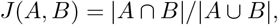

where | · | denotes the cardinality of a set. Values closer to 1 indicate stronger agreement between the predicted elite sets. The resulting Jaccard indices (Figure S13) confirm the visual evidence from the Venn diagrams, revealing substantial overlap in elite genotype selection across the different approaches. These results show that the methods differ mainly in marginal rankings rather than in the identification of the truly superior individuals. Consequently, even when PA varies slightly among approaches, the core biological conclusion remains stable: the principal elite genotypes are consistently identified across methods, demonstrating strong agreement in practical selection outcomes.

The quality of the falsely selected genotypes further clarifies how serious these selection errors actually were. To assess this quality, we examined the false positives (FP), defined as genotypes included in the predicted top 10% but not in the true top 5% (Table S21). Across all traits and methods, these FP were still relatively well ranked in the true ordering, with mean true percentiles ranging from 77.9% to 83.6%. For T1 and T2, most FP fell within the true top 20%, indicating that the misclassified selections were generally close to elite status. For T3, FP were somewhat more dispersed, with a larger share lying in the true 20–50% interval, although only a small fraction belonged to the true bottom 50%. These patterns show that the mistakes were usually near misses rather than wild misfires, a reassuring result for practical selection.

Differences among methods in FP quality were generally small, but a few trait-specific contrasts stood out. For T1, the classical RF produced slightly higher-quality FP on average, although the differences from the robust variants were minor. For T2, the robust RF-`k` approach yielded the most favorable FP profile, combining the highest mean true percentile with no FP in the true bottom 50%. For T3, all methods produced somewhat lower FP quality than for the other traits, but the proportion of clearly poor selections remained limited. Overall, these results indicate that even when a method failed to recover truly elite genotypes, the genotypes selected in error were typically still of reasonably high merit, with the robust variants performing comparably to, and in some cases slightly more conservatively than, the classical RF. Thus, the methods diverged less in the gravity of their mistakes than in the frequency with which they captured the truly best individuals.

Read together, the results from PA, recall, precision, and FP quality show that the main differences among methods lie in their ability to recover truly elite genotypes, not in the overall merit of the remaining selected individuals. The classical RF performed slightly better for T1, the preprocessing variants showed an advantage for T2, and all methods were broadly comparable for T3. Moreover, when errors occurred, the falsely selected genotypes usually still ranked relatively high in the true ordering, indicating that the robust preprocessing strategies produced selections broadly competitive with those of the classical RF. The broader message is therefore both measured and consequential: under clean simulated conditions, robustness does not uniformly improve performance, but neither does it meaningfully degrade selection quality, leaving preprocessing-based robust approaches as credible, competitive, and in some cases advantageous alternatives for genomic selection.

#### Real data analysis

We report only the real-data results for the standard RF, used as the baseline, and the two robust methods, RF-`k` and RF-`w`, which emerged as the strongest candidates in the sequential elimination tests based on PA and PEs (Tables S13-S15, S22; Figures S13-S16). Normality tests conducted before analysis yielded sharply contrasting patterns across datasets and traits. In the maize data, the single analyzed trait, *ear height*, failed the normality test (*p* − *values* ≃ 1.46 × 10^−4^; Table S2, Figure S2). In the soybean data, the first trait, *carbon isotope ratio*, was normally distributed, whereas the second trait, *canopy wilting*, was not (Figure S3). In the wheat data, the average *grain yield* was not normally distributed (Table S2, Figure S4). In the mice data, none of the traits was normally distributed (Table S2, Figure S5). These results show that the real-data benchmark spans both approximately normal and clearly non-normal traits, thereby providing a stringent test of whether robustification improves RF performance under realistic biological variation.

The location of zero in the response distribution provides the key diagnostic for interpreting weighted preprocessing in the real-data analyses (Table S2). Zero lies within the interquartile range only for wheat (GY), and thus falls within the central region of that trait. Consistently, the GY median is negligible relative to the response scale (|*m*|*/s*_*y*_ ≈ 0.01), showing that the median is effectively close to zero on the scale of the data. By contrast, all other traits are either strictly positive or strictly negative; for them, zero lies outside the distributional support and therefore outside the central region. Their medians are also large relative to the response scale, with |*m*|*/s*_*y*_ ranging from approximately 2 to over 100. Thus, shrinkage toward zero is justified only for GY; for the remaining traits zero is a false centre, and the median-centred formulation is therefore the coherent weighting scale.

This centring contrast explains why the two weighted-response formulations behave differently in real data. We compared the uncentred transformation ***ω***_*i*_*y*_*i*_ with the median-centred transformation *m* + ***ω***_*i*_(*y*_*i*_ − *m*), using correlations and pairwise inversion rates to quantify agreement between the original and transformed responses (Table S23), and predictive performance to evaluate model consequences (Tables S22 & S24). These diagnostics show not merely whether weighting changes the response, but how it changes the inferential structure on which RF-`w` is trained. Consistent with the |*m*|*/s*_*y*_ diagnostic, increasing departure from zero centring produced a progressive loss of structural agreement under the uncentred transformation ***ω***_*i*_*y*_*i*_: correlations with the original response decreased, whereas inversion rates increased, indicating poorer preservation of pairwise ordering. This effect is negligible for traits approximately centred at zero (e.g., wheat GY), but becomes pronounced for traits that are strictly positive, strictly negative, or otherwise far from zero. In contrast, the median-centred formulation preserved a stable relationship with the original response across all traits, supporting its use as the general preprocessing strategy (Tables S23-S24). Unless otherwise stated, therefore, the real-data analyses below refer to the median-centred formulation *m* + ***ω***_*i*_(*y*_*i*_ − *m*).

Across all real datasets and most traits, standard RF, RF-`k`, and median-centred RF-`w` produced broadly similar results (Table S22; shrinkage towards the median). Standard RF often achieved slightly higher PAb and slightly lower PEs, but these differences were generally modest and trait-dependent.

The direct comparison between ***ω***_*i*_*y*_*i*_ and *m* + ***ω***_*i*_(*y*_*i*_ − *m*) clarifies that the practical value of weighted preprocessing depends on response centring. For traits approximately centred at zero, the two formulations yielded nearly identical transformations and predictive results (Wheat - GY). For traits far from zero, however, the uncentred transformation induced substantial rank distortion, evident in lower correlations, higher inversion rates, marked deterioration in predictive performance (e.g., Soybean - CIR & Mice - BL). These results identify median-centred RF-`w` as the more stable and biologically portable weighting strategy across traits that differ in support, scale, and distributional shape.

## Discussion and Conclusion

The substantial variability of RF results across random seeds justifies systematic attention to seed dependence when optimizing predictive performance. A practical implication is that one should search for a favourable random seed during model development and then fix that seed for reproducibility. Even so, exact replication may still vary across machines, computing platforms, and software versions. To further reduce seed-driven variation, it is advisable to generate predictions from several replicated fits and average them. This strategy does not eliminate algorithmic randomness, but it does tame it and thus yields more stable inference.

### Simulation setting

The simulation results reveal most clearly when robustification matters and why. When the training data are contaminated and the test data remain clean, robust RF methods reduce the influence of atypical observations, recover the underlying signal more faithfully, and maintain more stable predictive performance across contamination scenarios. Under those conditions, robustification is not a cosmetic adjustment but a functional safeguard, because the predictive target is the uncontaminated signal rather than the contaminated empirical distribution. The standard RF, by contrast, is more exposed to contamination and therefore suffers sharper declines in predictive performance.

The uncontaminated simulation results show the complementary case in which robustness is not needed and may even exact a small cost. When both the training and test data are clean, the standard RF exploits the full structure of the data without having to discount any observations and therefore tends to perform best. In that setting, robust methods can lose some efficiency because they protect against contamination that is absent. The central lesson is therefore conditional rather than absolute: robust RF excels when contamination distorts the training data relative to the predictive target, whereas the standard RF excels when the data are clean and no protection is required.

### Real data setting

The real plant and animal datasets present a more difficult and more biologically realistic setting because the true data-generating mechanism is unknown. The training and test sets are sampled from the same observed data and therefore are likely to share non-normality, outliers, heavy tails, and heterogeneous structure. A crucial consequence follows: the test set is not the truth, but only a proxy for the unknown target, such as the underlying breeding value. Predictive evaluation is therefore conducted against the observed empirical distribution rather than against the unobserved signal that generated it.

This distinction helps explain why the standard RF seemingly performs better on real data. Because it learns the full empirical distribution, including its irregularities, it can align more closely with test data drawn from that same distribution. When unusual values or heterogeneous substructures occur in both training and test sets, fitting them is not necessarily a defect under empirical prediction; indeed, it may improve apparent predictive performance. In such cases, the standard RF benefits from matching the evaluation target, even when that target itself contains distortions.

By contrast, robust RF methods are designed to reduce the influence of atypical observations in the training data. That is an advantage when the analytic goal is to recover the underlying signal, as in the contaminated-versus-clean simulation setting, but it can become a disadvantage when the test data contain the same atypical structures as the training data. Robustification may then attenuate patterns that also occur in the evaluation set and thereby introduce a mismatch between the fitted model and the data used to assess it. This empirical asymmetry explains why robust RF methods may not consistently improve predictive performance in real-data analyses, even when they are conceptually better aligned with recovery of the latent biological signal.

The real-data results therefore argue against a simple verdict either for or against robustification. Superior performance of the standard RF on observed real datasets does not prove that robustness is unnecessary; rather, it shows that empirical prediction and signal recovery are not always the same objective. When breeders seek the best prediction within the observed population structure, the standard RF may often suffice and may indeed be optimal. When, however, there is reason to suspect substantial contamination, misrecording, phenotypic corruption, or a mismatch between observed responses and the latent target of selection, robust RF becomes much more attractive. The choice should thus depend on the inferential aim, the expected contamination structure, and the anticipated relation between training and deployment data.

The ranking- and weighting-based robustification strategies remain especially attractive because they are conceptually simple, flexible, and broadly portable across machine-learning methods. They operate through transformations of the response or through observation-level weighting and therefore do not require altering the core RF learning algorithm. That generality is a practical strength. Yet the results also show that not all robustification strategies are equally reliable. Across both simulations and real data, the ranking-based RF was generally the more stable robust alternative, often preserving much of the predictive performance of the standard RF while offering greater protection under contamination. Its main strength is that it dampens the influence of extreme responses without relying on highly sensitive weighting schemes.

The weighting-based approach remains flexible, but the present analyses show that its value depends chiefly on how the weighting transformation is formulated. In particular, the median-centred formulation produced substantially more stable behaviour across traits than the uncentred transformation, especially for responses far from zero. These findings show that weighted robust RF approaches must account for response support, scale, and centring; they also support shrinkage toward a robust central location as the most coherent general formulation.

The results also clarify what these RF-based models can and cannot say about predictor importance. RF methods can provide relative importance measures for markers or groups of predictors, but contamination can distort those rankings by altering splits, changing local partitions, and magnifying noise-driven structure. Robust RF can therefore improve the stability of importance measures when contamination is substantial, because it reduces the leverage of atypical responses on tree construction or aggregation. Even so, variable importance in these models remains fundamentally relative rather than absolute, and it should be interpreted cautiously, especially in high-dimensional genomic settings with correlated markers. Robustification may sharpen the signal, but it does not convert variable importance into a direct measure of causal effect.

For practical plant and animal breeding, the most defensible recommendation is neither to replace the standard RF wholesale nor to invoke robustification indiscriminately. When the data appear clean, when training and target populations are expected to share the same empirical structure, and when predictive performance under the observed distribution is the main objective, the standard RF should remain the default choice. When contamination is suspected to be substantial, when phenotypes may contain atypical or corrupted observations, or when the goal is to recover a cleaner latent signal for genomic selection, robust RF should be used alongside the standard RF. Among the robust options considered here, RF-`k` and the median-centred RF-`w` emerged as the most dependable alternatives across the simulated contamination scenarios and the real plant and animal datasets, delivering stable, broadly comparable performance across traits.

In conclusion, the results show that robust RF are not universally superior, but they are often necessary when contamination materially distorts the relation between the observed responses and the predictive target. Standard RF excels on clean data and often on real datasets evaluated against their own empirical distribution. Robust RF excels when contamination is substantial and the aim is to recover the underlying signal rather than merely to reproduce the observed distribution. The wisest practice is therefore comparative and context-sensitive: fit the standard RF routinely, pair it with robust alternatives when contamination is plausible, examine rank preservation and weighting behaviour carefully, and let the data and the breeding objective determine the final choice. Used in that way, robust RF methods can become a valuable complement to standard RF and to other machine-learning tools, helping breeders reduce the risks posed by contamination while preserving the gains of accurate GP and reliable selection.

## Funding

This work is funded by national funds through the FCT – Fundação para a Ciência e a Tecnologia, I.P., under the scope of the projects UID/00297/2025 (https://doi.org/10.54499/UID/00297/2025) and UID/PRR/00297/2025 (https://doi.org/10.54499/UID/PRR/00297/2025) (Center for Mathematics and Applications), project 2023.14934.PEX (https://doi.org/10.54499/2023.14934.PEX) and and FCT Mobility Grant

FCT/Mobility/1359099645/2024-25. This work is also supported by computational resources from FCT I.P., approved under the Project 2023.14934.PEX.F1 at Deucalion supercomputer, jointly funded by EuroHPC JU and Portugal. We further acknowledge the EuroHPC Joint Undertaking for awarding access to Deucalion at MACC, Portugal, under project EHPC-DEV-2026D04-013.

## Supplementary material

Supplementary material will be available at Briefings in Bioinformatics online upon publication of the article.

## Data and code availability

The data and R code used in the simulations and real data analyses will be made publicly available upon publication of the article.

## Competing interests

No competing interest is declared.

## Notes

### Competing Interest Statement

The authors have declared no competing interest.

### Summary of Updates

In this revised version of the manuscript, we introduce a refinement to the preprocessing step used in the weighted random forest (RF-w) method. The update concerns how the weighted response is constructed and provides a more general and coherent formulation across different types of response variables. In the original version, the weighting transformation effectively shrank observations toward zero. While appropriate when the response is centred near zero, this behaviour can distort the structure of the data when responses are strictly positive, strictly negative, or otherwise not centred at zero. The revised formulation instead shrinks observations toward a robust central value (the median), ensuring that the transformation respects the location of the response distribution. This change does not affect the conclusions of the simulation study. The simulated data were approximately centred at zero, so both formulations produced nearly identical results. In contrast, the real-data analyses have been updated, as several traits were not centred near zero. The revised formulation leads to modest but more coherent results for the RF-w method across these traits. The manuscript has been updated accordingly. Changes include a refined description of the RF-w preprocessing step, revisions to the real-data analysis and discussion sections, and updated supplementary materials. For the real-data analyses, the supplementary tables now report results for both the original and revised formulations, and an additional figure illustrates their comparative behaviour. Overall, this update should be viewed as a methodological refinement that generalises the original approach. The main findings and conclusions of the study remain unchanged, while the revised formulation ensures more consistent behaviour across a wider range of data settings.

## References

1. Lello L, Avery SG, Tellier L, Vazquez AI, de Los Campos G, Hsu SD. Accurate genomic prediction of human height. Genetics. 2018;210(2):477–497.

2. Bellot P, de Los Campos G, Pérez-Enciso M. Can deep learning improve genomic prediction of complex human traits? Genetics. 2018;210(3):809–819.

3. Crossa J, Beyene Y, Kassa S, Pérez P, Hickey JM, Chen C, et al. Genomic prediction in maize breeding populations with genotyping-by-sequencing. G3: Genes, genomes, genetics. 2013;3(11):1903–1926.

4. Crossa J, Perez P, Hickey J, Burgueno J, Ornella L, Cerón-Rojas J, et al. Genomic prediction in CIMMYT maize and wheat breeding programs. Heredity. 2014;112(1):48–60.

5. Millet EJ, Kruijer W, Coupel-Ledru A, Alvarez Prado S, Cabrera-Bosquet L, Lacube S, et al. Genomic prediction of maize yield across European environmental conditions. Nature genetics. 2019;51(6):952–956.

6. Ogutu JO, Piepho HP, Schulz-Streeck T. A comparison of random forests, boosting and support vector machines for genomic selection. In: BMC proceedings. vol. 5. Springer; 2011. p. S11.

7. Ogutu JO, Schulz-Streeck T, Piepho HP. Genomic selection using regularized linear regression models: ridge regression, lasso, elastic net and their extensions. In: BMC proceedings. vol. 6. Springer; 2012. p. S10.

8. Ogutu JO, Piepho HP. Regularized group regression methods for genomic prediction: Bridge, MCP, SCAD, group bridge, group lasso, sparse group lasso, group MCP and group SCAD. BMC Proceedings. 2014;8(5):1–9.

9. Montesinos-López A, Montesinos-López OA, Gianola D, Crossa J, Hernández-Suárez CM. Multi-environment genomic prediction of plant traits using deep learners with dense architecture. G3: Genes, Genomes, Genetics. 2018;8(12):3813–3828.

10. Montesinos-López OA, Martín-Vallejo J, Crossa J, Gianola D, Hernández-Suárez CM, Montesinos-López A, et al. A benchmarking between deep learning, support vector machine and Bayesian threshold best linear unbiased prediction for predicting ordinal traits in plant breeding. G3: Genes, Genomes, Genetics. 2019;9(2):601–618.

11. Montesinos-López OA, Martín-Vallejo J, Crossa J, Gianola D, Hernández-Suárez CM, Montesinos-López A, et al. New deep learning genomic-based prediction model for multiple traits with binary, ordinal, and continuous phenotypes. G3: Genes, genomes, genetics. 2019;9(5):1545–1556.

12. Pérez-Enciso M, Zingaretti LM. A guide on deep learning for complex trait genomic prediction. Genes. 2019;10(7):553.

13. Chung CW, Hsiao TH, Huang CJ, Chen YJ, Chen HH, Lin CH, et al. Machine learning approaches for the genomic prediction of rheumatoid arthritis and systemic lupus erythematosus. BioData mining. 2021;14(1):52.

14. Lourenço VM, Ogutu JO, Rodrigues RA, Posekany A, Piepho HP. Genomic prediction using machine learning: a comparison of the performance of regularized regression, ensemble, instance-based and deep learning methods on synthetic and empirical data. BMC genomics. 2024;25(1):152.

15. Liano K. Robust error measure for supervised neural network learning with outliers. IEEE Transactions on Neural Networks. 1996;7(1):246–250.

16. Hawkins S, He H, Williams G, Baxter R. Outlier detection using replicator neural networks. In: International conference on data warehousing and knowledge discovery. Springer; 2002. p. 170–180.

17. Beliakov G, Kelarev A, Yearwood J. Robust artificial neural networks and outlier detection. Technical report. arXiv preprint arXiv:11100169. 2011;.

18. Chuang CC, Lee ZJ. Hybrid robust support vector machines for regression with outliers. Applied Soft Computing. 2011;11(1):64–72.

19. Le T, Tran D, Ma W, Pham T, Duong P, Nguyen M. Robust support vector machine. In: 2014 International Joint Conference on Neural Networks (IJCNN). IEEE; 2014. p. 4137–4144.

20. Roy MH, Larocque D. Robustness of random forests for regression. Journal of Nonparametric Statistics. 2012;24(4):993–1006.

21. Xu G, Cao Z, Hu BG, Principe JC. Robust support vector machines based on the rescaled hinge loss function. Pattern Recognition. 2017;63:139–148.

22. Sage A. Random forest robustness, variable importance, and tree aggregation. 2018;.

23. Raymaekers J, Rousseeuw P, Servotte T, Verdonck T, Yao R. A Powerful Random Forest Featuring Linear Extensions (RaFFLE); 2025.

24. Liu H, Shah S, Jiang W. On-line outlier detection and data cleaning. Computers & chemical engineering. 2004;28(9):1635–1647.

25. Van den Broeck J, Argeseanu Cunningham S, Eeckels R, Herbst K. Data cleaning: detecting, diagnosing, and editing data abnormalities. PLoS medicine. 2005;2(10):e267.

26. Sharifnia AM, Kpormegbey DE, Thapa DK, Cleary M. A primer of data cleaning in quantitative research: Handling missing values and outliers. Journal of Advanced Nursing. 2026;82(1):970–975.

27. Montesinos-López OA, Crossa J, Vitale P, Gerard G, Crespo-Herrera L, Dreisigacker S, et al. GBLUP outperforms quantile mapping and outlier detection for enhanced genomic prediction. International Journal of Molecular Sciences. 2025;26(8):3620.

28. Welsch RE. Influence functions and regression diagnostics. In: Modern data analysis. Elsevier; 1982. p. 149–169.

29. Bernardo R. Outliers and their distribution in breeding populations. Crop Science. 2022;62(3):1107–1114.

30. Estaghvirou SBO, Ogutu JO, Piepho HP. Influence of outliers on accuracy estimation in genomic prediction in plant breeding. G3: Genes, Genomes, Genetics. 2014;4(12):2317–2328.

31. Lourenço VM, Rodrigues PC, Pires AM, Piepho HP. A robust DF-REML framework for variance components estimation in genetic studies. Bioinformatics. 2017;33(22):3584–3594.

32. Lourenço VM, Ogutu JO, Piepho HP. Robust estimation of heritability and predictive accuracy in plant breeding: evaluation using simulation and empirical data. BMC genomics. 2020;21(1):43.

33. Usai MG, Gaspa G, Macciotta NP, Carta A, Casu S. XVIth QTLMAS: simulated dataset and comparative analysis of submitted results for QTL mapping and genomic evaluation. In: BMC proceedings. vol. 8. Springer; 2014. p. S1.

34. Kaler AS, Purcell LC, Beissinger T, Gillman JD. Genomic prediction models for traits differing in heritability for soybean, rice, and maize. BMC Plant Biology. 2022;22(1). doi:10.1186/s12870-022-03479-y.

35. QuantGen. GPDatasets; 2023. https://github.com/QuantGen/GPDatasets.

36. Crossa J, Campos Gdl, Pérez P, Gianola D, Burgueno J, Araus JL, et al. Prediction of genetic values of quantitative traits in plant breeding using pedigree and molecular markers. Genetics. 2010;186(2):713–724.

37. Pérez P, de Los Campos G. Genome-wide regression and prediction with the BGLR statistical package. Genetics. 2014;198(2):483–495.

38. de Los Campos G, Perez P, Vazquez AI, Crossa J. Genome-enabled prediction using the BLR (Bayesian Linear Regression) R-package. Genome-wide association studies and genomic prediction. 2013; p. 299–320.

39. Wallace JG, Bradbury PJ, Zhang N, Gibon Y, Stitt M, Buckler ES. Association mapping across numerous traits reveals patterns of functional variation in maize. PLoS genetics. 2014;10(12):e1004845.

40. Song Q, Hyten DL, Jia G, Quigley CV, Fickus EW, Nelson RL, et al. Development and evaluation of SoySNP50K, a high-density genotyping array for soybean. PloS one. 2013;8(1):e54985.

41. Song Q, Hyten DL, Jia G, Quigley CV, Fickus EW, Nelson RL, et al. Fingerprinting soybean germplasm and its utility in genomic research. G3: Genes, genomes, genetics. 2015;5(10):1999–2006.

42. Knox EM, Ng RT. Algorithms for mining distancebased outliers in large datasets. In: Proceedings of the international conference on very large data bases. Citeseer; 1998. p. 392–403.

43. Knorr EM, Ng RT. Finding intensional knowledge of distance-based outliers. In: Vldb. vol. 99; 1999. p. 211–222.

44. Su S, Xiao L, Ruan L, Gu F, Li S, Wang Z, et al. An efficient density-based local outlier detection approach for scattered data. IEEE Access. 2018;7:1006–1020.

45. Ru X, Liu Z, Jiang W. Normalized residual-based outlier detection. In: 2014 IEEE International Conference on Signal Processing, Communications and Computing (ICSPCC). IEEE; 2014. p. 190–193.

46. Huber PJ. Robust Estimation of a Location Parameter. Annals of Mathematical Statistics. 1964;35(1):73–101. doi:10.1214/aoms/1177703732.

47. Huber PJ. Robust estimation of a location parameter. In: Breakthroughs in statistics: Methodology and distribution. Springer; 1992. p. 492–518.

48. Breiman L. Random forests. Machine learning. 2001;45:5–32.

49. Huber PJ. Robust Statistics: A Review. Annals of Mathematical Statistics. 1972;43:1041–1067.

50. Maronna RA, Martin RD, Yohai VJ. Robust Statistics. Chichester: Wiley; 2006.

51. Amado C, Pires AM. Robust bootstrap with non random weights based on the influence function. Communications in Statistics-Simulation and Computation. 2004;33(2):377–396.

52. Salibian-Barrera M, Van Aelst S, Willems G. Fast and robust bootstrap. Statistical Methods and Applications. 2008;17(1):41–71.

53. Amado C, Bianco AM, Boente Boente GL, Pires AM. Robust bootstrap: an alternative to bootstrapping robust estimators. 2014;.

54. Box GE, Cox DR. An analysis of transformations. Journal of the Royal Statistical Society Series B: Statistical Methodology. 1964;26(2):211–243.

55. Sakia RM. The Box-Cox transformation technique: a review. Journal of the Royal Statistical Society: Series D (The Statistician). 1992;41(2):169–178.

56. Carroll RJ, Ruppert D. Transformation and weighting in regression. Chapman and Hall/CRC; 2017.

57. Atkinson AC, Riani M, Corbellini A. The box–cox transformation: Review and extensions. 2021;.

58. Raymaekers J, Rousseeuw PJ. Transforming variables to central normality. Machine Learning. 2024;113(8):4953–4975.

59. Duan N. Smearing estimate: a nonparametric retransformation method. Journal of the American Statistical Association. 1983;78(383):605–610.

60. Taylor JM. The retransformed mean after a fitted power transformation. Journal of the American Statistical Association. 1986;81(393):114–118.

61. Yeo IK, Johnson RA. A new family of power transformations to improve normality or symmetry. Biometrika. 2000;87(4):954–959.

62. Dixon WJ, Tukey JW. Approximate behavior of the distribution of Winsorized t (Trimming/Winsorization 2). Technometrics. 1968;10(1):83–98.

63. Dixon WJ, Yuen KK. Trimming and winsorization: A review. Statistische Hefte. 1974;15(2):157–170.

64. Yale C, Forsythe AB. Winsorized regression. Technometrics. 1976;18(3):291–300.

65. Conover WJ, Iman RL. Rank Transformations as a Bridge Between Parametric and Nonparametric Statistics. The American Statistician. 1981;35:121–129.

66. Conover WJ. The rank transformation—an easy and intuitive way to connect many nonparametric methods to their parametric counterparts for seamless teaching introductory statistics courses. Wiley Interdisciplinary Reviews: Computational Statistics. 2012;4(5):432–438.

67. Meinshausen N, Ridgeway G. Quantile regression forests. Journal of machine learning research. 2006;7(6).

68. Azodi CB, Bolger E, McCarren A, Roantree M, de Los Campos G, Shiu SH. Benchmarking parametric and machine learning models for genomic prediction of complex traits. G3: Genes, Genomes, Genetics. 2019;9(11):3691–3702.

69. Rousseeuw PJ, Van Zomeren BC. Unmasking multivariate outliers and leverage points. Journal of the American Statistical association. 1990;85(411):633–639.

70. Chiang JT, et al. The masking and swamping effects using the planted mean-shift outliers models. Int J Contemp Math Sciences. 2007;2(7):297–307.

71. Serfling R, Wang S. General foundations for studying masking and swamping robustness of outlier identifiers. Statistical Methodology. 2014;20:79–90.

