## Supplementary material for "Robust Random Forests for Genomic Prediction: Challenges and Remedies"

### Supplementary Materials

### Methods

### Details of model fitting

#### Computational resources and software

All computations were performed using the R statistical computing environment (R Core Team). The analyses were executed on the Deucalion high-performance computing (HPC) system, a EuroHPC petascale supercomputer hosted in Portugal. The simulations and model-fitting procedures were run in a containerized R environment based on the `rocker/rstudio` image (R version 4.5.1), ensuring reproducibility of the software setup across computational runs. Parallel execution on the HPC system was used to reduce the computational burden associated with the simulation study and the robust random forest procedures considered.

Random forest models were implemented using the `ranger` package, except for models employing the least absolute deviation (LAD) split criterion, which were fitted using the `randomForestSRC` package (approximation using pinball loss function with $\tau = 0.5$). The use of the pinball loss provides a principled way to approximate the LAD-based splitting within a tree-based framework. The pinball loss is the standard objective for quantile regression, and for a quantile level $\tau \in (0, 1)$ it is defined as

$$L_\tau(y, \hat{q}) = \begin{cases} \tau(y - \hat{q}), & \text{if } y \geq \hat{q}, \\ (1 - \tau)(\hat{q} - y), & \text{if } y < \hat{q}. \end{cases}$$

This asymmetric loss is minimized by the conditional $\tau$–quantile. In the special case $\tau = 0.5$, the loss becomes symmetric and reduces to a scaled version of the least absolute deviation (LAD) loss,

$$L_{0.5}(y, \hat{q}) = 0.5\,|y - \hat{q}| \propto |y - \hat{q}|.$$

Thus, LAD regression corresponds to quantile regression with $\tau = 0.5$. Using pinball loss with $\tau = 0.5$ in the splitting rule therefore encourages splits that reduce median absolute deviation rather than variance, providing a computationally tractable approximation to the behavior of a true LAD impurity criterion in random forests.

Robust pre-processing procedures relied on the `MASS`, `cellWise`, and `statar` packages. Parallel execution of the simulation study was handled using the `foreach`, `parallel` and `doParallel` packages in order to reduce the computational burden associated with the large number of simulation runs across contamination scenarios and methods, combined with the cost of fitting random forest models with a substantial number of trees.

To ensure reproducibility and comparability across methods, the contaminated datasets were generated once and kept fixed across all simulation runs. The datasets used in the simulations are provided together with the replication materials accompanying this paper, ensuring full reproducibility independently of the random-number generator, software version, or computational platform.

#### RF hyperparameter tuning and other specifics

##### Simulation data

Random forest (RF) models were fitted using the hyperparameter configuration previously tuned in Lourenço et al. (2024). Specifically, the number of trees was fixed to $n_{\text{trees}} = 1000$, the number of candidate variables at each split was set to $mtry = p/6$, where $p$ is the total number of features (covariates) and the minimum node size was fixed at 1. Under this configuration, the reported predictive accuracy (PA) for the trait under study (Trait 1) was 0.741, with corresponding MSPE and MAPE values of 7945.8 (RMSPE $\approx 89.1$) and 72.4, respectively.

The objective of this study is not to further improve the predictive performance of the standard random forests but rather to investigate how its predictive performance degrades across a range of simulated contamination scenarios. Accordingly, no additional hyperparameter tuning was performed, and the previously validated configuration was retained throughout all the experiments, including those involving contaminated data. In addition to the substantial computational burden that would be incurred by re-tuning the RF model for each simulation run and contamination scenario, retaining a fixed hyperparameter configuration ensures that observed changes in performance can be attributed solely to data contamination rather than to model re-optimization as well. Moreover, re-tuning the random forest hyperparameters for contaminated datasets would not be expected to improve predictive performance beyond that obtained using the configuration tuned under uncontaminated conditions, as contamination-induced degradation cannot be mitigated through hyperparameter adjustment alone.

Nevertheless, because random forest predictions depend on stochastic elements introduced by random initialization, we conducted an auxiliary seed search under the uncontaminated setting. This step was not intended as a form of model optimization, but rather to provide a stronger and more stable baseline from which performance degradation under contamination could be evaluated. The distribution of predictive performance across 1000 random number seeds (from 667 to 1666) is shown in Figure S7(a), illustrating the inherent variability induced solely by random initialization (PA$= 0.742$ ($sd = 0.004$)).

Based on this analysis, the seed that performed the best was selected, yielding a predictive accuracy of 0.755 and predictive errors RMSE$= 88.6$ and MAPE$= 72.1$, resulting, relative to the mean values, in an increase in PA of approximately 1.8%, and decrease in RMSPE and MAPE of approximately 0.56% and 0.55%, respectively. The associated performance measures are reported in Table S4. All subsequent contamination experiments were conducted using this fixed seed, ensuring that any observed loss in performance reflects data contamination rather than stochastic variability.

Although performance can vary between pre-processing and algorithmic strategies, conducting a complete seed search for each method would be computationally infeasible. We therefore retain the seed identified from fitting the standard RF model to the uncontaminated data throughout all simulations, providing a consistent and reproducible basis for comparison.

For completeness, Figure S7b-d summarize the distribution of predictive performance in the 1000-seed search for the weighting (RF-`w`), winsorization (RF-`win`) and rank (RF-`k`) strategies, with the corresponding best seeds and model accuracies reported in Table S4, illustrating how PA fluctuates across random initialisations.

The distribution of PA values is approximately normal for all methods according to the Shapiro–Francia test. While differences in mean PA across methods are small, RF shows slightly higher values (0.742) than RF-`win` (0.737), with RF-`k` and RF-`-w` yielding similar performance.

Because the animal response used to calibrate the simulations is effectively centred at zero, the weighted preprocessing step was implemented in the reduced form $w_i = \boldsymbol{\omega}_i y_i$. This implementation follows from the distributional features of Trait T1 in the animal dataset (Table S1): zero falls within the central mass of the observations, as indicated by the interquartile range, and the median is trivial relative to the response scale ($|m|/s \approx 0.01$). Consequently, the median-centred transformation, $m + \boldsymbol{\omega}_i(y_i - m)$, differs from $\boldsymbol{\omega}_i y_i$ only by the small residual term $(1 - \boldsymbol{\omega}_i)m$; with $m$ close to zero, the two transformations are practically indistinguishable. The reduced expression therefore preserves the intended weighting while avoiding unnecessary algebraic clutter, providing an adequate and transparent preprocessing rule for the simulation study.

##### Animal milk traits $T_1$, $T_2$ and $T_3$

The same procedure used for trait $T_1$ in the simulation study was also applied to traits $T_2$ and $T_3$. While the seed for $T_1$ (1019) was fixed from the simulation setup, the seed search for the remaining traits yielded 805 for $T_2$ and 834 for $T_3$. Random forest (RF) models were fitted using the same hyperparameter configuration as that adopted for $T_1$, as previously tuned in Lourenço et al. (2024).

##### Real data

For all species (maize, soybean, wheat and mice), hyperparameter tuning of the RF models was performed on the 80% training portion of each 80–20 split, using a fixed random seed to ensure full reproducibility across traits and datasets. Tuning was conducted via 5-fold cross-validation over the grid $mtry \in \{p/3, p/6, p/10\}$ and $min.node.size \in \{1, 3, 5, 10\}$, while the number of trees was fixed at $ntrees = 100$ for the maize and soybean data, 1000 for the wheat data and 250 for the mice data. These values were chosen to ensure the computational feasibility of the cross-validation procedure given the differing sizes of the SNP matrices and the numbers of genotypes or individuals in the datasets.

In contrast to the simulation study, where the exploration of different random seeds was intentional and scientifically justified because the objective was to characterize seed driven variability, we did not conduct a random-seed search for the real datasets (wheat, maize, and mice). Instead, given the non-negligible seed-to-seed variability observed in the simulations, a more principled strategy for real data analysis is to fit the tuned RF model under several fixed seeds and average the resulting predictions. This approach stabilizes performance estimates while avoiding potential seed-based overfitting.

Consistent with the criteria outlined in the main text, the analyses were restricted to quantitative traits with complete observations, and marker-level missingness was assessed before model fitting. For the mice dataset, only 3 of the 21 quantitative traits were retained for analysis—body mass index, body length, and final normalized body weight—because they were the only traits without missing values. A similar constraint applied to the maize dataset, for which only ear height satisfied this requirement and was therefore retained.

For the genomic data, the wheat, soybean, and mice datasets exhibited negligible SNP missingness and required no filtering before analysis. By contrast, the maize dataset displayed substantial marker-level missingness; consequently, SNPs with more than 20% missing values, representing about 7.15% of the markers, were removed. The remaining SNPs exhibited at most 19.8% missingness and were used in all subsequent analyses.

The final real-data step evaluated both response-weighting transformations, $\omega_i y_i$ and $m + \omega_i(y_i - m)$, so that RF performance could be compared under two distinct shrinkage targets: zero and a robust central location. This paired evaluation made the consequences of each transformation explicit in predictive ability, response-correlation structure, and inversion rates between transformed and original responses.

### Tables

Table S1: Summary statistics for the training and testing datasets for the three simulated animal quantitative traits. The last column reports the p-value of the Shapiro–Francia normality test.

| **Trait** | **Dataset** | **Min** | **Q1** | **Median** | **Mean** | **Q3** | **Max** | **SD** | **SF** $p - value$ |
|---|---|---|---|---|---|---|---|---|---|
| $T_1$ | Training | -584.99 | -116.24 | -1.71 | 0.00 | 112.25 | 587.19 | 176.52 | $5.30 \times 10^{-2}$ |
| | Testing | -325.04 | -59.65 | 16.88 | 14.70 | 87.98 | 356.29 | 107.71 | $5.57 \times 10^{-1}$ |
| $T_2$ | Training | -32.23 | -6.50 | 0.08 | 0.00 | 6.62 | 32.51 | 9.51 | $8.15 \times 10^{-1}$ |
| | Testing | -25.30 | -11.11 | -6.76 | -6.52 | -2.30 | 12.55 | 6.12 | $1.15 \times 10^{-2}$ |
| $T_3$ | Training | -0.10 | -0.02 | 0.00 | 0.00 | 0.02 | 0.09 | 0.02 | $5.75 \times 10^{-2}$ |
| | Testing | -0.08 | -0.04 | -0.03 | -0.03 | -0.01 | 0.02 | 0.02 | $4.33 \times 10^{-3}$ |

Table S2: Summary statistics for the maize (EH: ear height), soybean (CIR: *carbon isotope ratio*; CW: *canopy wilting*), wheat (GY: *grain yield*), and mice (BMI: body mass index; BL: body length; and FNBW: final normalized body weight) quantitative traits based on the full datasets. The last column reports the $p$-value for the Shapiro–Francia normality test.

| **Trait** | **†p** | **†n** | **Min** | **Q1** | **Median** | **Mean** | **Q3** | **Max** | **†SD** | **SF $p$-value** |
|---|---|---|---|---|---|---|---|---|---|---|
| *Maize* | | | | | | | | | | |
| EH | 47,259 | 278 | 8 | 47.60 | 60.25 | 61.47 | 72.50 | 136 | 20.25 | $1.46 \times 10^{-4}$ |
| *Soybean* | | | | | | | | | | |
| CIR | 31,260 | 346 | -29.82 | -29.22 | -29.05 | -29.06 | -28.87 | -28.37 | 0.273 | $6.26 \times 10^{-1}$ |
| CW | 31,260 | 346 | 7.50 | 12.50 | 15.62 | 16.99 | 20.00 | 45.62 | 6.464 | $1.22 \times 10^{-12}$ |
| *Wheat* | | | | | | | | | | |
| GY | 1,279 | 599 | -2.215 | -0.391 | 0.009 | 0.000 | 0.424 | 1.618 | 0.625 | $2.08 \times 10^{-3}$ |
| *Mice* | | | | | | | | | | |
| BMI | 10,346 | 1814 | -0.626 | -0.500 | -0.459 | -0.457 | -0.417 | -0.271 | 0.060 | $2.64 \times 10^{-3}$ |
| BL | 10,346 | 1814 | 5.900 | 7.200 | 7.600 | 7.597 | 8.000 | 9.300 | 0.564 | $7.45 \times 10^{-6}$ |
| FNBW | 10,346 | 1814 | 12.200 | 20.625 | 23.750 | 23.997 | 27.075 | 39.100 | 4.191 | $1.46 \times 10^{-10}$ |

† $p$: number of covariates/features; $n$: number of observations/samples; SD: standard deviation.

Table S3: Pre-processing, algorithm-based, and hybrid strategies evaluated in the first simulation stage, including notation for each method.

| **Notation** | **Method** | |
|---|---|---|
| RF | Standard Random Forests model. | |
| **Preprocessing-based approaches** | | |
| RF-k (i) | Rank transformation | (back-tranformation via linear interpolation) |
| RF-k (ii) | Rank transformation | (back-tranformation via linear interpolation + robust central correction) |
| RF-win (i) | Winsorisation | |
| RF-win (ii) | Winsorisation with the median | |
| RF-BC (i) | Box–Cox transformation | |
| RF-BC (ii) | Box–Cox transformation | (robust central correction of back-transformed predictions) |
| RF-YJ (i) | Yeo–Johnson transformation | |
| RF-YJ (ii) | Yeo–Johnson transformation | (robust central correction of back-transformed predictions) |
| RF-rBC (i) | Robust Box-Cox transformation | |
| RF-rBC (ii) | Robust Box-Cox transformation | (robust central correction of back-transformed predictions) |
| RF-rYJ (i) | Robust Yeo–Johnson transformation | |
| RF-rYJ (ii) | Robust Yeo–Johnson transformation | (robust central correction of back-transformed predictions) |
| RF-w | Robust weighting | |
| **Algorithm-based approaches** | | |
| RF-rboot | Robust boostrapping | |
| RF-lad | LAD loss impurity (median aggregation is default) | |
| RF-m | Median aggregation of predictions of the ensemble of trees | |
| RF-q | Quantile regression aggregation (conditional median) | |
| **Hybrid Strategies** | | |
| RF-k-m (ii) | Combination of rank transformation with median aggregation | |
| RF-k-q (ii) | Combination of rank transformation with LAD impurity (median aggregation is default) | |
| RF-w-m | Combination of robust weighting with median aggregation | |
| RF-w-q | Combination of robust weighting with quantile regression | |

Table S4: Random Forests performance (PA: predictive accuracy; RMSPE: mean squared prediction error; and MAPE: mean absolute error) across the preprocessing strategies for the best four random seeds identified by searching over 1000 random number seeds. Blue values refer to the performance of the robust strategies when the best seed obtained for the standard Random Forests is used. Additional methodological details relevant to interpreting this table are provided in Section *RF hyperparameter tuning and other specifics*, Subsection *Simulation data* at the beginning of the Supplementary Materials.

| Method | Iteration | Seed | PA | RMSPE | MAPE |
|---|---|---|---|---|---|
| RF | 353 | 1019 | 0.755 | 88.61 | 72.13 |
| RF-`w` | 785 | 1451 | 0.744 | 93.02 | 75.44 |
| | | 1019 | 0.731 | 92.98 | 75.37 |
| RF-`k` | 169 | 835 | 0.742 | 94.86 | 76.78 |
| | | 1019 | 0.739 | 95.10 | 77.21 |
| RF-`win` | 994 | 1660 | 0.750 | 90.63 | 73.70 |
| | | 1019 | 0.747 | 90.56 | 73.65 |

Table S5: Estimated predictive accuracy (PA), mean squared prediction error (RMSPE) and mean absolute error (MAPE) for the non-contaminated and `shift` contaminated scenarios — Preprocessing-based Approaches. *sd* is the standard deviation. **Stage 1**

| Method | | | Contamination Level | | | | | | | | | | *sd* |
|---|---|---|---|---|---|---|---|---|---|---|---|---|---|
| | | | 0% | 2% | | | 5% | | | 10% | | | |
| | | | | $k = 5$ | $k = 7$ | $k = 9$ | $k = 5$ | $k = 7$ | $k = 9$ | $k = 5$ | $k = 7$ | $k = 9$ | range |
| RF | | PA | 0.755 | 0.683 | 0.610 | 0.546 | 0.636 | 0.532 | 0.456 | 0.609 | 0.499 | 0.430 | 0.012–0.044 |
| | | RMSPE | 88.61 | 85.62 | 87.83 | 91.71 | 89.88 | 103.74 | 121.68 | 109.11 | 144.85 | 183.71 | 0.78–5.50 |
| | | MAPE | 72.13 | 68.90 | 70.08 | 72.48 | 71.24 | 81.59 | 94.14 | 87.20 | 119.36 | 155.69 | 0.74–5.31 |
| **Preprocessing-based Approaches** | | | | | | | | | | | | | |
| RF-`k` | | PA | 0.739 | 0.734 | 0.733 | 0.734 | 0.728 | 0.725 | 0.728 | 0.723 | 0.718 | 0.718 | 0.005–0.006 |
| | (i) | RMSPE | 95.10 | 94.76 | 94.82 | 94.66 | 94.74 | 94.75 | 94.61 | 93.92 | 94.17 | 94.17 | 0.16–0.38 |
| | | MAPE | 77.21 | 76.88 | 76.93 | 76.78 | 76.81 | 76.91 | 76.78 | 76.17 | 76.32 | 76.32 | 0.11–0.34 |
| | (ii) | RMSPE | 93.93 | 93.62 | 93.67 | 93.51 | 93.68 | 93.69 | 93.51 | 92.97 | 93.20 | 93.17 | 0.17–0.45 |
| | | MAPE | 76.11 | 75.82 | 75.87 | 75.70 | 75.82 | 75.91 | 75.77 | 75.26 | 75.37 | 75.38 | 0.13–0.34 |
| RF-`win` | | PA | 0.747 | 0.739 | 0.735 | 0.738 | 0.712 | 0.702 | 0.706 | 0.626 | 0.507 | 0.433 | 0.004–0.025 |
| | (i) | RMSPE | 90.56 | 88.61 | 88.81 | 88.64 | 86.87 | 86.94 | 86.63 | 106.31 | 141.16 | 179.35 | 0.20–5.38 |
| | | MAPE | 73.65 | 71.86 | 72.01 | 71.87 | 69.53 | 69.49 | 69.14 | 84.96 | 116.18 | 152.03 | 0.24–5.46 |
| | | PA | 0.685 | 0.694 | 0.696 | 0.695 | 0.710 | 0.711 | 0.709 | 0.615 | 0.480 | 0.377 | 0.005–0.038 |
| | (ii) | RMSPE | 97.41 | 95.60 | 95.53 | 95.58 | 91.48 | 91.56 | 91.66 | 95.18 | 109.07 | 127.74 | 0.10–4.40 |
| | | MAPE | 78.84 | 77.43 | 77.30 | 77.39 | 73.92 | 73.98 | 74.04 | 75.43 | 85.86 | 98.82 | 0.12–3.25 |
| RF-`BC` | | PA | 0.747 | 0.731 | 0.713 | 0.702 | 0.712 | 0.690 | 0.674 | 0.703 | 0.665 | 0.646 | 0.006–0.018 |
| | (i) | RMSPE | 90.30 | 88.85 | 88.99 | 88.48 | 86.09 | 84.43 | 83.76 | 82.72 | 85.96 | 90.37 | 0.55–1.32 |
| | | MAPE | 73.58 | 72.22 | 72.28 | 71.80 | 69.69 | 68.28 | 67.37 | 66.26 | 68.37 | 71.81 | 0.49–1.27 |
| | (ii) | RMSPE | 87.24 | 85.84 | 85.68 | 85.10 | 85.71 | 85.02 | 84.65 | 83.77 | 84.34 | 84.52 | 0.67–1.36 |
| | | MAPE | 70.77 | 69.61 | 69.46 | 68.94 | 69.34 | 68.86 | 68.31 | 67.66 | 68.04 | 68.06 | 0.52–1.26 |
| RF-`YJ` | | PA | 0.755 | 0.730 | 0.718 | 0.709 | 0.715 | 0.702 | 0.694 | 0.708 | 0.677 | 0.672 | 0.005–0.18 |
| | (i) | RMSPE | 89.12 | 88.96 | 89.56 | 89.80 | 86.53 | 85.15 | 84.35 | 82.46 | 83.47 | 84.01 | 0.51–1.21 |
| | | MAPE | 72.52 | 72.30 | 72.82 | 72.90 | 70.15 | 68.96 | 68.18 | 66.33 | 66.83 | 67.06 | 0.52–1.12 |
| | (ii) | RMSPE | 86.90 | 85.84 | 85.51 | 85.03 | 85.55 | 84.72 | 84.08 | 83.68 | 84.04 | 83.48 | 0.70–1.36 |
| | | MAPE | 70.50 | 69.62 | 69.39 | 68.94 | 69.24 | 68.56 | 67.94 | 67.70 | 67.84 | 67.25 | 0.55–1.14 |
| RF-`rBC` | | PA | 0.750 | 0.705 | 0.651 | 0.596 | 0.674 | 0.592 | 0.526 | 0.679 | 0.564 | 0.503 | 0.009–0.032 |
| | (i) | RMSPE | 90.30 | 85.70 | 86.03 | 87.61 | 85.88 | 92.73 | 103.49 | 88.27 | 117.82 | 144.44 | 0.71–4.51 |
| | | MAPE | 73.58 | 69.37 | 69.24 | 70.14 | 68.61 | 73.50 | 80.57 | 70.01 | 94.75 | 118.54 | 0.56–4.12 |
| | (ii) | RMSPE | 87.28 | 86.24 | 86.98 | 88.21 | 86.48 | 88.20 | 92.45 | 84.59 | 89.54 | 95.07 | 0.75–2.59 |
| | | MAPE | 70.94 | 69.93 | 70.37 | 71.19 | 69.87 | 71.09 | 73.16 | 68.25 | 71.71 | 75.62 | 0.68–1.82 |
| RF-`rYJ` | | PA | 0.757 | 0.705 | 0.650 | 0.598 | 0.675 | 0.592 | 0.523 | 0.677 | 0.565 | 0.506 | 0.011–0.032 |
| | (i) | RMSPE | 89.08 | 85.61 | 86.03 | 87.50 | 85.81 | 92.71 | 103.72 | 88.34 | 117.81 | 144.56 | 0.81–4.70 |
| | | MAPE | 72.57 | 69.38 | 69.20 | 69.97 | 68.49 | 73.36 | 80.82 | 70.18 | 94.82 | 118.87 | 0.72–4.39 |
| | (ii) | RMSPE | 86.88 | 86.20 | 87.01 | 88.05 | 86.47 | 88.20 | 92.60 | 84.74 | 89.52 | 94.79 | 0.80–2.60 |
| | | MAPE | 70.57 | 69.60 | 70.35 | 70.94 | 69.86 | 71.01 | 73.28 | 68.43 | 71.69 | 75.51 | 0.74–1.79 |
| | | PA | 0.731 | 0.733 | 0.732 | 0.734 | 0.729 | 0.733 | 0.732 | 0.724 | 0.722 | 0.723 | 0.003–0.008 |
| RF-`w` | | RMSPE | 92.98 | 92.21 | 92.26 | 92.07 | 91.00 | 90.90 | 90.92 | 88.83 | 89.03 | 88.91 | 0.18–0.42 |
| | | MAPE | 75.37 | 74.86 | 74.82 | 74.69 | 73.85 | 73.67 | 73.67 | 71.68 | 71.78 | 71.72 | 0.17–0.36 |

**Competing methods:** ■ Highest PAs; ■ Smallest PEs; ■ Smallest PAs; ■ Highest PEs.

Table S6: Estimated predictive accuracy (PA), mean squared prediction error (RMSPE) and mean absolute error (MAPE) for the non-contaminated and `shift` contaminated scenarios — Algorithm-based approaches. *sd* is the standard deviation. **Stage 1**

| Method | | Contamination Level | | | | | | | | | | *sd* |
|---|---|---|---|---|---|---|---|---|---|---|---|---|
| | | 0% | 2% | | | 5% | | | 10% | | | range |
| | | | $k=5$ | $k=7$ | $k=9$ | $k=5$ | $k=7$ | $k=9$ | $k=5$ | $k=7$ | $k=9$ | |
| RF | PA | 0.755 | 0.683 | 0.610 | 0.546 | 0.636 | 0.532 | 0.456 | 0.609 | 0.499 | 0.430 | 0.012–0.044 |
| | RMSPE | 88.61 | 85.62 | 87.83 | 91.71 | 89.88 | 103.74 | 121.68 | 109.11 | 144.85 | 183.71 | 0.78–5.50 |
| | MAPE | 72.13 | 68.90 | 70.08 | 72.48 | 71.24 | 81.59 | 94.14 | 87.20 | 119.36 | 155.69 | 0.74–5.31 |
| **Algorithm-based approaches** | | | | | | | | | | | | |
| RF-`rboot` | PA | 0.737 | 0.724 | 0.711 | 0.704 | 0.701 | 0.662 | 0.640 | 0.670 | 0.596 | 0.545 | 0.006–0.020 |
| | RMSPE | 90.95 | 88.54 | 88.50 | 88.53 | 86.68 | 86.90 | 87.27 | 87.17 | 90.84 | 94.05 | 0.26–0.98 |
| | MAPE | 73.96 | 71.71 | 71.71 | 71.74 | 69.91 | 69.86 | 70.01 | 69.26 | 72.03 | 74.20 | 0.25–0.77 |
| RF-`lad` | PA | 0.637 | 0.588 | 0.588 | 0.575 | 0.572 | 0.541 | 0.506 | 0.552 | 0.520 | 0.491 | 0.020–0.044 |
| | RMSPE | 99.15 | 97.36 | 96.82 | 96.83 | 98.53 | 103.26 | 110.13 | 110.90 | 131.45 | 156.92 | 0.58–2.00 |
| | MAPE | 80.43 | 78.52 | 77.91 | 77.68 | 78.73 | 82.21 | 87.57 | 88.66 | 107.65 | 133.20 | 0.41–2.00 |
| RF-`m` | PA | 0.738 | 0.718 | 0.705 | 0.703 | 0.701 | 0.679 | 0.664 | 0.675 | 0.639 | 0.625 | 0.004–0.017 |
| | RMSPE | 88.87 | 88.67 | 89.09 | 88.84 | 88.59 | 89.18 | 89.30 | 88.01 | 89.31 | 89.97 | 0.40–0.70 |
| | MAPE | 72.22 | 71.96 | 72.36 | 72.02 | 71.89 | 72.20 | 72.25 | 71.14 | 72.15 | 72.54 | 0.45–0.81 |
| RF-`q` | PA | 0.730 | 0.710 | 0.697 | 0.697 | 0.696 | 0.684 | 0.667 | 0.678 | 0.653 | 0.645 | 0.007–0.018 |
| | RMSPE | 91.22 | 91.22 | 91.62 | 91.35 | 91.39 | 91.82 | 9191 | 90.96 | 91.94 | 92.41 | 0.53–0.84 |
| | MAPE | 74.30 | 74.06 | 74.35 | 74.07 | 74.19 | 74.48 | 74.47 | 73.68 | 74.38 | 74.84 | 0.45–0.70 |

**Competing methods:** ■ Highest PAs; ■ Smallest PEs; ■ Smallest PAs; ■ Highest PEs.

Table S7: Estimated predictive accuracy (PA), mean squared prediction error (RMSPE) and mean absolute error (MAPE) for the non-contaminated and `variance-inflated` contaminated scenarios — Preprocessing- and Algorithm-based approaches. *sd* is the standard deviation. **Stage 2**

| Method | | Contamination Level | | | | | | | *sd* |
|---|---|---|---|---|---|---|---|---|---|
| | | 0% | 2% | | 5% | | 10% | | range |
| | | | $s=5$ | $s=7$ | $s=5$ | $s=7$ | $s=5$ | $s=7$ | |
| RF | PA | 0.755 | 0.661 | 0.571 | 0.536 | 0.409 | 0.429 | 0.284 | 0.04–0.08 |
| | RMSPE | 88.61 | 91.96 | 95.29 | 97.77 | 105.49 | 105.33 | 120.88 | 1.91–11.21 |
| | MAPE | 72.13 | 74.34 | 75.95 | 78.17 | 81.95 | 84.00 | 92.22 | 1.33–6.28 |

**a) Preprocessing-based Approaches**

| Method | | 0% | 2% $s=5$ | 2% $s=7$ | 5% $s=5$ | 5% $s=7$ | 10% $s=5$ | 10% $s=7$ | *sd* range |
|---|---|---|---|---|---|---|---|---|---|
| RF-`k` | PA | 0.739 | 0.731 | 0.729 | 0.713 | 0.710 | 0.692 | 0.688 | 0.008–0.014 |
| (i) | RMSPE | 95.10 | 96.49 | 96.60 | 99.46 | 99.49 | 104.46 | 104.54 | 0.26–0.95 |
| | MAPE | 77.21 | 78.31 | 76.93 | 79.27 | 79.26 | 84.77 | 84.86 | 0.21–0.74 |
| (ii) | RMSPE | 93.93 | 95.12 | 95.22 | 97.94 | 97.98 | 102.86 | 102.96 | 0.28–0.98 |
| | MAPE | 76.11 | 77.10 | 77.16 | 79.45 | 79.46 | 83.47 | 83.55 | 0.20–0.78 |
| RF-`BC` | PA | 0.747 | 0.666 | 0.592 | 0.551 | 0.426 | 0.435 | 0.291 | 0.051–0.080 |
| (i) | RMSPE | 89.95 | 90.61 | 92.95 | 95.67 | 102.42 | 102.56 | 117.08 | 1.41–7.51 |
| | MAPE | 73.21 | 73.24 | 74.29 | 76.62 | 79.91 | 81.91 | 89.62 | 0.88-4.12 |
| (ii) | RMSPE | 87.24 | 89.60 | 91.64 | 95.11 | 101.79 | 103.12 | 117.36 | 1.48–6.77 |
| | MAPE | 70.77 | 72.37 | 73.11 | 76.12 | 79.35 | 83.43 | 89.89 | 0.89–3.45 |
| RF-`YJ` | PA | 0.755 | 0.660 | 0.581 | 0.535 | 0.407 | 0.426 | 0.283 | 0.047—0.078 |
| (i) | RMSPE | 89.12 | 91.47 | 93.75 | 98.33 | 105.51 | 107.09 | 121.88 | 1.7–8.31 |
| | MAPE | 72.52 | 73.90 | 74.82 | 78.67 | 82.04 | 85.34 | 93.21 | 1.26–4.04 |
| (ii) | RMSPE | 86.90 | 89.80 | 92.35 | 96.10 | 103.38 | 104.24 | 119.21 | 1.35–8.90 |
| | MAPE | 70.50 | 72.47 | 73.60 | 76.73 | 80.13 | 82.87 | 90.75 | 0.84–4.16 |
| RF-`w` | PA | 0.731 | 0.729 | 0.729 | 0.714 | 0.715 | 0.702 | 0.698 | 0.005–0.012 |
| | RMSPE | 92.98 | 94.12 | 94.25 | 96.37 | 96.21 | 99.33 | 99.32 | 0.28–1.04 |
| | MAPE | 75.37 | 76.45 | 76.50 | 78.20 | 78.08 | 80.64 | 80.62 | 0.19–0.83 |

**Competing methods:** ■ Highest PAs; ■ Smallest PEs; ■ Smallest PAs; ■ Highest PEs.

**b) Algorithm-based approaches**

| Method | | 0% | 2% $s=5$ | 2% $s=7$ | 5% $s=5$ | 5% $s=7$ | 10% $s=5$ | 10% $s=7$ | *sd* range |
|---|---|---|---|---|---|---|---|---|---|
| RF-`m` | PA | 0.738 | 0.718 | 0.711 | 0.688 | 0.676 | 0.648 | 0.614 | 0.010–0.030 |
| | RMSPE | 88.87 | 91.24 | 91.40 | 94.79 | 95.24 | 100.50 | 101.46 | 0.47–1.77 |
| | MAPE | 72.22 | 74.15 | 74.26 | 76.93 | 77.26 | 81.71 | 82.42 | 0.32–1.54 |
| RF-`q` | PA | 0.730 | 0.709 | 0.702 | 0.680 | 0.669 | 0.643 | 0.613 | 0.010–0.027 |
| | RMSPE | 91.22 | 93.40 | 93.60 | 96.98 | 97.37 | 102.61 | 103.44 | 0.44–1.51 |
| | MAPE | 74.30 | 75.88 | 76.10 | 78.73 | 78.96 | 83.38 | 83.97 | 0.30–1.36 |

**Competing methods:** ■ Highest PAs; ■ Smallest PEs; ■ Smallest PAs; ■ Highest PEs.

Table S8: Percentage loss in predictive accuracy (PA) under `variance-inflated` contamination, computed from the 0% scenario ($PA_{0\%}$) to the smallest observed PA across all `variance-inflated` scenarios ($s = 5, 7$; 2%, 5%, 10%).

| **Method** | $\mathbf{PA_{0\%}}$ | **Min PA (Var.-infl.)** | **% Loss** |
|---|---|---|---|
| RF | 0.755 | 0.284 | 62.4 |
| | **Preprocessing-based Approaches** | | |
| RF-`k` | 0.739 | 0.688 | 6.9 |
| RF-`BC` | 0.747 | 0.291 | 61.0 |
| RF-`YJ` | 0.755 | 0.283 | 62.5 |
| RF-`w` | 0.731 | 0.698 | 4.5 |
| | **Algorithm-based approaches** | | |
| RF-`m` | 0.738 | 0.614 | 16.8 |
| RF-`q` | 0.730 | 0.613 | 16.0 |

Table S9: Estimated predictive accuracy (PA), mean squared prediction error (RMSPE) and mean absolute error (MAPE) for the non-contaminated and `central variance-deflated` contaminated scenarios — Preprocessing- and Algorithm-based approaches. *sd* is the standard deviation. **Stage 3**

| **Method** | | **0%** | **2%** $\gamma = 10^3$ | **2%** $\gamma = 10^4$ | **5%** $\gamma = 10^3$ | **5%** $\gamma = 10^4$ | **10%** $\gamma = 10^3$ | **10%** $\gamma = 10^4$ | *sd* range |
|---|---|---|---|---|---|---|---|---|---|
| RF | PA | 0.755 | 0.748 | 0.747 | 0.742 | 0.740 | 0.738 | 0.739 | 0.003–0.007 |
| | RMSPE | 88.61 | 89.93 | 89.95 | 91.98 | 92.05 | 94.78 | 94.77 | 0.28–0.61 |
| | MAPE | 72.13 | 73.11 | 73.13 | 74.75 | 74.81 | 76.97 | 76.97 | 0.22–0.48 |
| **a)** | | **Preprocessing-based Approaches** | | | | | | | |
| RF-`k` | PA | 0.739 | 0.726 | 0.725 | 0.691 | 0.690 | 0.653 | 0.652 | 0.004-0.009 |
| (i) | RMSPE | 95.10 | 97.61 | 97.62 | 100.88 | 100.89 | 102.55 | 102.53 | 0.12–0.24 |
| | MAPE | 77.21 | 79.16 | 79.20 | 81.59 | 81.59 | 82.66 | 82.65 | 0.10–0.17 |
| (ii) | RMSPE | 93.93 | 96.33 | 96.34 | 99.52 | 99.53 | 101.23 | 101.20 | 0.13–0.28 |
| | MAPE | 76.11 | 78.04 | 78.95 | 80.45 | 80.45 | 81.57 | 81.55 | 0.13–0.21 |
| RF-`w` | PA | 0.731 | 0.732 | 0.732 | 0.730 | 0.729 | 0.729 | 0.728 | 0.003–0.008 |
| | RMSPE | 92.98 | 94.33 | 94.29 | 96.28 | 96.28 | 99.35 | 99.37 | 0.20–0.41 |
| | MAPE | 75.37 | 76.58 | 76.54 | 78.13 | 78.14 | 80.64 | 80.66 | 0.17–0.35 |

**Competing methods:** ■ Highest PAs; ■ Smallest PEs; ■ Smallest PAs; ■ Highest PEs.

| **Method** | | **0%** | **2%** $\gamma = 10^3$ | **2%** $\gamma = 10^4$ | **5%** $\gamma = 10^3$ | **5%** $\gamma = 10^4$ | **10%** $\gamma = 10^3$ | **10%** $\gamma = 10^4$ | *sd* range |
|---|---|---|---|---|---|---|---|---|---|
| **b)** | | **Algorithm-based approaches** | | | | | | | |
| RF-`m` | PA | 0.738 | 0.730 | 0.729 | 0.711 | 0.707 | 0.679 | 0.680 | 0.005–0.008 |
| | RMSPE | 88.87 | 91.68 | 91.73 | 95.42 | 95.56 | 99.10 | 99.03 | 0.31–0.57 |
| | MAPE | 72.22 | 74.50 | 74.54 | 77.40 | 77.55 | 80.06 | 80.00 | 0.21–0.38 |
| RF-`q` | PA | 0.730 | 0.719 | 0.717 | 0.697 | 0.694 | 0.661 | 0.661 | 0.006-0.009 |
| | RMSPE | 91.22 | 94.08 | 94.12 | 97.85 | 97.94 | 100.72 | 100.67 | 0.24–0.37 |
| | MAPE | 74.30 | 76.43 | 76.47 | 79.32 | 79.42 | 81.28 | 81.25 | 0.20–0.30 |

**Competing methods:** ■ Highest PAs; ■ Smallest PEs; ■ Smallest PAs; ■ Highest PEs.

Table S10: Percentage loss in predictive accuracy (PA) under `central variance-deflated` contamination (`C.Var.-defl.`), computed from the 0% scenario ($PA_{0\%}$) to the smallest observed PA across all `central variance-deflated` scenarios ($\gamma = 1000, 10000$; 2%, 5%, 10%).

| **Method** | **$PA_{0\%}$** | **Min PA (C.Var.-defl.)** | **% Loss** |
|---|---|---|---|
| RF | 0.755 | 0.738 | 2.3 |
| | **Preprocessing-based Approaches** | | |
| RF-k | 0.739 | 0.652 | 11.8 |
| RF-w | 0.731 | 0.728 | 0.4 |
| | **Algorithm-based approaches** | | |
| RF-m | 0.738 | 0.679 | 8.0 |
| RF-q | 0.730 | 0.661 | 9.5 |

Table S11: Estimated predictive accuracy (PA), mean squared prediction error (RMSPE) and mean absolute error (MAPE) for the non-contaminated and `tail variance-deflated` contaminated scenarios — Preprocessing- and Algorithm-based approaches. *sd* is the standard deviation. **Stage 4**

| Method | | 0% | 2% | | | 5% | | | 10% | | | *sd* |
|---|---|---|---|---|---|---|---|---|---|---|---|---|
| | | | $\gamma = 10^4$ | | | | | | | | | |
| | | | $k=5$ | $k=7$ | $k=9$ | $k=5$ | $k=7$ | $k=9$ | $k=5$ | $k=7$ | $k=9$ | range |
| RF | PA | 0.755 | 0.694 | 0.617 | 0.548 | 0.615 | 0.541 | 0.445 | 0.618 | 0.513 | 0.446 | 0.10–0.047 |
| | RMSPE | 88.61 | 85.44 | 87.51 | 91.79 | 89.09 | 102.50 | 120.92 | 108.27 | 145.24 | 182.08 | 0.53–6.22 |
| | MAPE | 72.13 | 68.72 | 69.81 | 72.08 | 70.74 | 80.66 | 94.11 | 86.68 | 120.23 | 154.66 | 0.41–6.60 |
| **a)** | **Preprocessing-based Approaches** | | | | | | | | | | | |
| RF-`k` | PA | 0.739 | 0.729 | 0.732 | 0.734 | 0.732 | 0.730 | 0.731 | 0.723 | 0.716 | 0.722 | 0.004–0.009 |
| (i) | RMSPE | 95.10 | 94.96 | 94.84 | 94.89 | 94.61 | 94.74 | 94.74 | 94.05 | 94.22 | 94.08 | 0.23–0.47 |
| | MAPE | 77.21 | 77.03 | 76.94 | 76.96 | 76.79 | 76.84 | 76.83 | 76.31 | 76.39 | 76.30 | 0.20–0.41 |
| (ii) | RMSPE | 93.93 | 93.85 | 93.71 | 93.75 | 93.52 | 93.65 | 93.66 | 93.07 | 93.27 | 93.07 | 0.23–0.53 |
| | MAPE | 76.11 | 75.99 | 75.90 | 75.92 | 75.77 | 75.83 | 75.82 | 75.39 | 75.48 | 75.33 | 0.19–0.45 |
| RF-`w` | PA | 0.731 | 0.729 | 0.732 | 0.733 | 0.732 | 0.731 | 0.730 | 0.727 | 0.723 | 0.727 | 0.004–0.008 |
| | RMSPE | 92.98 | 92.40 | 92.30 | 92.29 | 91.05 | 91.06 | 91.19 | 88.93 | 88.09 | 88.97 | 0.22–0.48 |
| | MAPE | 75.37 | 74.98 | 74.91 | 74.89 | 73.78 | 73.79 | 73.88 | 71.69 | 71.84 | 71.81 | 0.16–0.39 |

**Competing methods:** ■ Highest PAs; ■ Smallest PEs; ■ Smallest PAs; ■ Highest PEs.

| Method | | 0% | 2% | | | 5% | | | 10% | | | *sd* |
|---|---|---|---|---|---|---|---|---|---|---|---|---|
| | | | $k=5$ | $k=7$ | $k=9$ | $k=5$ | $k=7$ | $k=9$ | $k=5$ | $k=7$ | $k=9$ | range |
| **b)** | **Algorithm-based approaches** | | | | | | | | | | | |
| RF-`m` | PA | 0.738 | 0.717 | 0.706 | 0.702 | 0.701 | 0.684 | 0.670 | 0.679 | 0.645 | 0.626 | 0.005–0.022 |
| | RMSPE | 88.87 | 88.88 | 88.97 | 88.97 | 88.90 | 88.98 | 89.63 | 88.11 | 89.43 | 90.20 | 0.26–1.38 |
| | MAPE | 72.22 | 72.21 | 72.28 | 72.23 | 72.17 | 72.00 | 72.54 | 71.35 | 72.04 | 72.88 | 0.22–1.08 |
| RF-`q` | PA | 0.730 | 0.707 | 0.699 | 0.697 | 0.699 | 0.689 | 0.673 | 0.682 | 0.655 | 0.643 | 0.005–0.021 |
| | RMSPE | 91.22 | 91.43 | 91.54 | 91.43 | 91.67 | 91.65 | 92.28 | 91.18 | 92.30 | 92.77 | 0.28–1.35 |
| | MAPE | 74.30 | 74.34 | 74.39 | 74.22 | 74.38 | 74.22 | 74.75 | 73.92 | 74.62 | 75.14 | 0.24–1.08 |

**Competing methods:** ■ Highest PAs; ■ Smallest PEs; ■ Smallest PAs; ■ Highest PEs.

Table S12: Percentage loss in predictive accuracy (PA) under `tail variance-deflated` contamination ($\gamma = 10^4$), computed from the 0% scenario ($PA_{0\%}$) to the smallest observed PA across all settings ($k = 5, 7, 9$; 2%, 5%, 10%).

| Method | $PA_{0\%}$ | Min PA | % Loss |
|---|---|---|---|
| RF | 0.755 | 0.445 | 41.1 |
| RF-`k` | 0.739 | 0.716 | 3.1 |
| RF-`w` | 0.731 | 0.723 | 1.1 |
| RF-`m` | 0.738 | 0.626 | 15.2 |
| RF-`q` | 0.730 | 0.643 | 11.9 |

Table S13: Estimated predictive accuracy (PA), mean squared prediction error (RMSPE) and mean absolute error (MAPE) for the non-contaminated and `shift` contaminated scenarios — Hybrid Approaches. *sd* is the standard deviation. **Stage 1**

| **Method** | | **Contamination Level** | | | | | | | | | | *sd* |
|---|---|---|---|---|---|---|---|---|---|---|---|---|
| | | **0%** | **2%** | | | **5%** | | | **10%** | | | |
| | | | $k=5$ | $k=7$ | $k=9$ | $k=5$ | $k=7$ | $k=9$ | $k=5$ | $k=7$ | $k=9$ | range |
| RF | PA | 0.755 | 0.683 | 0.610 | 0.546 | 0.636 | 0.532 | 0.456 | 0.609 | 0.499 | 0.430 | 0.012–0.044 |
| | RMSPE | 88.61 | 85.62 | 87.83 | 91.71 | 89.88 | 103.74 | 121.68 | 109.11 | 144.85 | 183.71 | 0.78–5.50 |
| | MAPE | 72.13 | 68.90 | 70.08 | 72.48 | 71.24 | 81.59 | 94.14 | 87.20 | 119.36 | 155.69 | 0.74–5.31 |
| **Hybrid Approaches** | | | | | | | | | | | | |
| RF-`k-m` (ii) | PA | 0.728 | 0.721 | 0.719 | 0.720 | 0.717 | 0.714 | 0.717 | 0.713 | 0.710 | 0.709 | 0.005–0.007 |
| | RMSPE | 88.53 | 90.43 | 90.55 | 90.45 | 90.75 | 90.80 | 90.62 | 90.85 | 91.12 | 91.03 | 0.29–0.43 |
| | MAPE | 71.64 | 73.45 | 73.51 | 73.42 | 73.70 | 73.80 | 73.65 | 73.78 | 73.92 | 73.90 | 0.23–0.41 |
| RF-`k-q` (ii) | PA | 0.718 | 0.711 | 0.709 | 0.711 | 0.709 | 0.705 | 0.708 | 0.704 | 0.701 | 0.702 | 0.003–0.009 |
| | RMSPE | 90.52 | 90.28 | 90.45 | 90.11 | 90.00 | 90.18 | 89.90 | 89.17 | 89.29 | 89.20 | 0.26–0.51 |
| | MAPE | 73.38 | 73.17 | 73.25 | 72.96 | 72.89 | 73.15 | 72.83 | 72.20 | 72.17 | 72.16 | 0.22–0.56 |
| RF-`w-m` | PA | 0.714 | 0.719 | 0.719 | 0.722 | 0.719 | 0.721 | 0.721 | 0.714 | 0.714 | 0.714 | 0.004–0.009 |
| | RMSPE | 90.21 | 89.81 | 89.79 | 89.57 | 89.16 | 89.04 | 89.05 | 87.85 | 87.84 | 87.82 | 0.22–0.61 |
| | MAPE | 73.06 | 72.98 | 72.87 | 72.71 | 72.46 | 72.36 | 72.35 | 71.31 | 71.24 | 71.32 | 0.16–0.49 |
| RF-`w-q` | PA | 0.709 | 0.712 | 0.708 | 0.713 | 0.713 | 0.710 | 0.712 | 0.709 | 0.708 | 0.705 | 0.005–0.009 |
| | RMSPE | 92.31 | 91.72 | 91.86 | 91.56 | 91.11 | 91.36 | 91.14 | 90.08 | 90.12 | 90.23 | 0.24–0.53 |
| | MAPE | 74.97 | 74.52 | 74.58 | 74.32 | 73.96 | 74.27 | 74.10 | 73.16 | 73.10 | 73.29 | 0.17–0.39 |

**Competing methods:** ■ Highest PAs; ■ Smallest PEs; ■ Smallest PAs; ■ Highest PEs.

Table S14: Estimated predictive accuracy (PA), mean squared prediction error (RMSPE) and mean absolute error (MAPE) for the non-contaminated and `variance-inflated` contaminated scenarios — Hybrid Approaches. *sd* is the standard deviation. **Stage 2**

| **Method** | | **Contamination Level** | | | | | | | *sd* |
|---|---|---|---|---|---|---|---|---|---|
| | | **0%** | **2%** | | **5%** | | **10%** | | |
| | | | $s = 5$ | $s = 7$ | $s = 5$ | $s = 7$ | $s = 5$ | $s = 7$ | range |
| RF | PA | 0.755 | 0.661 | 0.571 | 0.536 | 0.409 | 0.429 | 0.284 | 0.04–0.08 |
| | MAPE | 72.13 | 74.34 | 75.95 | 78.17 | 81.95 | 84.00 | 92.22 | 1.91–11.21 |
| | RMSPE | 88.61 | 91.96 | 95.29 | 97.77 | 105.49 | 105.33 | 120.88 | 1.33–6.28 |
| **Hybrid Approaches** | | | | | | | | | |
| RF-`k-m` (ii) | PA | 0.728 | 0.718 | 0.716 | 0.703 | 0.699 | 0.683 | 0.682 | 0.008–0.14 |
| | RMSPE | 88.53 | 89.05 | 89-12 | 90.25 | 90.35 | 91.93 | 92.02 | 0.36–0.97 |
| | MAPE | 71.64 | 72.12 | 72.15 | 73.02 | 73.09 | 74.43 | 74.42 | 0.32–0.82 |
| RF-`k-q` (ii) | PA | 0.718 | 0.707 | 0.706 | 0.696 | 0.695 | 0.674 | 0.677 | 0.009–0.015 |
| | RMSPE | 90.52 | 91.88 | 91.90 | 94.50 | 94.50 | 99.64 | 99.35 | 0.35–1.36 |
| | MAPE | 73.38 | 74.53 | 74.54 | 76.68 | 76.72 | 80.94 | 80.72 | 0.30–1.10 |
| RF-`w-m` | PA | 0.714 | 0.717 | 0.716 | 0.703 | 0.705 | 0.691 | 0.689 | 0.005–0.11 |
| | RMSPE | 90.21 | 91.65 | 91.71 | 94.59 | 94.42 | 99.27 | 99.32 | 0.35–1.16 |
| | MAPE | 73.06 | 74.40 | 74.45 | 76.77 | 76.66 | 80.62 | 80.69 | 0.26–0.94 |
| RF-`w-q` | PA | 0.709 | 0.707 | 0.708 | 0.696 | 0.695 | 0.684 | 0.682 | 0.006–0.011 |
| | RMSPE | 92.31 | 93.61 | 93.64 | 96.36 | 96.34 | 101.21 | 101.16 | 0.27–1.13 |
| | MAPE | 74.97 | 76.02 | 76.05 | 78.23 | 78.22 | 82.25 | 82.20 | 0.20–0.90 |

**Competing methods:** ■ Highest PAs; ■ Smallest PEs; ■ Smallest PAs; ■ Highest PEs.

Table S15: Estimated predictive accuracy (PA), mean squared prediction error (RMSPE) and mean absolute error (MAPE) for the non-contaminated and `central variance-deflated` contaminated scenarios — Hybrid Approaches. *sd* is the standard deviation.

**Stage 3**

| **Method** | | **Contamination Level** 0% | 2% $\gamma = 10^3$ | 2% $\gamma = 10^4$ | 5% $\gamma = 10^3$ | 5% $\gamma = 10^4$ | 10% $\gamma = 10^3$ | 10% $\gamma = 10^4$ | *sd* range |
|---|---|---|---|---|---|---|---|---|---|
| RF | PA | 0.755 | 0.748 | 0.747 | 0.742 | 0.740 | 0.738 | 0.739 | 0.003–0.007 |
| | RMSPE | 88.61 | 89.93 | 89.95 | 91.98 | 92.05 | 94.78 | 94.77 | 0.28–0.61 |
| | MAPE | 72.13 | 73.11 | 73.13 | 74.75 | 74.81 | 76.97 | 76.97 | 0.22–0.48 |
| **Hybrid Approaches** | | | | | | | | | |
| RF-`k-m` (ii) | PA | 0.728 | 0.719 | 0.719 | 0.714 | 0.715 | 0.710 | 0.710 | 0.004–0.008 |
| | RMSPE | 88.53 | 88.74 | 88.73 | 89.39 | 89.33 | 89.73 | 89.72 | 0.17–0.51 |
| | MAPE | 71.64 | 71.87 | 71.84 | 72.30 | 72.26 | 72.67 | 72.66 | 0.20–0.44 |
| RF-`k-q` (ii) | PA | 0.718 | 0.705 | 0.706 | 0.680 | 0.680 | 0.644 | 0.645 | 0.003–0.007 |
| | RMSPE | 90.52 | 92.97 | 92.96 | 96.43 | 96.50 | 99.18 | 99.16 | 0.18–0.47 |
| | MAPE | 73.38 | 75.35 | 75.35 | 78.00 | 78.07 | 79.99 | 80.00 | 0.16–0.38 |
| RF-`w-m` | PA | 0.714 | 0.713 | 0.713 | 0.696 | 0.694 | 0.667 | 0.668 | 0.005–0.007 |
| | RMSPE | 90.21 | 92.76 | 92.74 | 96.30 | 96.28 | 99.69 | 99.67 | 0.25–0.42 |
| | MAPE | 73.06 | 75.31 | 75.29 | 78.04 | 78.03 | 80.53 | 80.49 | 0.21–0.34 |
| RF-`w-q` | PA | 0.709 | 0.705 | 0.702 | 0.682 | 0.683 | 0.651 | 0.652 | 0.004–0.010 |
| | RMSPE | 92.31 | 94.67 | 94.79 | 98.30 | 98.22 | 100.91 | 100.90 | 0.20–0.42 |
| | MAPE | 74.97 | 76.78 | 76.91 | 79.65 | 79.50 | 81.45 | 81.48 | 0.16–0.36 |

**Competing methods:** ■ Highest PAs; ■ Smallest PEs; ■ Smallest PAs; ■ Highest PEs.

Table S16: Estimated predictive accuracy (PA), mean squared prediction error (RMSPE) and mean absolute error (MAPE) for the non-contaminated and `tail variance-deflated` contaminated scenarios — Preprocessing- and Algorithm-based approaches. *sd* is the standard deviation. **Stage 4**

| Method | | 0% | 2% | | | 5% | | | 10% | | | *sd* |
|---|---|---|---|---|---|---|---|---|---|---|---|---|
| | | | Contamination Level, $\gamma = 10^4$ | | | | | | | | | |
| | | | $k=5$ | $k=7$ | $k=9$ | $k=5$ | $k=7$ | $k=9$ | $k=5$ | $k=7$ | $k=9$ | range |
| RF | PA | 0.755 | 0.694 | 0.617 | 0.548 | 0.615 | 0.541 | 0.445 | 0.618 | 0.513 | 0.446 | 0.10–0.047 |
| | RMSPE | 88.61 | 85.44 | 87.51 | 91.79 | 89.09 | 102.50 | 120.92 | 108.27 | 145.24 | 182.08 | 0.53–6.22 |
| | MAPE | 72.13 | 68.72 | 69.81 | 72.08 | 70.74 | 80.66 | 94.11 | 86.68 | 120.23 | 154.66 | 0.41–6.60 |
| **Hybrid Approaches** | | | | | | | | | | | | |
| RF-`k-m` (ii) | PA | 0.728 | 0.716 | 0.719 | 0.721 | 0.721 | 0.719 | 0.720 | 0.714 | 0.707 | 0.714 | 0.005–0.007 |
| | RMSPE | 88.53 | 88.45 | 88.31 | 88.34 | 87.78 | 87.90 | 87.94 | 87.03 | 87.30 | 87.07 | 0.26–0.48 |
| | MAPE | 71.64 | 71.65 | 71.56 | 71.54 | 71.14 | 71.23 | 71.94 | 70.45 | 70.62 | 70.44 | 0.23–0.43 |
| RF-`k-q` (ii) | PA | 0.718 | 0.708 | 0.710 | 0.711 | 0.715 | 0.711 | 0.711 | 0.705 | 0.702 | 0.705 | 0.004–0.009 |
| | RMSPE | 90.52 | 90.47 | 90.30 | 90.35 | 89.97 | 89.86 | 89.95 | 89.25 | 89.38 | 89.30 | 0.28–0.58 |
| | MAPE | 73.38 | 73.28 | 73.13 | 73.17 | 72.78 | 72.92 | 72.87 | 72.22 | 72.32 | 72.25 | 0.18–0.52 |
| RF-`w-m` | PA | 0.714 | 0.718 | 0.719 | 0.721 | 0.721 | 0.720 | 0.719 | 0.719 | 0.714 | 0.718 | 0.004–0.009 |
| | RMSPE | 90.21 | 89.88 | 89.91 | 89.83 | 89.14 | 89.26 | 89.24 | 87.69 | 87.82 | 87.80 | 0.22–0.51 |
| | MAPE | 73.06 | 73.01 | 73.05 | 72.98 | 72.41 | 72.52 | 72.46 | 71.13 | 71.21 | 71.23 | 0.24–0.49 |
| RF-`w-q` | PA | 0.709 | 0.709 | 0.709 | 0.710 | 0.715 | 0.712 | 0.710 | 0.713 | 0.707 | 0.710 | 0.004–0.009 |
| | RMSPE | 92.31 | 91.83 | 91.87 | 91.77 | 91.21 | 91.35 | 91.35 | 89.98 | 90.18 | 90.18 | 0.17–0.54 |
| | MAPE | 74.97 | 74.60 | 74.68 | 74.56 | 74.05 | 74.23 | 74.14 | 73.05 | 73.15 | 73.19 | 0.21–0.44 |

**Competing methods:** ■ Highest PAs; ■ Smallest PEs; ■ Smallest PAs; ■ Highest PEs.

Table S17: Baseline efficiency and robustness of hybrid approaches in terms of predictive accuracy (PA). The column "% Loss vs RF (0%)" reports the relative decrease in PA under the uncontaminated scenario with respect to standard Random Forests (RF) ($\text{PA}_{0\%}$ = 0.755). For each contamination scenario, the loss is computed from the method's 0% PA to the smallest observed PA across all settings within that scenario.

| Method | $\text{PA}_{0\%}$ | % Loss vs RF (0%) | `Shift` | | `Var.-infl.` | | `C.Var.-defl.` | | `T.Var.-defl.` | |
|---|---|---|---|---|---|---|---|---|---|---|
| | | | Min PA | % Loss | Min PA | % Loss | Min PA | % Loss | Min PA | % Loss |
| RF | 0.755 | 0.0 | 0.430 | 43.0 | 0.284 | 62.4 | 0.738 | 2.3 | 0.445 | 41.1 |
| **Hybrid Approaches** | | | | | | | | | | |
| RF-`k-m` (ii) | 0.728 | 3.6 | 0.709 | 2.6 | 0.684 | 6.0 | 0.710 | 2.5 | 0.707 | 2.9 |
| RF-`k-q` (ii) | 0.718 | 4.9 | 0.701 | 2.3 | 0.674 | 6.1 | 0.644 | 10.2 | 0.702 | 2.2 |
| RF-`w-m` | 0.714 | 5.4 | 0.714 | 0.0 | 0.689 | 3.5 | 0.667 | 6.6 | 0.714 | 0.0 |
| RF-`w-q` | 0.709 | 6.1 | 0.705 | 0.6 | 0.682 | 3.8 | 0.651 | 8.2 | 0.709 | 0.0 |

Table S18: Breakdown-point assessment (stress-test) of selected best performing methods across all contamination scenarios. Standard radom forests (RF) performance (PA: predictive accuracy; RMSPE: mean squared prediction error; and MAPE: mean absolute error) is included for reference. *sd* is the standard deviation.

| | | Contamination level | | | | | | | | | | | | | | | | | | | | | | | | | | | | | | | |
|---|---|---|---|---|---|---|---|---|---|---|---|---|---|---|---|---|---|---|---|---|---|---|---|---|---|---|---|---|---|---|---|---|---|
| | | 0% | 15% | | | | | | | | | | 20% | | | | | | | | | | 25% | | | | | | | | | | |
| Method | Metric | NoCont | Shift | | | Var.-infl. | | C.Var.-defl. | | T.Var.-defl. | | | Shift | | | Var.-infl. | | C.Var.-defl. | | T.Var.-defl. | | | Shift | | | Var.-infl. | | C.Var.-defl. | | T.Var.-defl. | | | *sd* |
| | | – | $k=5$ | $k=7$ | $k=9$ | $s=5$ | $s=7$ | $\gamma=10^3$ | $\gamma=10^4$ | $k=5$ | $k=7$ | $k=9$ | $k=5$ | $k=7$ | $k=9$ | $s=5$ | $s=7$ | $\gamma=10^3$ | $\gamma=10^4$ | $k=5$ | $k=7$ | $k=9$ | $k=5$ | $k=7$ | $k=9$ | $s=5$ | $s=7$ | $\gamma=10^3$ | $\gamma=10^4$ | $k=5$ | $k=7$ | $k=9$ | range |
| RF | PA | 0.755 | 0.599 | 0.522 | 0.473 | 0.381 | 0.280 | 0.732 | 0.735 | 0.615 | 0.535 | 0.482 | 0.612 | 0.539 | 0.492 | 0.338 | 0.232 | 0.732 | 0.732 | 0.630 | 0.550 | 0.505 | 0.630 | 0.574 | 0.567 | 0.342 | 0.210 | 0.725 | 0.726 | 0.640 | 0.588 | 0.575 | 0.005–0.093 |
| | RMSPE | 88.61 | 136.40 | 190.82 | 249.88 | 107.78 | 123.65 | 98.58 | 98.04 | 134.09 | 188.55 | 246.88 | 164.44 | 236.98 | 316.25 | 112.71 | 129.86 | 101.92 | 102.11 | 161.52 | 234.17 | 310.82 | 195.25 | 286.67 | 374.95 | 116.57 | 130.61 | 106.33 | 106.26 | 192.21 | 283.03 | 372.13 | 0.47–9.67 |
| | MAPE | 72.13 | 113.85 | 168.53 | 228.33 | 85.71 | 93.26 | 80.01 | 79.62 | 111.83 | 166.85 | 225.78 | 143.65 | 218.63 | 299.52 | 89.93 | 99.98 | 82.72 | 82.96 | 141.18 | 216.14 | 294.61 | 177.38 | 272.32 | 362.21 | 93.02 | 101.64 | 86.27 | 86.21 | 174.34 | 269.03 | 359.77 | 0.36–7.16 |
| **Preprocessing-level** | | | | | | | | | | | | | | | | | | | | | | | | | | | | | | | | | |
| RF-k (ii) | PA | 0.739 | 0.716 | 0.716 | 0.717 | 0.680 | 0.681 | 0.645 | 0.637 | 0.717 | 0.716 | 0.717 | 0.708 | 0.705 | 0.712 | 0.648 | 0.640 | 0.639 | 0.639 | 0.709 | 0.705 | 0.712 | 0.706 | 0.702 | 0.705 | 0.616 | 0.609 | 0.638 | 0.630 | 0.706 | 0.702 | 0.705 | 0.004–0.043 |
| | RMSPE | 93.93 | 92.52 | 92.42 | 92.19 | 107.83 | 107.97 | 101.52 | 101.55 | 92.50 | 92.41 | 92.19 | 90.99 | 91.41 | 91.23 | 113.68 | 114.05 | 101.64 | 101.63 | 90.96 | 91.41 | 91.23 | 89.63 | 89.98 | 89.50 | 119.90 | 120.67 | 101.88 | 101.95 | 89.65 | 89.98 | 89.50 | 0.13–1.57 |
| | MAPE | 76.11 | 74.84 | 74.76 | 74.62 | 87.39 | 87.57 | 81.81 | 81.85 | 74.83 | 74.86 | 74.62 | 73.56 | 73.87 | 73.74 | 92.13 | 92.55 | 81.92 | 81.89 | 73.55 | 73.87 | 73.74 | 72.31 | 72.65 | 72.21 | 97.44 | 98.07 | 82.13 | 82.16 | 72.32 | 72.65 | 72.21 | 0.13–1.30 |
| RF-w | PA | 0.731 | 0.718 | 0.721 | 0.719 | 0.685 | 0.692 | 0.716 | 0.717 | 0.720 | 0.720 | 0.719 | 0.708 | 0.706 | 0.712 | 0.661 | 0.653 | 0.717 | 0.716 | 0.710 | 0.706 | 0.713 | 0.691 | 0.689 | 0.694 | 0.632 | 0.620 | 0.711 | 0.707 | 0.602 | 0.690 | 0.693 | 0.004–0.039 |
| | RMSPE | 92.98 | 87.95 | 87.69 | 87.76 | 102.84 | 102.67 | 102.98 | 102.77 | 87.81 | 87.77 | 87.83 | 90.16 | 90.06 | 90.13 | 106.65 | 106.97 | 106.09 | 106.24 | 90.04 | 90.05 | 90.05 | 116.66 | 114.31 | 110.84 | 111.68 | 111.94 | 109.18 | 109.16 | 117.56 | 113.95 | 110.77 | 0.23–1.87 |
| | MAPE | 75.37 | 70.40 | 70.25 | 70.35 | 83.35 | 83.31 | 85.55 | 83.38 | 70.30 | 70.34 | 70.42 | 71.66 | 71.53 | 71.56 | 85.02 | 85.41 | 86.07 | 86.21 | 71.54 | 71.55 | 71.54 | 95.74 | 93.47 | 90.03 | 90.66 | 90.87 | 88.54 | 88.50 | 96.40 | 92.83 | 89.69 | 0.21–1.52 |
| **Hybrid approaches** | | | | | | | | | | | | | | | | | | | | | | | | | | | | | | | | | |
| RF-k-m (ii) | PA | 0.728 | 0.708 | 0.709 | 0.710 | 0.673 | 0.676 | 0.633 | 0.629 | 0.709 | 0.709 | 0.710 | 0.703 | 0.700 | 0.706 | 0.643 | 0.642 | 0.628 | 0.628 | 0.704 | 0.700 | 0.706 | 0.700 | 0.696 | 0.699 | 0.617 | 0.612 | 0.629 | 0.624 | 0.699 | 0.696 | 0.699 | 0.003–0.33 |
| | RMSPE | 88.53 | 86.17 | 86.06 | 85.87 | 102.64 | 102.70 | 98.78 | 98.89 | 86.13 | 86.06 | 85.87 | 84.41 | 84.83 | 84.59 | 109.43 | 109.65 | 98.99 | 99.02 | 84.38 | 84.83 | 84.59 | 82.86 | 82.01 | 82.63 | 116.32 | 117.22 | 99.17 | 99.22 | 82.90 | 83.00 | 82.64 | 0.24–2.27 |
| | MAPE | 71.64 | 69.65 | 69.57 | 69.44 | 83.19 | 83.37 | 79.58 | 79.69 | 69.63 | 69.57 | 69.44 | 68.19 | 68.42 | 68.25 | 88.69 | 89.00 | 79.74 | 79.73 | 68.22 | 68.42 | 68.25 | 66.67 | 66.87 | 66.60 | 94.60 | 95.32 | 79.82 | 79.90 | 66.72 | 66.87 | 66.60 | 0.21–1.75 |
| RF-w-m | PA | 0.714 | 0.709 | 0.713 | 0.714 | 0.674 | 0.682 | 0.646 | 0.647 | 0.712 | 0.713 | 0.712 | 0.702 | 0.699 | 0.706 | 0.652 | 0.649 | 0.646 | 0.644 | 0.705 | 0.700 | 0.707 | 0.687 | 0.685 | 0.688 | 0.629 | 0.622 | 0.649 | 0.645 | 0.686 | 0.685 | 0.687 | 0.004–0.031 |
| | RMSPE | 90.21 | 86.60 | 86.23 | 86.23 | 105.11 | 104.91 | 100.80 | 100.77 | 86.35 | 86.28 | 86.38 | 84.18 | 84.68 | 84.34 | 110.91 | 111.40 | 101.03 | 101.10 | 83.99 | 84.66 | 84.33 | 82.64 | 82.63 | 82.45 | 117.67 | 118.51 | 101.29 | 101.34 | 82.64 | 82.71 | 82.48 | 0.13–1.60 |
| | MAPE | 73.06 | 70.18 | 69.90 | 69.96 | 85.23 | 85.21 | 81.16 | 81.16 | 69.95 | 69.98 | 70.10 | 68.19 | 68.57 | 68.33 | 90.07 | 90.66 | 81.40 | 81.46 | 67.99 | 68.55 | 68.27 | 66.55 | 66.45 | 66.58 | 95.89 | 96.57 | 81.61 | 81.61 | 66.65 | 66.63 | 66.55 | 0.15–1.27 |

**Competing methods:** ■ Highest PAs; ■ Smallest PEs; ■ Smallest PAs; ■ Highest PEs.

Table S19: Degradation in predictive accuracy (PA) of standard Random Forests (RF) under increasing contamination. Losses are computed relative to the uncontaminated baseline ($PA_{0\%} = 0.755$). The column "$\leq 10\%$" summarizes the worst predictive accuracy observed in the main simulation study (contamination levels 2–10%), whereas the column "15–25%" summarizes the worst accuracy observed in the breakdown-point (BP) analysis.

| **Scenario** | **Min PA** ($\leq 10\%$) | **% Loss** | **Min PA (15–25%)** | **% Loss** |
|---|---|---|---|---|
| `Shift` | 0.430 | 43.0 | 0.473 | 37.4 |
| `Variance-inflated` | 0.284 | 62.4 | 0.210 | 72.2 |
| `C. Variance-deflated` | 0.734 | 2.8 | 0.725 | 4.0 |
| `T. Variance-deflated` | 0.445 | 41.1 | 0.482 | 36.2 |

Table S20: Comparative analysis of standard random forests (RF) against two competing robust RF methods for milk traits $T_1$ (used in the simulations), $T_2$ and $T_3$.

| **Method** \ **Trait** | **T1** | | | **T2** | | | **T3** | | |
|---|---|---|---|---|---|---|---|---|---|
| | **PA** | **RMSPE** | **MAPE** | **PA** | **RMSPE** | **MAPE** | **PA** | **RMSPE** | **MAPE** |
| RF | 0.755 | 88.61 | 72.13 | 0.795 | 7.66 | 6.54 | 0.725 | 0.030 | 0.027 |
| RF-`k` (ii) | 0.739 | 93.93 | 76.11 | 0.787 | 8.27 | 7.06 | 0.724 | 0.030 | 0.027 |
| RF-`w` | 0.731 | 92.98 | 75.37 | 0.791 | 7.88 | 6.71 | 0.716 | 0.031 | 0.027 |

Table S21: Quality of false positives (FP) among the predicted top 10% genotypes. FP are genotypes selected in the predicted top 10% that do not belong to the true top 5%. Higher values of mean and median true percentile indicate that these genotypes remain close to the top of the true ranking.

| Trait | Method | FP | Mean TPerc (%) | Med TPerc (%) | P(5–10) | P(10–20) | P(20–50) | P(B50) |
|---|---|---|---|---|---|---|---|---|
| | RF | 70 | 82.1 | 86.8 | 0.314 | 0.357 | 0.286 | 0.043 |
| $T_1$ | RF-`k` (ii) | 72 | 80.0 | 85.5 | 0.278 | 0.361 | 0.306 | 0.056 |
| | RF-`w` | 76 | 80.9 | 86.7 | 0.303 | 0.342 | 0.316 | 0.039 |
| | RF | 68 | 83.1 | 86.1 | 0.309 | 0.412 | 0.265 | 0.015 |
| $T_2$ | RF-`k` (ii) | 66 | 83.6 | 86.0 | 0.288 | 0.424 | 0.288 | 0.000 |
| | RF-`w` | 65 | 82.4 | 84.1 | 0.308 | 0.369 | 0.308 | 0.015 |
| | RF | 72 | 78.0 | 80.1 | 0.292 | 0.208 | 0.458 | 0.042 |
| $T_3$ | RF-`k` (ii) | 72 | 77.9 | 82.1 | 0.250 | 0.292 | 0.389 | 0.069 |
| | RF-`w` | 73 | 78.3 | 82.8 | 0.260 | 0.301 | 0.370 | 0.068 |

Abbreviations: FP = false positives; TPerc = true percentile - indicates where a genotype stands in the true ranking; P(5–10), P(10–20), and P(20–50) denote the proportion of FP whose true ranking falls within the top 5–10%, 10–20%, and 20–50% of the true distribution, respectively; P(B50) denotes the proportion of FP in the true bottom 50%.

Table S22: Prediction performance of the standard RF, RF-k (ii), and RF-w models across the real maize, soybean, wheat and mice traits. Values represent the mean across the ten independent splits, with standard deviations shown in parentheses. Blue values refer to using transformed $\omega y$ values instead of the suggested shrinkage to the median $m + \omega(y - m)$, with $m = median(y)$. (EH: *ear height*; CIR: *carbon isotope ratio*; CW: *canopy wilting*; GY: *grain yield*; BMI: *body mass index*; BL: *body length*; FNBW: *final normalized body weight*).

| | | RF | | | RF-k (ii) | | | RF-w | | |
|---|---|---|---|---|---|---|---|---|---|---|
| Dataset | Trait | PAb | RMSPE | MAPE | PAb | RMSPE | MAPE | PAb | RMSPE | MAPE |
| Maize | EH | 0.570 (0.087) | 17.40 (2.06) | 13.33 (1.73) | 0.535 (0.075) | 18.38 (1.99) | 14.04 (1.68) | 0.539 (0.079) | 18.10 (1.99) | 13.70 (1.68) |
| | EH | - | - | - | - | - | - | 0.467 (0.068) | 19.21 (1.88) | 14.14 (1.49) |
| Soybean | CIR | 0.300 (0.087) | 0.252 (0.027) | 0.203 (0.021) | 0.324 (0.074) | 0.250 (0.025) | 0.203 (0.019) | 0.292 (0.081) | 0.252 (0.027) | 0.204 (0.021) |
| | CIR | - | - | - | - | - | - | -0.042 (0.107) | 1.78 (0.28) | 1.47 (0.19) |
| | CW | 0.415 (0.079) | 5.82 (0.83) | 4.41 (0.53) | 0.375 (0.065) | 6.04 (1.11) | 4.17 (0.65) | 0.376 (0.065) | 5.91 (1.05) | 4.23 (0.63) |
| | CW | - | - | - | - | - | - | 0.312 (0.070) | 6.12 (1.12) | 4.25 (0.68) |
| Wheat | GY | 0.481 (0.056) | 0.540 (0.024) | 0.421 (0.022) | 0.456 (0.053) | 0.556 (0.028) | 0.437 (0.024) | 0.454 (0.051) | 0.552 (0.028) | 0.432 (0.023) |
| | GY | - | - | - | - | - | - | 0.454 (0.049) | 0.552 (0.028) | 0.432 (0.023) |
| Mice | BMI | 0.453 (0.028) | 0.054 (0.002) | 0.042 (0.002) | 0.452 (0.022) | 0.055 (0.003) | 0.043 (0.002) | 0.455 (0.024) | 0.054 (0.002) | 0.043 (0.002) |
| | BMI | - | - | - | - | - | - | 0.405 (0.033) | 0.058 (0.002) | 0.047 (0.002) |
| | BL | 0.388 (0.025) | 0.509 (0.021) | 0.409 (0.021) | 0.374 ((0.029) | 0.518 ((0.024) | 0.414 ((0.024) | 0.382 (0.028) | 0.514 (0.023) | 0.413 (0.023) |
| | BL | - | - | - | - | - | - | 0.225 (0.038) | 0.650 (0.013) | 0.528 (0.013) |
| | FNBW | 0.729 (0.014) | 3.03 (0.11) | 2.38 (0.11) | 0.730 (0.015) | 3.22 (0.10) | 2.53 (0.09) | 0.727 (0.015) | 3.09 (0.11) | 2.43 (0.10) |
| | FNBW | - | - | - | - | - | - | 0.696 (0.016) | 3.29 (0.11) | 2.57 (0.11) |

Table S23: Ranking-preservation pre-check for the weighted-response random forest (RF-w) across real datasets and traits. Values represent the mean across the ten independent splits, with standard deviations shown in parentheses. Blue values refer to using transformed $\omega y$ values instead of the suggested shrinkage to the median $m + \omega(y - m)$, with $m = median(y)$. (EH: *ear height*; CIR: *carbon isotope ratio*; CW: *canopy wilting*; GY: *grain yield*; BMI: *body mass index*; BL: *body length*; FNBW: *final normalized body weight*).

| Dataset | Trait | Pearson | Spearman | Inversion rate | Min weight | Median weight |
|---|---|---|---|---|---|---|
| Maize | EH | 0.956 (0.003) | 0.999 (0.001) | 0.007 (0.003) | 0.335 (0.008) | 1.000 (0.000) |
| | EH | 0.751 (0.025) | 0.890 (0.016) | 0.075 (0.007) | 0.335 (0.008) | 1.000 (0.000) |
| Soybean | CIR | 0.977 (0.002) | 0.997 (0.001) | 0.013 (0.003) | 0.477 (0.026) | 1.000 (0.000) |
| | CIR | -0.083 (0.063) | 0.433 (0.024) | 0.210 (0.010) | 0.477 (0.026) | 1.000 (0.000) |
| | CW | 0.918 (0.010) | 0.996 (0.001) | 0.014 (0.002) | 0.271 (0.019) | 1.000 (0.000) |
| | CW | 0.622 (0.046) | 0.877 (0.025) | 0.083 (0.011) | 0.271 (0.019) | 1.000 (0.000) |
| Wheat | GY | 0.966 (0.003) | 0.999 (0.000) | 0.007 (0.001) | 0.365 (0.006) | 1.000 (0.000) |
| | GY | 0.966 (0.003) | 0.999 (0.000) | 0.007 (0.002) | 0.365 (0.006) | 1.000 (0.000) |
| Mice | BMI | 0.979 (0.000) | 0.998 (0.000) | 0.008 (0.001) | 0.441 (0.009) | 1.000 (0.000) |
| | BMI | 0.797 (0.013) | 0.852 (0.011) | 0.075 (0.005) | 0.441 (0.009) | 1.000 (0.000) |
| | BL | 0.978 (0.002) | 0.999 (0.000) | 0.007 (0.001) | 0.455 (0.011) | 1.000 (0.000) |
| | BL | 0.607 (0.020) | 0.758 (0.019) | 0.105 (0.008) | 0.455 (0.011) | 1.000 (0.000) |
| | FNBW | 0.989 (0.001) | 0.999 (0.000) | 0.005 (0.000) | 0.452 (0.053) | 1.000 (0.000) |
| | FNBW | 0.866 (0.011) | 0.927 (0.006) | 0.049 (0.002) | 0.452 (0.053) | 1.000 (0.000) |

Inversion rate: proportion of pairs whose ordering is reversed after weighting.

Table S24: Qualitative interpretation of the ranking-preservation pre-check for the weighted-response random forest (RF-w) across datasets and traits. Blue values refer to using transformed $\omega y$ values instead of the suggested shrinkage to the median $m + \omega(y - m)$, with $m = median(y)$. (EH: *ear height*; CIR: *carbon isotope ratio*; CW: *canopy wilting*; GY: *grain yield*; BMI: *body mass index*; BL: *body length*; FNBW: *final normalized body weight*)

| Dataset | Trait | Ranking preservation | Expected RF-w behaviour |
|---|---|---|---|
| Maize | EH | excellent | best |
| | EH | good | good |
| Soybean | CIR | excellent | best |
| | CIR | poor | likely to fail |
| | CW | excellent | best |
| | CW | good | good |
| Wheat | GY | excellent | best |
| | GY | excellent | best |
| Mice | BMI | excellent | best |
| | BMI | good | good |
| | BL | excellent | best |
| | BL | borderline | potentially unstable |
| | FNBW | excellent | best |
| | FNBW | good | good |

### Figures

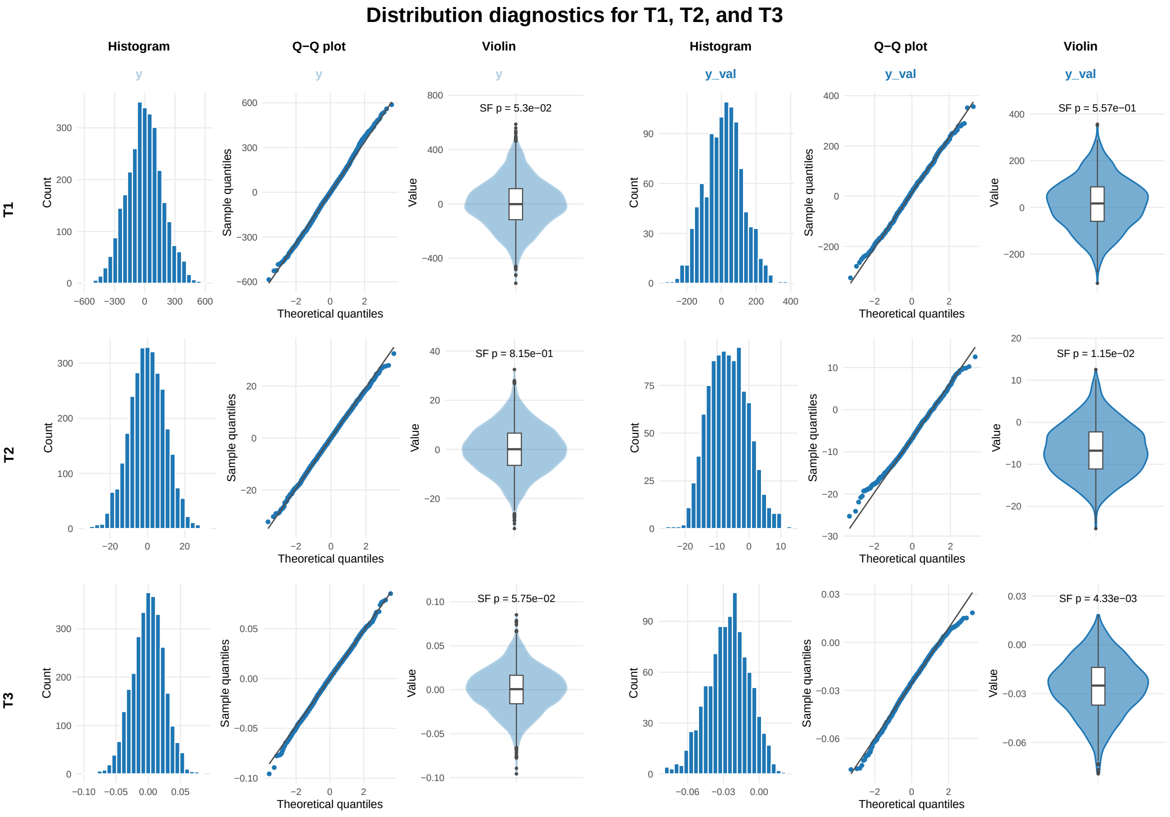


Figure S1: Distribution diagnostic plots for the three animal milk traits in the simulated animal dataset. Shapiro–Francia normality test p-values are shown alongside the violin plots.

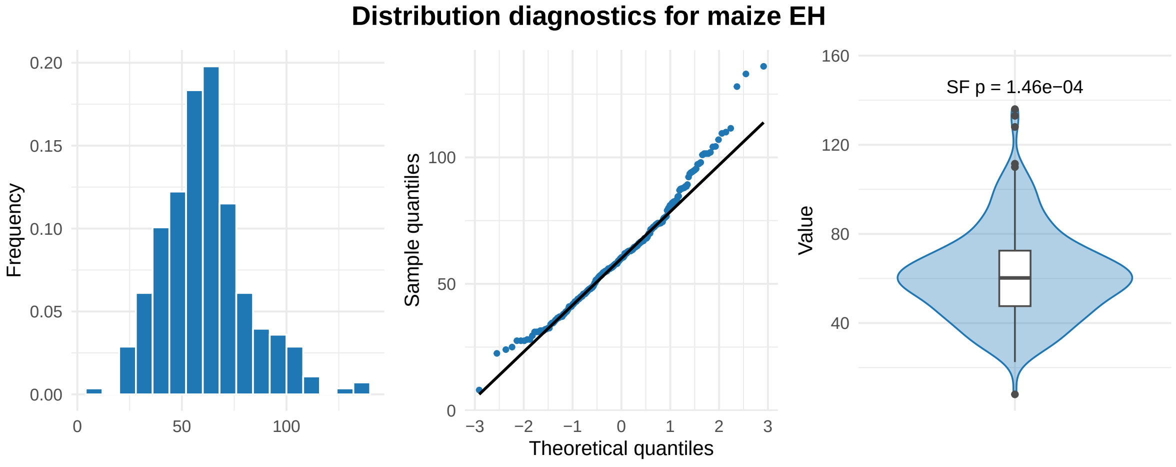


Figure S2: Distribution diagnostic plots for the maize trait (*ear height* in *cm*). Shapiro–Francia normality test p-values are shown alongside the violin plots.

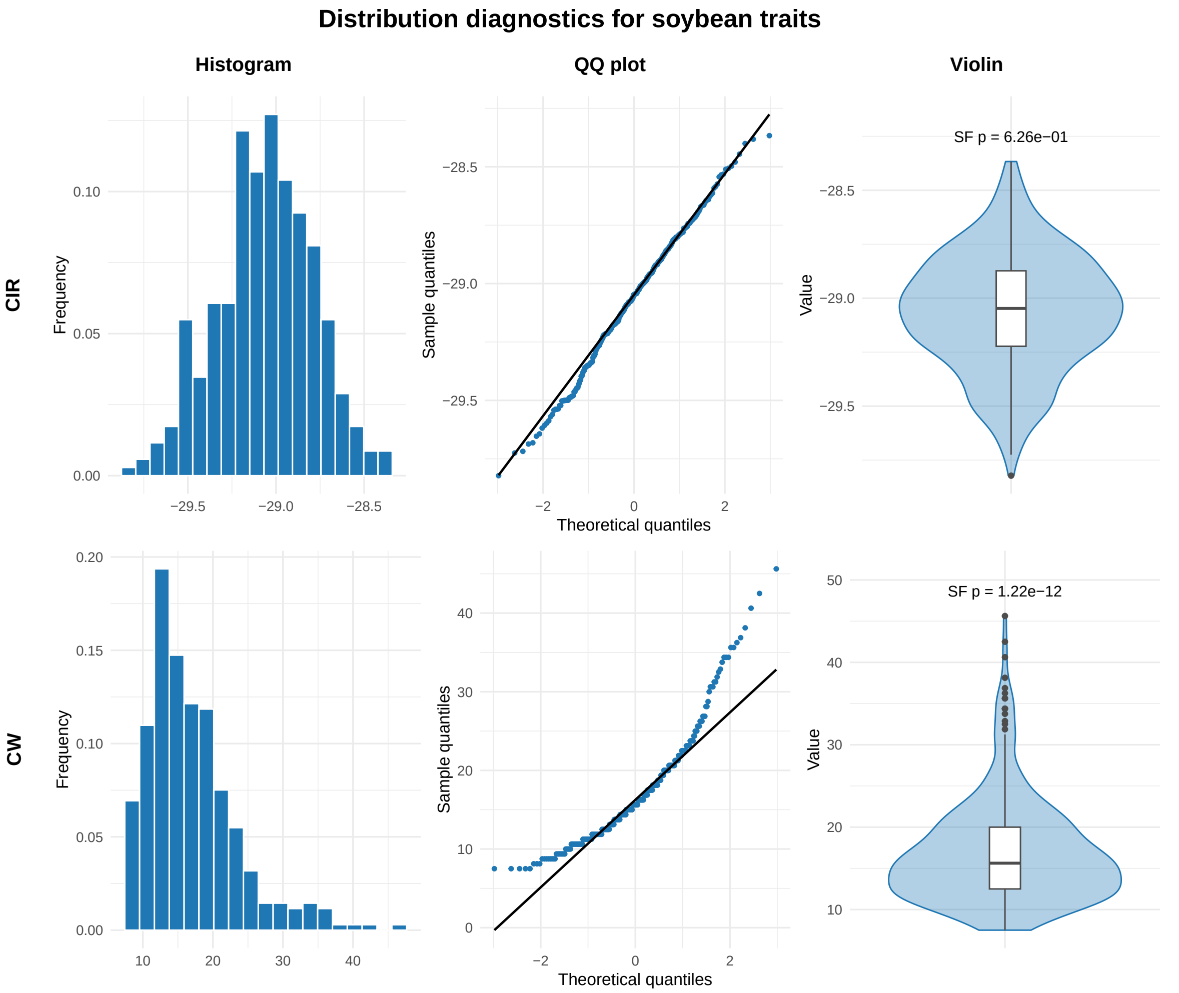


Figure S3: Distribution diagnostic plots for the soybean traits (*carbon isotope ratio* - CIR & *canopy wilting* - CW). Shapiro–Francia normality test p-values are shown alongside the violin plots.

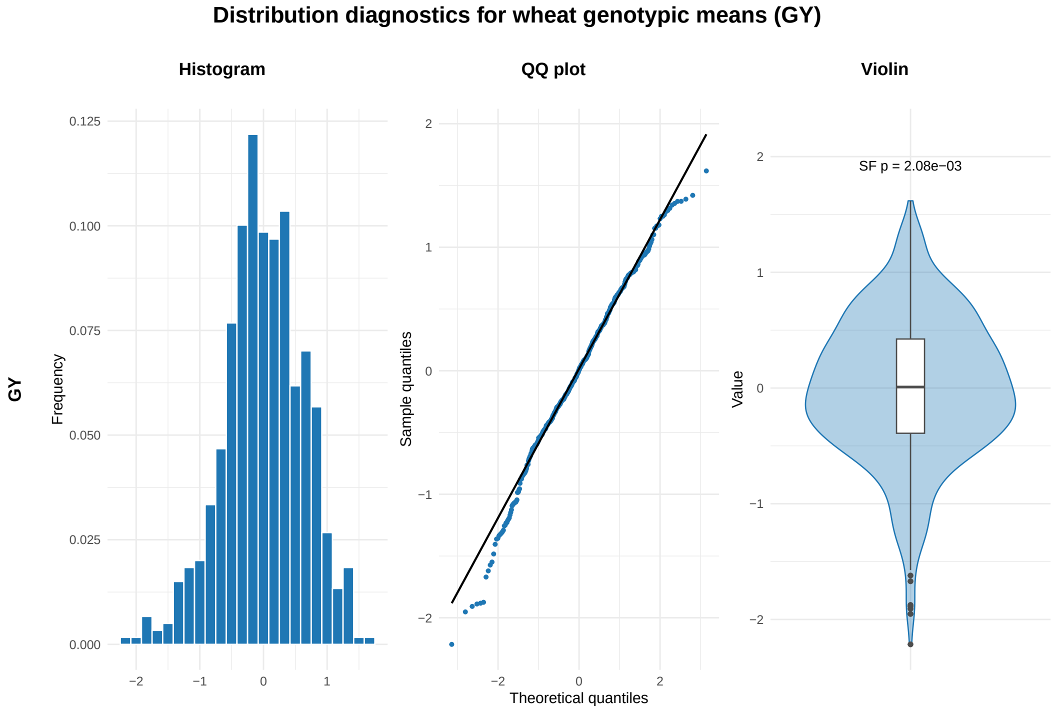


Figure S4: Distribution diagnostic plots for the wheat *grain yield* classical (GY) genotypic means. Shapiro–Francia normality test p-values are shown alongside the violin plots.

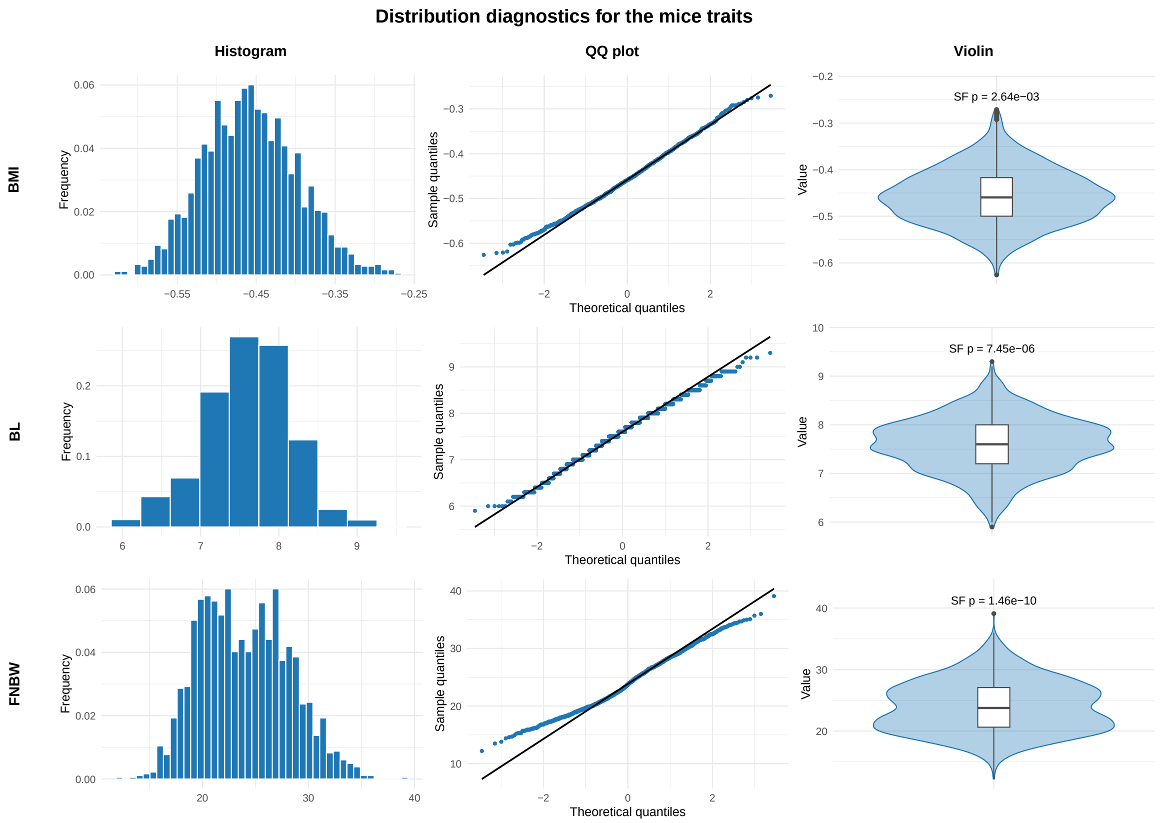


Figure S5: Distribution diagnostic plots for the three obesity mice traits (Trait 1: body mass index; Trait 2: body length & Trait 3: final normalized body weight). Shapiro–Francia normality test p-values are shown alongside the violin plots.

(a)

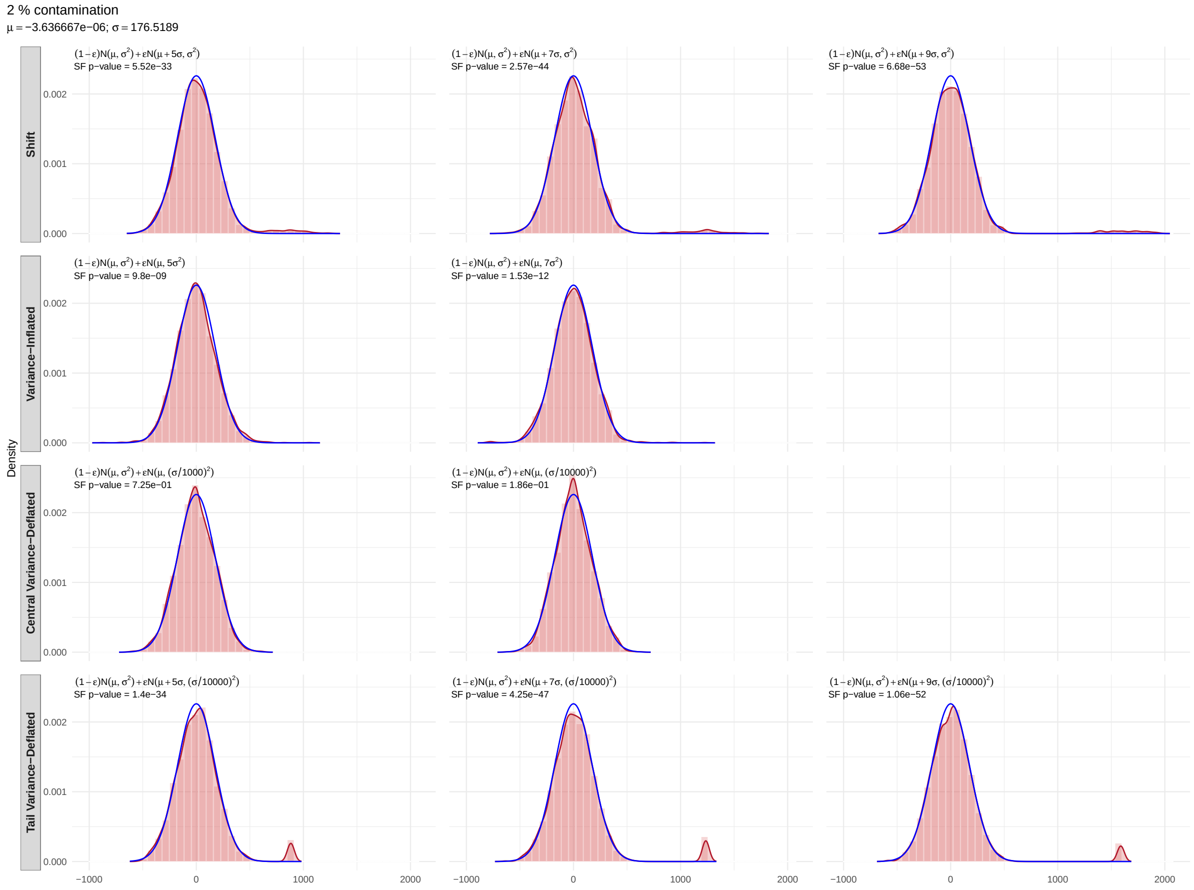


Figure S6: Distribution of the contaminated data under the three contamination scenarios and across contamination percentages: (a) 2% contamination; (b) 5% contamination; (c) 10% contamination.

(b)

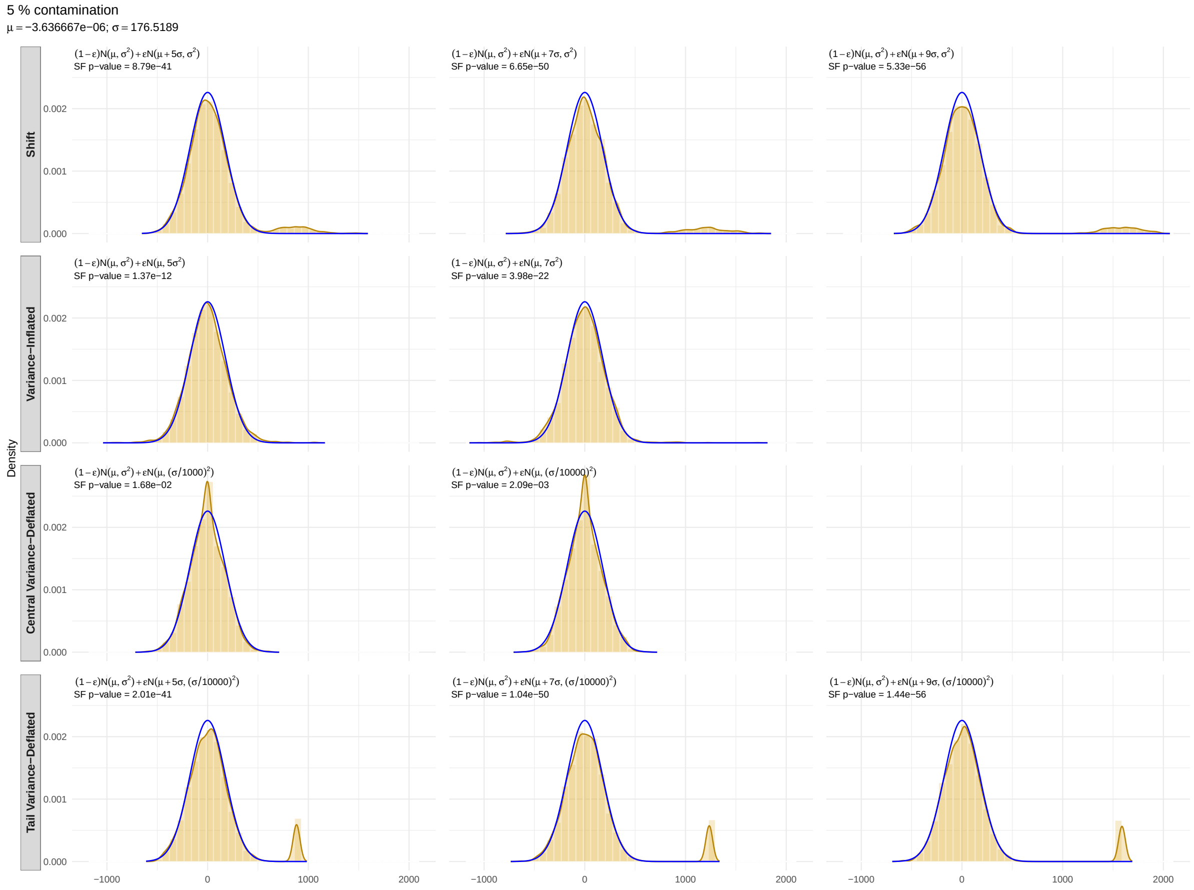


Figure S6: Distribution of the contaminated data (continued).

(c)

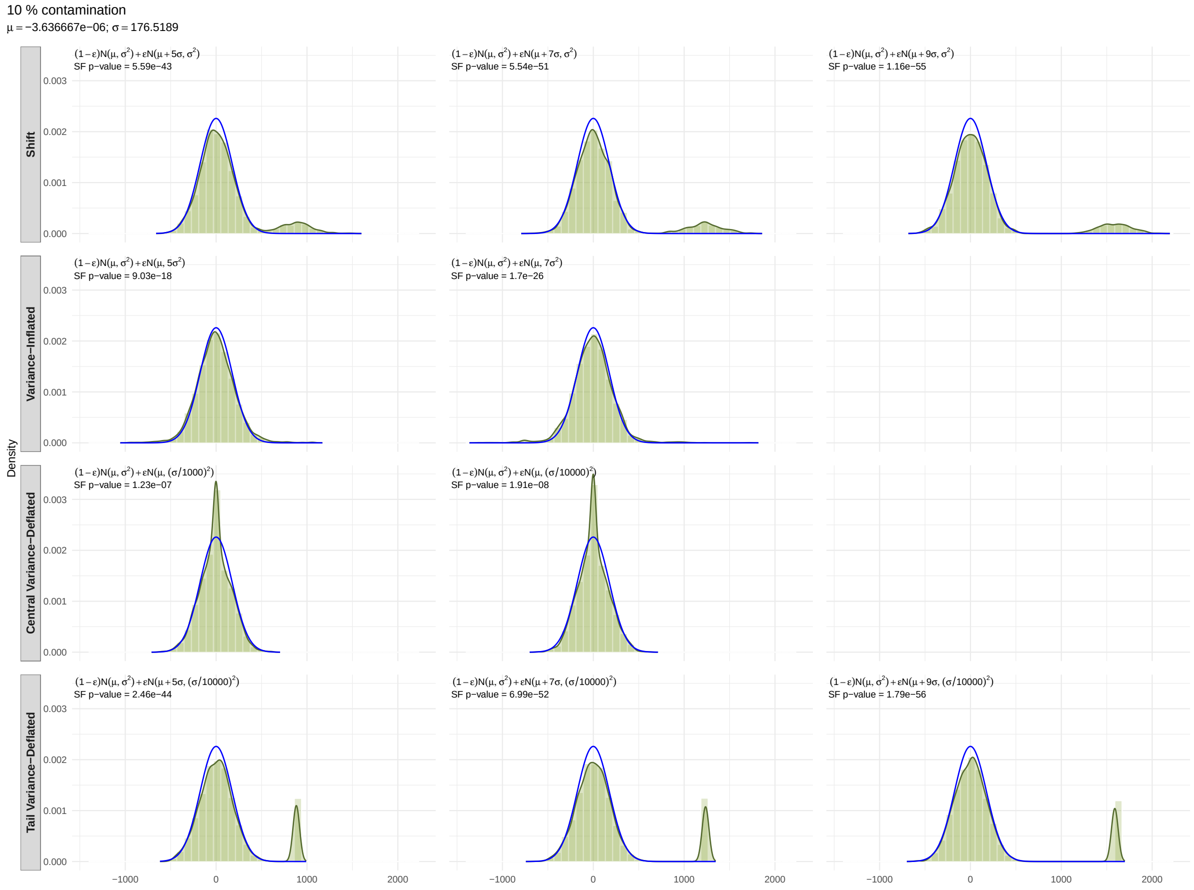


Figure S6: Distribution of the contaminated data (continued).

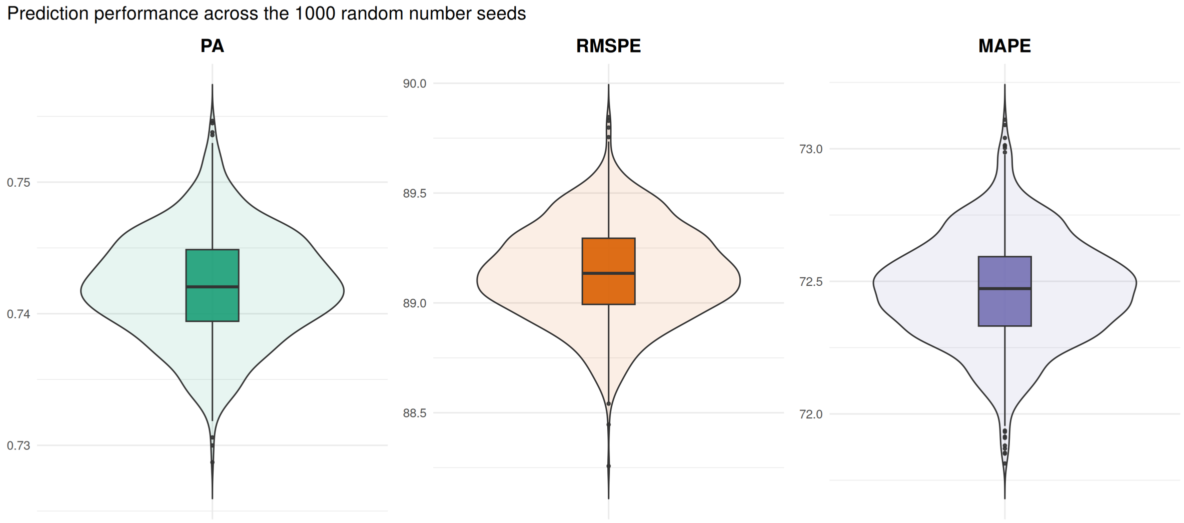


| Metric | Mean | SD | Min | Max | Range |
|---|---|---|---|---|---|
| PA | 0.742 | 0.004 | 0.729 | 0.755 | 0.026 |
| RMSPE | 89.100 | 0.200 | 88.300 | 89.800 | 1.600 |
| MAPE | 72.500 | 0.200 | 71.800 | 73.100 | 1.300 |

(a) standard Random Forests (RF).

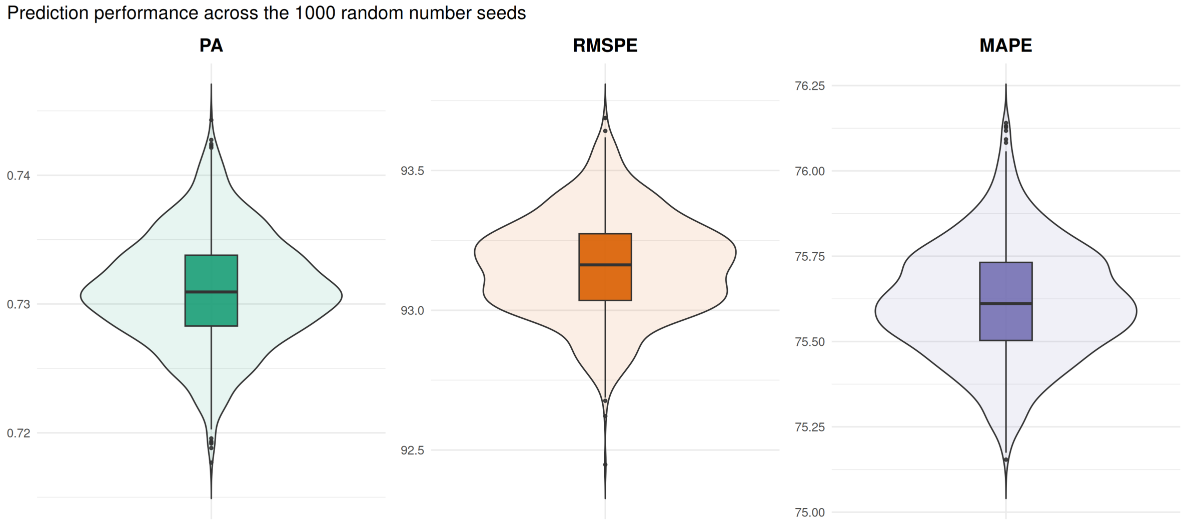


| Metric | Mean | SD | Min | Max | Range |
|---|---|---|---|---|---|
| PA | 0.731 | 0.004 | 0.718 | 0.744 | 0.027 |
| RMSPE | 93.200 | 0.200 | 92.400 | 93.700 | 1.200 |
| MAPE | 75.600 | 0.200 | 75.200 | 76.100 | 1.000 |

(b) RF with weighted observations.

Figure S7: Prediction performance (PA: predictive accuracy; RMSPE: mean squared prediction error; and MAPE: mean absolute error; SD: standard deviation) of the standard Random Forests (RF) across the 1000 random number seeds when the different preprocessing techniques (transformations) are applied to the original simulated data. Additional methodological details relevant to interpreting this figure are provided in Section *RF hyperparameter tuning and other specifics*, Subsection *Simulation data* at the beginning of the Supplementary Materials.

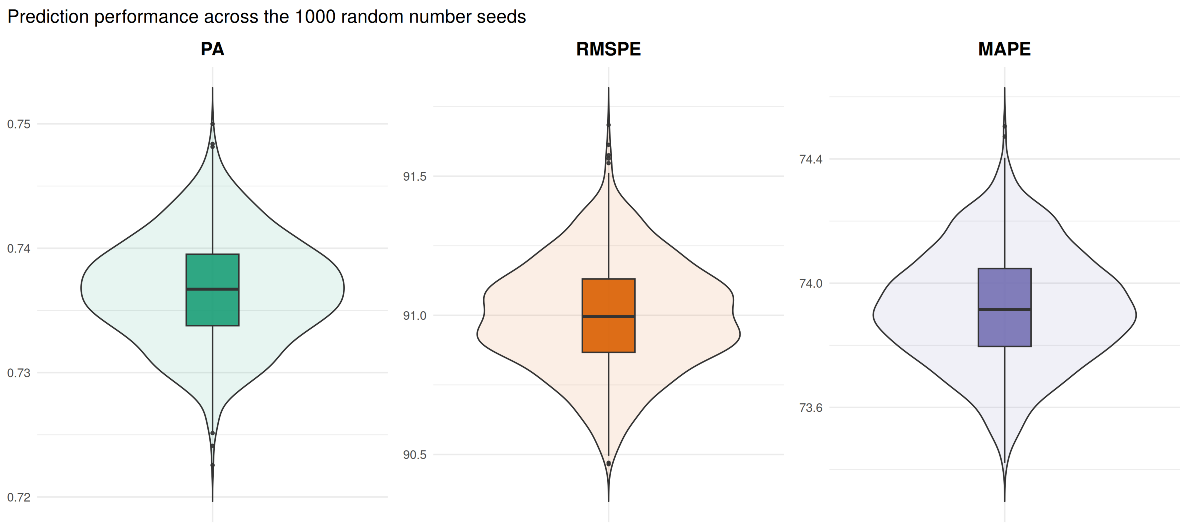


| Metric | Mean | SD | Min | Max | Range |
|---|---|---|---|---|---|
| PA | 0.737 | 0.004 | 0.723 | 0.75 | 0.027 |
| RMSPE | 91.000 | 0.200 | 90.500 | 91.70 | 1.200 |
| MAPE | 73.900 | 0.200 | 73.400 | 74.50 | 1.100 |

(c) RF with winsorised observations.

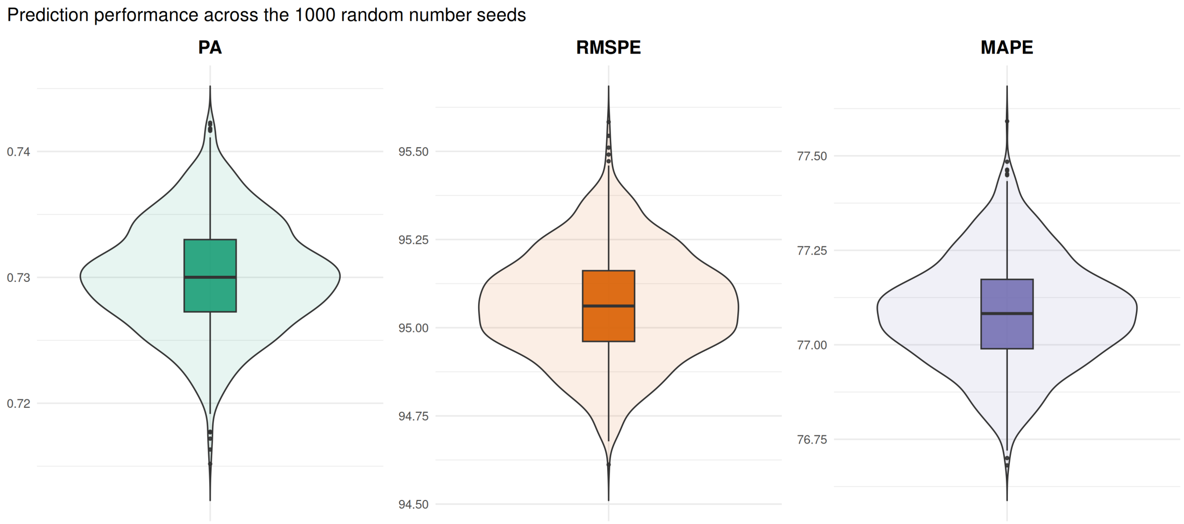


| Metric | Mean | SD | Min | Max | Range |
|---|---|---|---|---|---|
| PA | 0.73 | 0.004 | 0.715 | 0.742 | 0.027 |
| RMSPE | 95.10 | 0.200 | 94.600 | 95.600 | 1.000 |
| MAPE | 77.10 | 0.100 | 76.700 | 77.600 | 0.900 |

(d) RF with ranked observations.

Figure S7: Prediction performance (PA: predictive accuracy; RMSPE: mean squared prediction error; and MAPE: mean absolute error; SD: standard deviation) of the standard Random Forests (RF) across the 1000 random number seeds (continued).

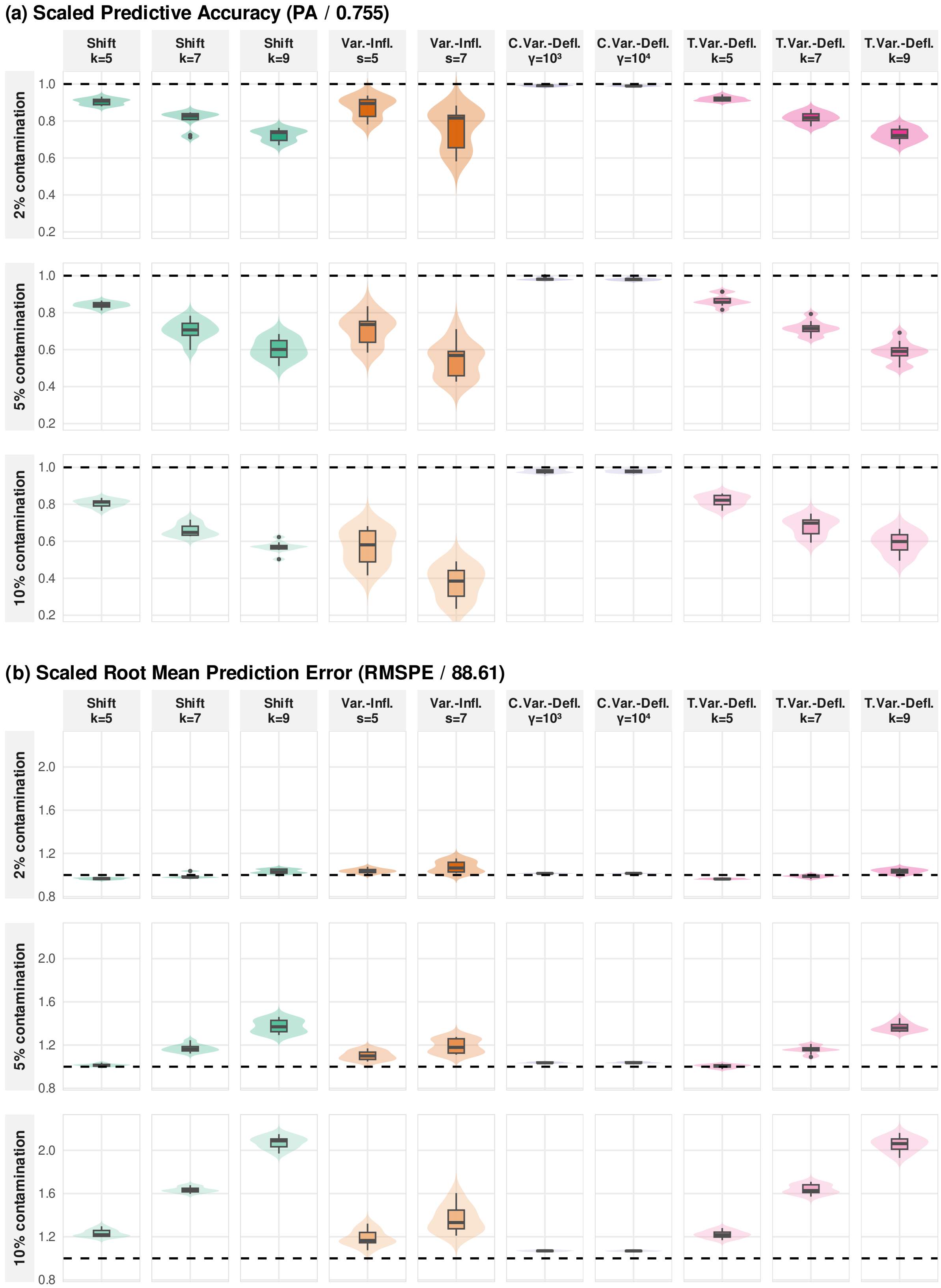


Figure S8: Estimated scaled predictive accuracy (PA), scaled root mean squared prediction error (RMSPE) and scaled mean absolute prediction error (MAPE) for the standard Random Forests (RF) model across all contamination scenarios, the three contamination levels, and the 10 simulation runs. Each simulation run corresponds to one contaminated simulated dataset.

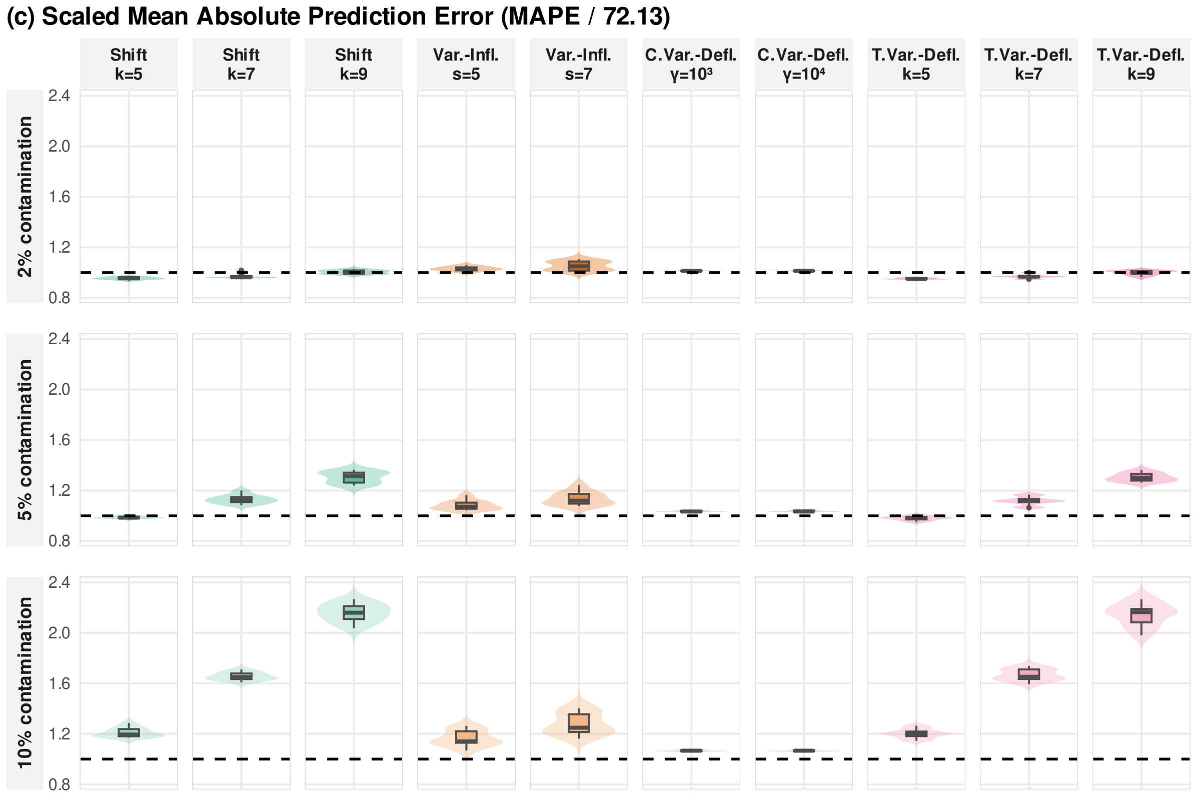


Figure S8: Estimated scaled predictive accuracy (PA), scaled root mean squared prediction error (RMSPE) and scaled mean absolute prediction error (MAPE) for the standard Random Forests (RF) model (continued).

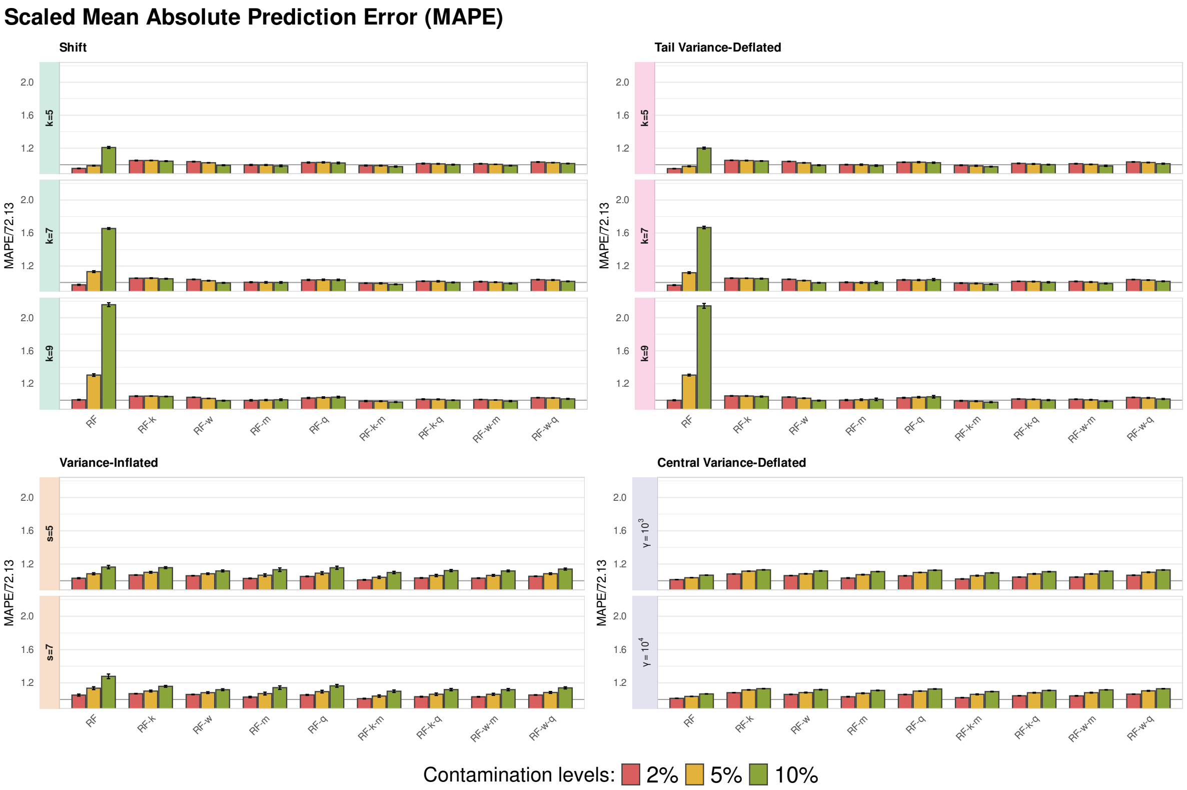


Figure S9: Estimated scaled mean absolute prediction error (MAPE) of the competing methods under the four contamination scenarios: `Shift`, `Variance-Inflated`, `Central Variance-Deflated` and `Tail Variance-Deflated`. Vertical bars indicating standard errors across simulation runs at contamination levels of 2%, 5%, and 10%. Facets correspond to the contamination parameter values ($k$, $s$, or $\gamma$, depending on the scenario).

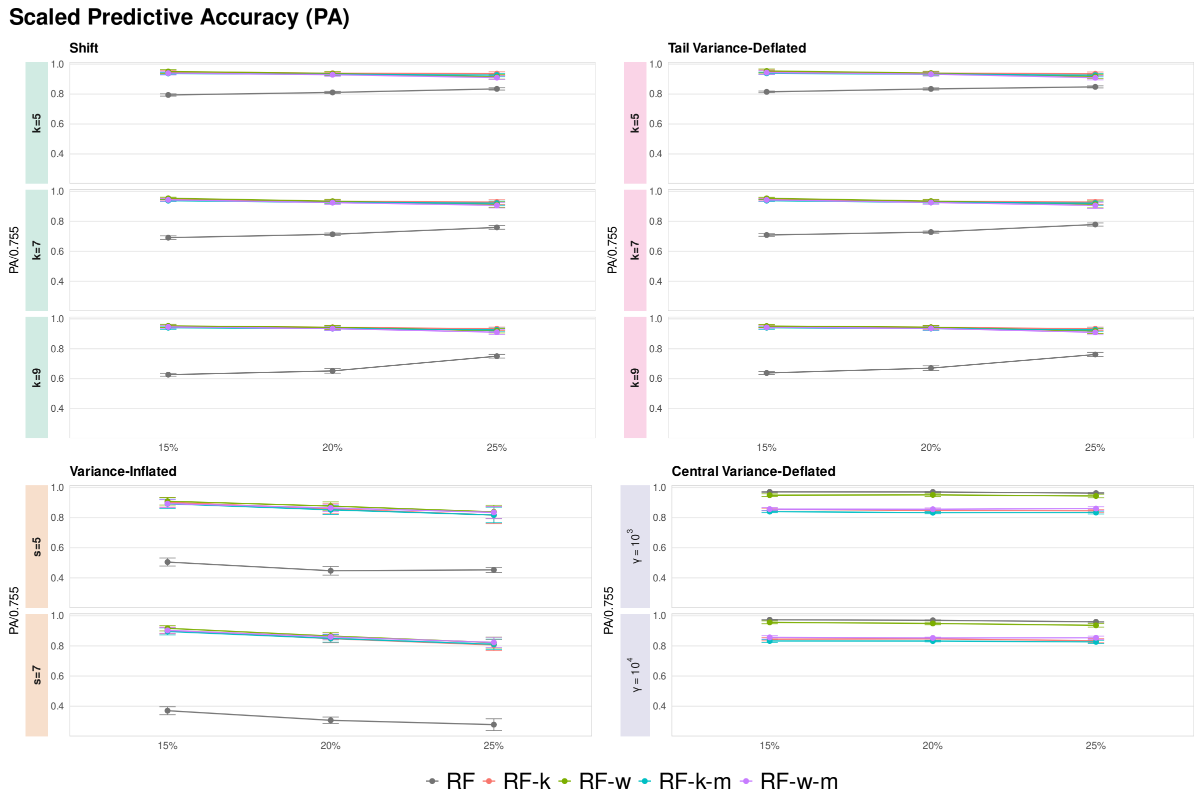


Figure S10: Estimated scaled predictive accuracy (PA) of the competing methods under the four contamination scenarios: `Shift`, `Variance-Inflated`, `Central Variance-Deflated` and `Tail Variance-Deflated`. Points represent the mean PA at contamination levels of 15%, 20%, and 25%. Facets correspond to the contamination parameter values ($k$, $s$, or $\gamma$, depending on the scenario).

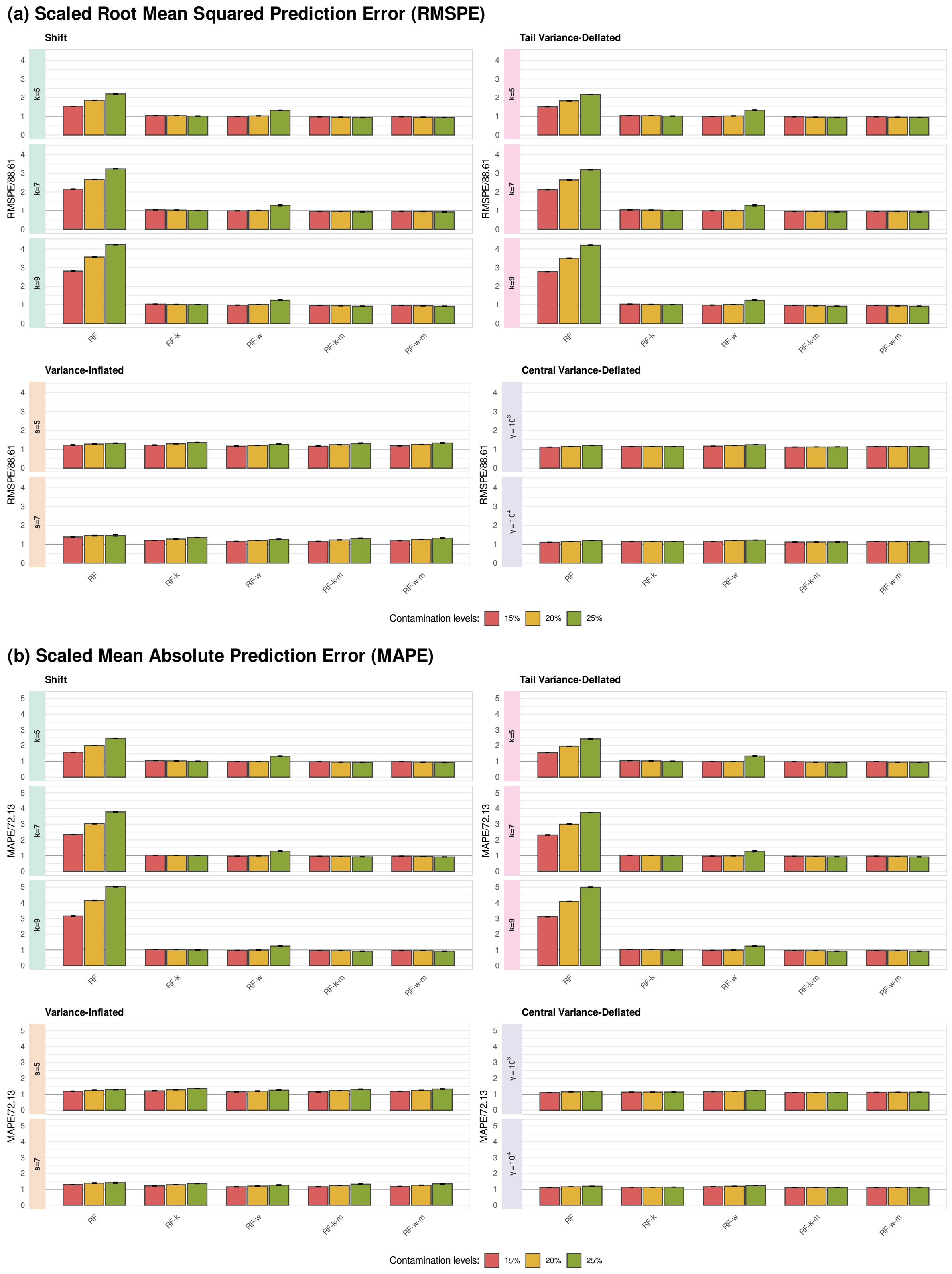


Figure S11: Estimated scaled mean squared prediction error (RMSPE) and mean absolute prediction error (MAPE) of the competing methods under the four contamination scenarios: `Shift`, `Variance-Inflated`, `Central Variance-Deflated` and `Tail Variance-Deflated`. Facets correspond to the contamination parameter values ($k$, $s$, or $\gamma$, depending on the scenario).

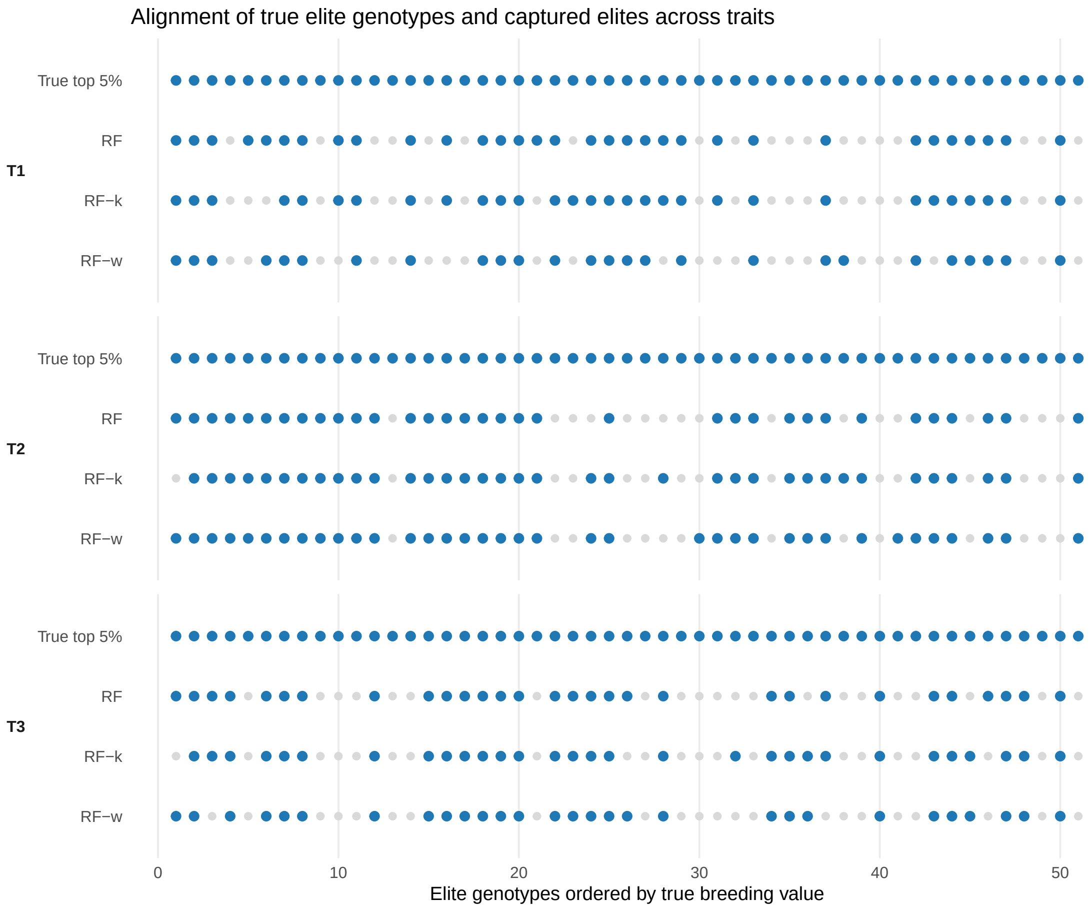


Figure S12: Alignment of captured elite genotypes across random forests (RF) methods and traits. The first row shows the true top 5% genotypes ranked by breeding value, while the rows below indicate which of these elites are recovered within the predicted top 10% by RF, RF-k, and RF-w.

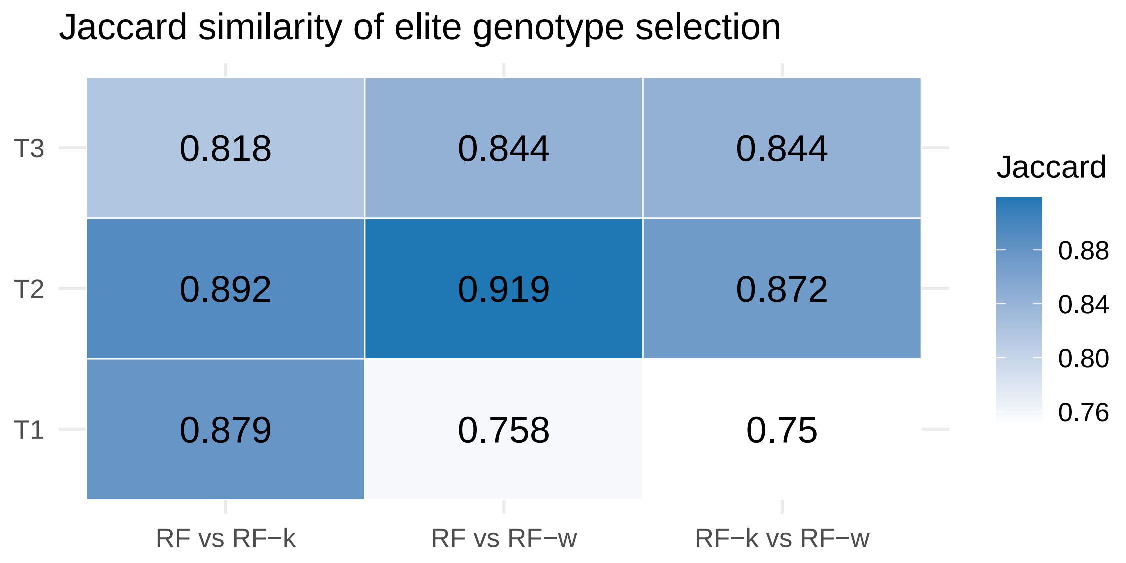


Figure S13: Jaccard similarity indices measuring the overlap between the sets of captured true top 5% genotypes identified by RF, RF-k, and RF-w for each trait. Values closer to 1 indicate stronger agreement between the methods in selecting elite individuals.

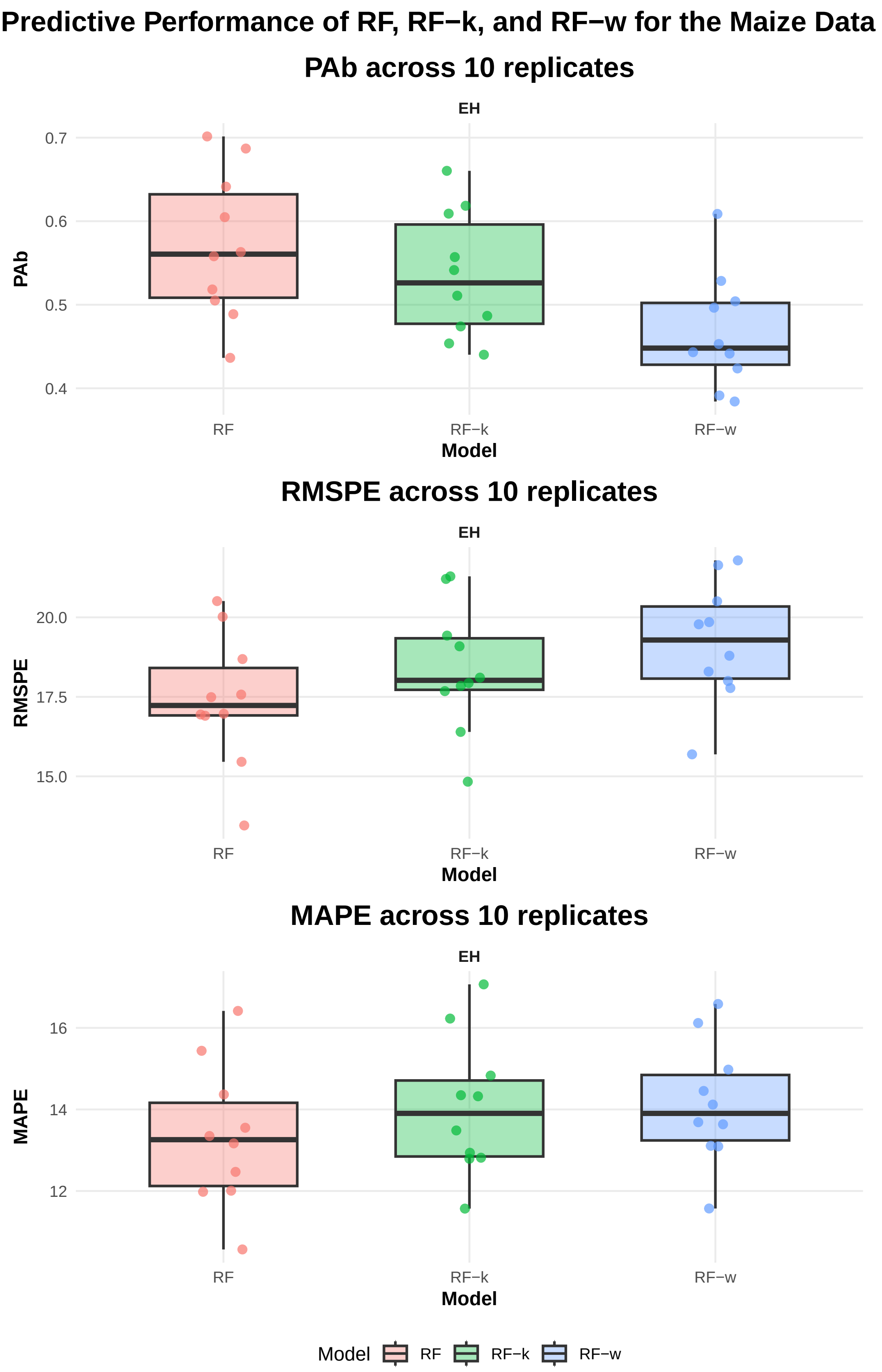


Figure S14: Predictive performance of RF, RF-k, and RF-w for the maize *ear height* (EH) trait across 10 replications. The boxplots summarize variability in predictive accuracy (PA), root mean squared prediction error (RMSPE), and mean absolute prediction error (MAPE). Points correspond to individual replicate results.

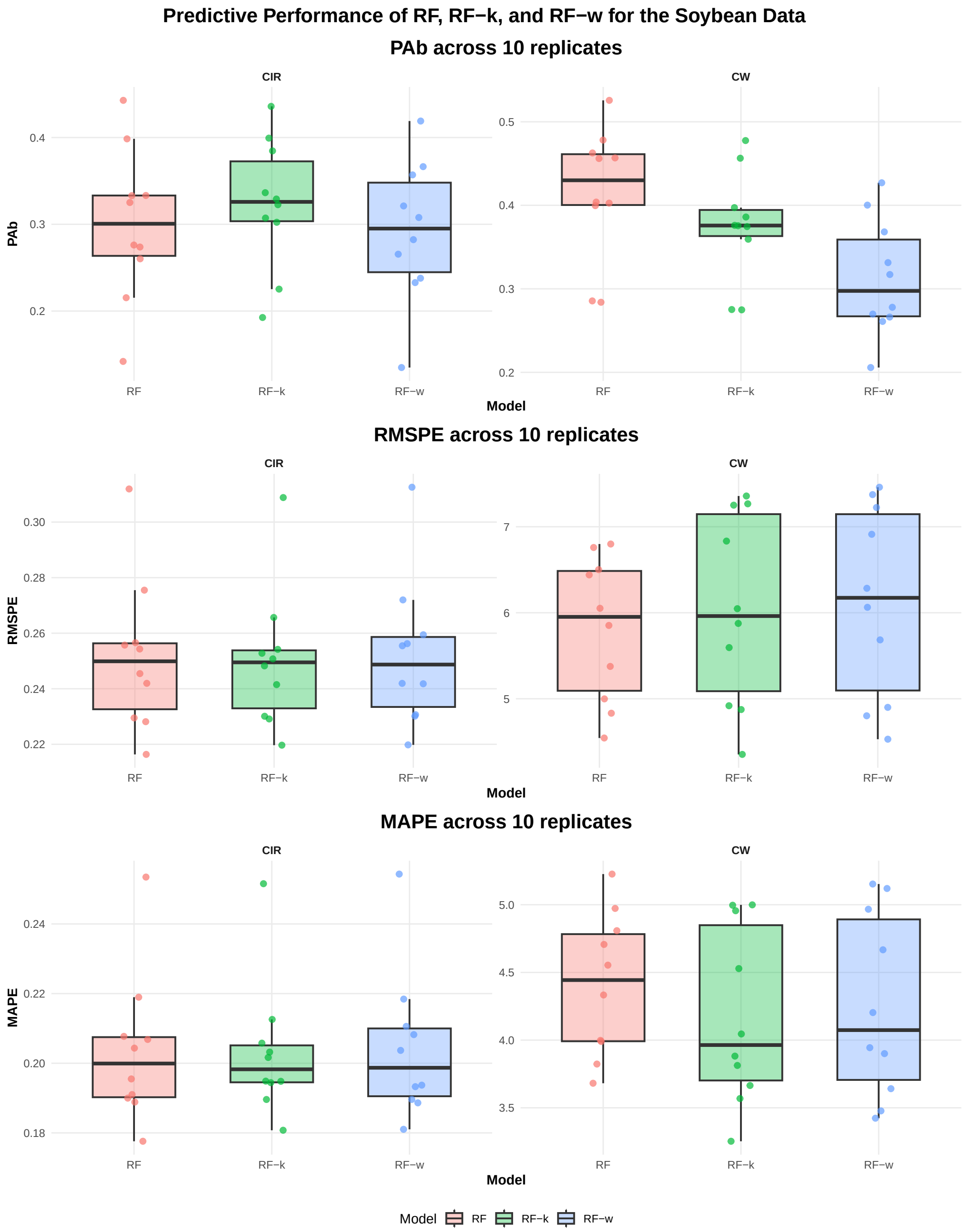


Figure S15: Predictive performance of RF, RF-`k`, and RF-`w` for the soybean *carbon isotope ratio* (CIR) and *canopy wilting* (CW) across 10 replications. The boxplots summarize variability in predictive accuracy (PA), root mean squared prediction error (RMSPE), and mean absolute prediction error (MAPE). Points correspond to individual replicate results.

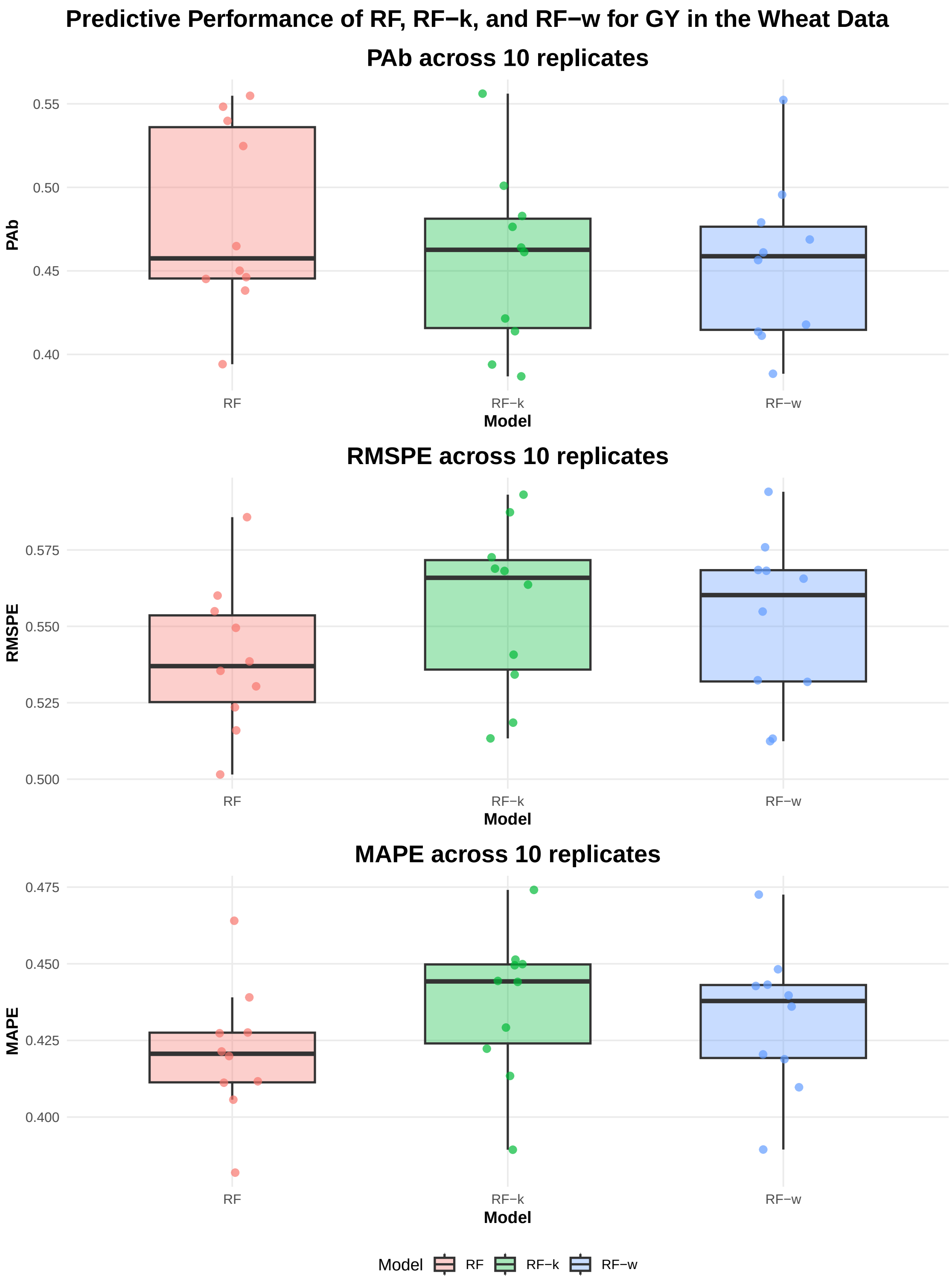


Figure S16: Predictive performance of RF, RF-k, and RF-w for the wheat *grain yield* genotypic means (GY) across 10 replications. The boxplots summarize variability in predictive accuracy (PA), root mean squared prediction error (RMSPE), and mean absolute prediction error (MAPE). Points correspond to individual replicate results.

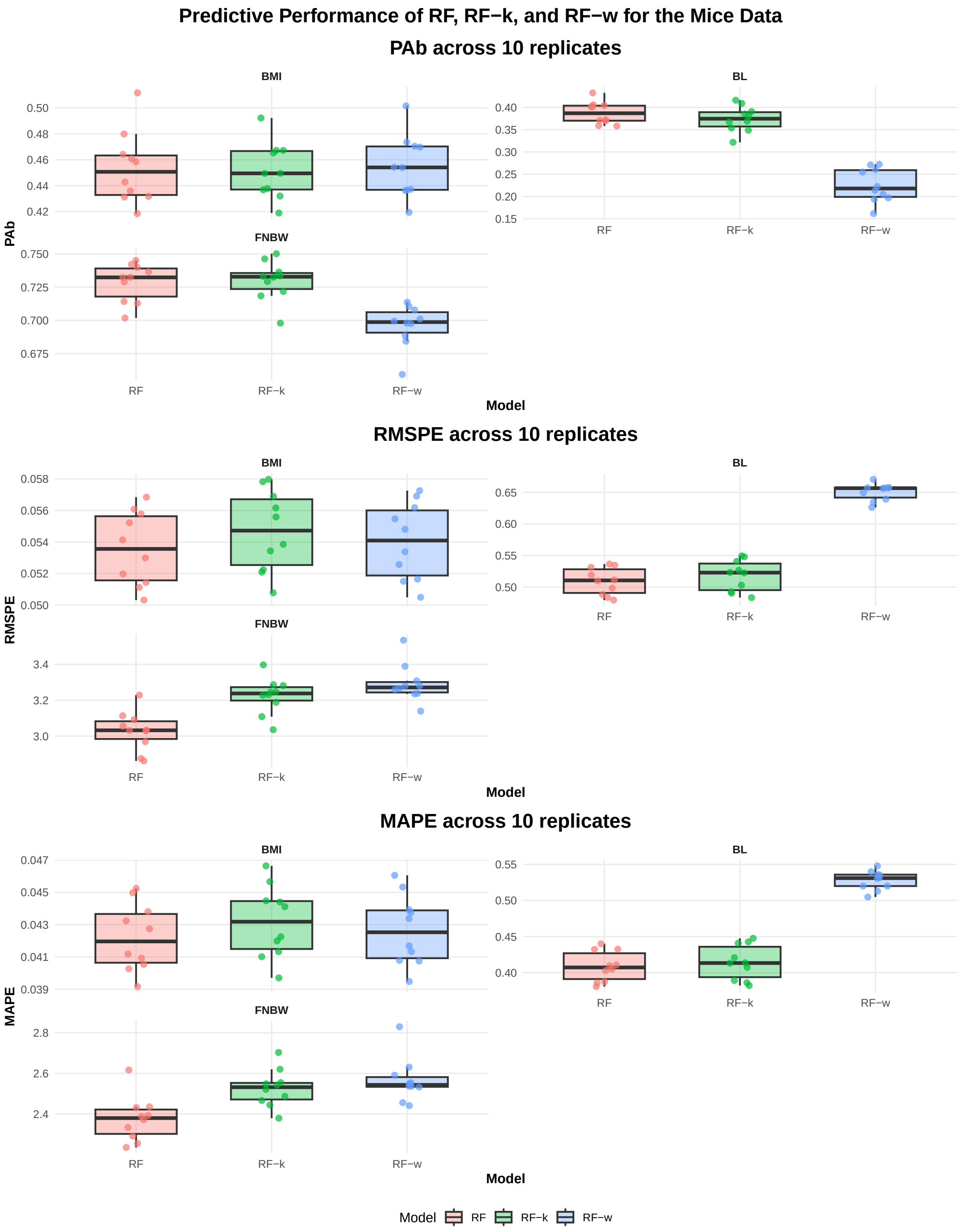


Figure S17: Predictive performance of RF, RF-k, and RF-w for the mice traits, *body mass index* (BMI), *body length* (BL) and *final normalized body weight* (FNBW) across 10 replications. The boxplots summarize seed-to-seed variability in predictive accuracy (PA), root mean squared prediction error (RMSPE), and mean absolute prediction error (MAPE). Points correspond to individual seed results.

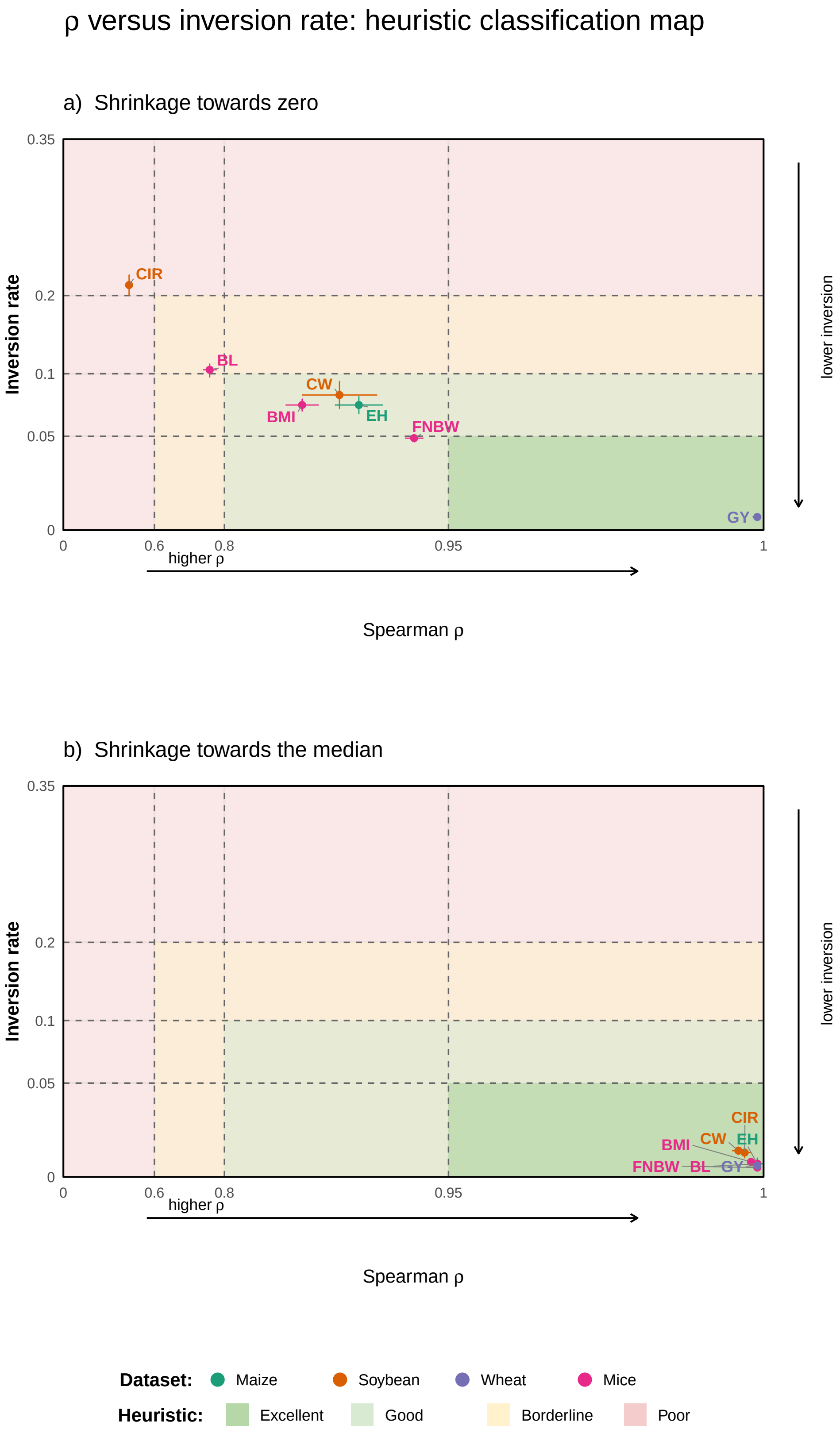


Figure S18: Relationship between Spearman rank correlation ($\rho$) and inversion rate used as a heuristic diagnostic for ranking preservation under weighting. The coloured regions define qualitative regimes (Excellent, Good, Borderline, Poor). Each point represents a dataset–trait combination, with error bars showing variability across splits. Combinations located in the bottom-right region (high $\rho$, low inversion) indicate strong agreement with the original ranking.
